# ArpC5 isoforms regulate Arp2/3 complex-dependent protrusion through differential Ena/VASP positioning

**DOI:** 10.1101/2022.07.28.501813

**Authors:** Florian Fäßler, Manjunath G Javoor, Julia Datler, Hermann Döring, Florian W Hofer, Georgi Dimchev, Victor-Valentin Hodirnau, Klemens Rottner, Florian KM Schur

## Abstract

Tight regulation of Arp2/3 complex is required to allow productive nucleation of force-generating, branched actin networks. An emerging aspect of regulation is the incorporation of subunit isoforms into Arp2/3 complex. Specifically, both isoforms of the ArpC5 subunit, ArpC5 and ArpC5L, have been reported to fine-tune nucleation activity and branch junction stability. Elevated levels of ArpC5 have also been linked to increased cancer progression and metastasis. Here, we have combined genetic engineering of cells and cellular structural biology to describe how ArpC5 and ArpC5L differentially regulate cell migration. They do so by defining the structural stability of ArpC1 in branch junctions and, in turn, by determining protrusion characteristics, protein dynamics, and actin network ultrastructure. ArpC5 isoforms also have an impact on the positioning of actin assembly factors from the Ena/VASP family, which act downstream of Arp2/3 complex-mediated nucleation. This suggests that ArpC5 and Ena/VASP proteins, both predictors for poor outcome in cancer, are part of a signaling pathway enhancing cell migration and, by inference, metastasis.

## Introduction

The actin filament nucleating Arp2/3 complex is involved in essential cellular processes such as endocytosis, cell shape control, DNA repair, and cell migration (*1–3*). The Arp2/3 complex also plays important roles in pathological conditions, such as in the actin cytoskeleton-dependent motility of selected viruses and bacteria or cancer metastasis (*4–7*).

The Arp2/3 complex, consisting of the two actin-related proteins (Arp) 2 and Arp3 and five scaffolding subunits ArpC1-5 (Figure 1A), generates branched actin networks via nucleating new daughter actin filaments on the sides of pre-existing mother filaments (*2, 8–11*). This leads to the formation of so-called actin filament Arp2/3 complex branch junctions showing a defined geometry of ∼70° (*10, 12–16*). As the Arp2/3 complex is able to nucleate its own substrate, branched actin networks are potentially capable of exponential growth. This requires Arp2/3 complex activity to be tightly regulated to determine actin network turnover and stability, with regulation occurring both upstream and downstream of Arp2/3 complex-mediated branch junction formation (*5*). Specifically, Arp2/3 complex activation is driven by nucleation-promoting factors (NPFs), such as members of the Wiskott-Aldrich syndrome protein (WASP) family. In addition, Cortactin binds to branch junctions to increase their stability and might enhance the release of NPFs (*17*). Inhibitors of Arp2/3 complex-dependent actin assembly include Arpin (*18*), Coronin (*19*) or glia maturation factor (GMF) (*20*), all of which act at different stages of the Arp2/3 complex activity cycle, from interfering with NPF-activation to branch junction disassembly.

**Figure 1:**
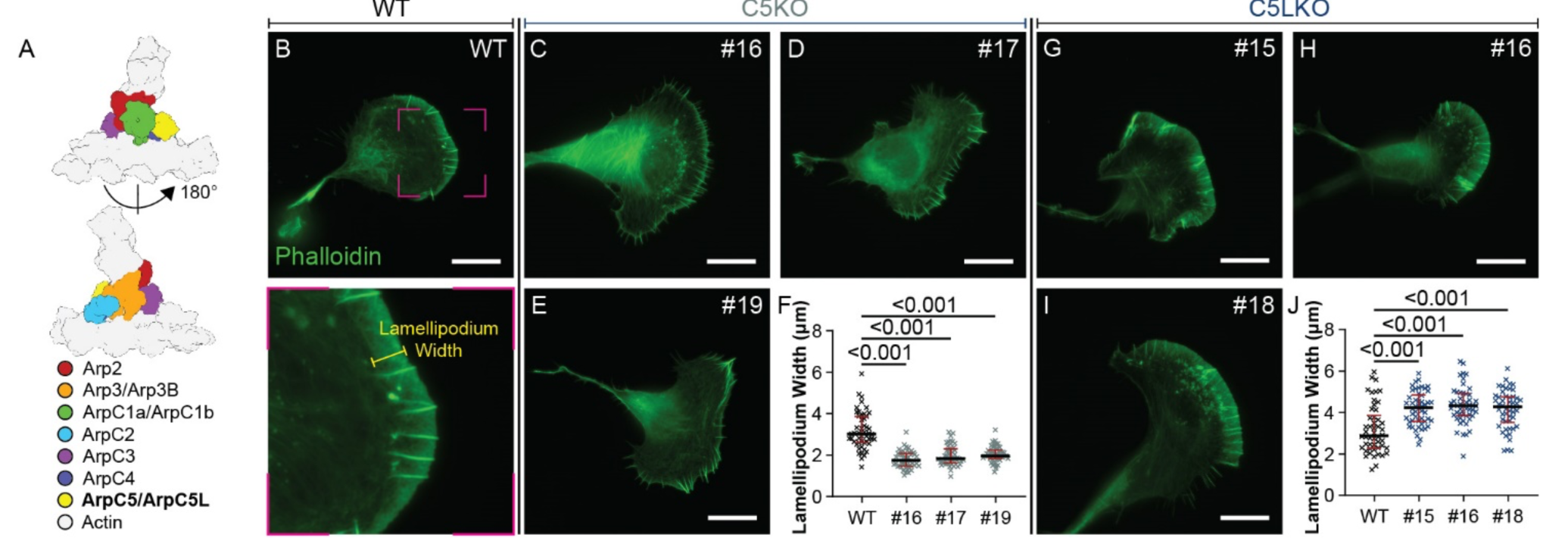
ArpC5 isoforms affect lamellipodium morphology. **(A)** Schematic representation of the branch junction highlighting the positions of the individual Arp2/3 complex subunits within this assembly. Subunit colors are annotated. **(B-J)** Analysis of lamellipodium width phenotype. (B-E, G-I) Representative epifluorescence micrographs of B16-F1 wildtype (B), C5KO (C-E), and C5LKO (G-I) cells visualizing the actin cytoskeleton using fluorescent phalloidin. For the knockout cells, three individual clones are shown. The lower panel in (B) shows a zoom-in of the indicated area in the upper panel and exemplifies which areas were considered as lamellipodia for width measurements. Three such measurements were performed per cell, and their average was then employed for statistical analysis. (F) Lamellipodium width of B16-F1 C5KO lines and B16-F1 wildtype cells. Kruskal-Wallis test combined with Dunn’s multiple comparison test, n=50 cells for each experimental group, p values shown in the chart. Red lines indicate medians (3.020μm, 1.742μm, 1.830μm, 1.972μm), and black lines indicate quartile ranges. (J) Lamellipodium width of B16-F1 C5LKO lines and B16-F1 wildtype cells. Kruskal-Wallis test combined with Dunn’s multiple comparison test, n=50 cells for each experimental group, p values shown in the chart. Red lines indicate medians (2.884μm, 4.237μm, 4.324μm, 4.280μm), and black lines indicate quartile ranges. All scale bars, 20µm

Arp2/3 complex nucleation activity is also determined by post-translational modifications or isoform specificity of Arp2/3 complex subunits. The Arp2/3 complex is not a uniform macromolecular entity but can exist in eight different subunit compositions, caused by the presence of Arp3a/Arp3b, ArpC1a/ArpC1b, and ArpC5/ArpC5L subunit variants (commonly referred to as isoforms in the literature). This compositional heterogeneity of the Arp2/3 complex allows for a fine-tuned nucleation activity and branch junction stability (*21–23*). Importantly, certain ArpC5-ArpC1-isoform combinations seem to be preferentially assembled into one complex (*21*). Together, these mechanisms define the overall activity of the complex in *in vitro* reconstituted actin networks or in Vaccinia Virus (VACV) actin tails in infected cells, with ArpC1a and ArpC5 being associated with lower Arp2/3 activity compared to the combination of ArpC1b and ArpC5L (*24*). Specifically, Arp2/3 complex isoform-specific activity was described to depend on the preferred stabilization of ArpC5L-containing branches by Cortactin against Coronin-induced debranching (Abella et al. 2016). However, it can be anticipated that additional regulatory layers beyond Cortactin stabilization of isoform-specific branch junctions are at play to modulate Arp2/3 complex activity.

Single-particle cryo-EM of ArpC1a/ArpC5 and ArpC1b/ArpC5L-containing recombinant Arp2/3 complexes showed slightly different stability of the ArpC5 isoforms in the inactive complex but did not otherwise reveal substantial structural differences in the different Arp2/3 complex variants. This suggests that their specific activities either result from different energetic barriers for activation (*2*) or that major structural differences, which cause changes in nucleation activity, are only prevalent in the active conformation of the Arp2/3 complex within the branch junction. In a more recent study, the isoform-specific regulation of the Arp2/3 complex was extended to Arp3 by showing that the methionine monooxygenase MICAL2 enhances branched network disassembly by oxidizing Arp3b-containing Arp2/3 complexes (*22*). These exciting findings warrant further studies to identify isoform-specific Arp2/3 complex interactions with proteins upstream or downstream of branch junction nucleation.

Moreover, a better understanding is required of how isoform-specific properties of the Arp2/3 complex might differ among biological systems. Intriguingly, the ArpC5L isoform has been shown to increase Arp2/3 complex activity (Abella et al. 2016), a trait normally associated with enhanced cell migration and metastasis (*25–28*). In contrast, elevated ArpC5 levels have been linked to cancer progression (Kinoshita et al. 2012; Xiong and Luo 2018; Huang et al. 2021), as ArpC5 depletion reduces motility and invasiveness of carcinoma cells (Kinoshita et al. 2012). This raises the question if and how ArpC5/5L isoforms uniquely affect actin networks involved in cell migration and thus metastasis.

An established model to study the regulation of branched actin networks in cell motility is the lamellipodium, a thin sheet-like protrusion at the leading edge of migrating cells, densely filled with branch junctions (*29–31*). The prototypical initiation of lamellipodia depends on the activation of the Arp2/3 complex via the NPF WAVE regulatory complex (WRC). Polymerization at free barbed ends of Arp2/3 complex-nucleated filaments is potentially enhanced by the activity of actin filament elongating proteins of the Ena/VASP family (Breitsprecher et al., 2008; Damiano-Guercio et al., 2020; Faix and Rottner, 2022). However, it should also be noted that Ena/VASP proteins can potentially counteract lamellipodial Arp2/3 complex accumulation, as implied by genetic Ena/VASP disruption (*32*). Due to their morphological simplicity and their accessibility to a large variety of experimental approaches, lamellipodia have been used as model systems across different disciplines, from cell biology to structural biology. Genetic engineering of migratory cell lines via CRISPR/Cas9 has been successfully used to delineate lamellipodium formation and to understand the activity of several important actin-binding proteins (ABPs), such as WAVE and its isoforms (*33*), Lamellipodin (*34*) or FMNL subfamily formins (*35*) and the capping protein regulator Twinfilin (*36*). The lamellipodium has also been used extensively to study the ultrastructure and topology of actin networks under varying experimental conditions using electron microscopy approaches (*16, 29, 30, 37–40*). We have recently been able to exploit the high number of branch junctions in lamellipodia in a cellular structural biology approach using cryo-electron tomography (cryo-ET) to determine the structure of the actin filament Arp2/3 complex branch junction to subnanometer resolution (*13*). This allowed for a more precise understanding of the structural transitions that occur upon Arp2/3 complex activation and interactions that are formed with actin mother and daughter filament. The structural description of the branch junction has been recently extended to a 3.9 Å resolution single-particle cryo-EM structure of *in vitro* reconstituted actin filament Arp2/3 complex branch junctions (*12*). However, as the branches observed in our previous *in situ* study or in the aforementioned *in vitro* work both contained all possible Arp2/3 complex isoform compositions, no insights into isoform-specific branch junction structures could be obtained.

To explore ArpC5 isoform-dependent actin network organization and related emergent cell migration characteristics, which are expected to play different roles in cancer progression, we chose a multiscale approach combining recent developments in genetic engineering of cells and cellular structural biology. This offered the opportunity to integrate high-resolution information on isoform-specific branch junction structures and the resulting ultrastructural cytoskeleton architectures with Arp2/3 complex isoform-specific cellular morphology, actin network dynamics, and ABP-localization. To this end, we have generated KO cell lines lacking either one or both of the ArpC5/5L isoforms and describe across multiple experimental scales how the two distinct ArpC5 isoforms differentially regulate branched actin networks in cell migration. We reveal that incorporating a specific ArpC5 isoform into the Arp2/3 complex determines lamellipodial protrusion characteristics, protein dynamics, and actin network ultrastructure. While cells exclusively containing ArpC5L exhibited narrower and less protrusive lamellipodia, ArpC5-containing cells displayed even wider lamellipodia than wildtype (WT) cells. In explaining this phenotype, our data suggest that the Arp2/3 complex is not merely a nucleator of actin filaments but also influences actin elongation characteristics in an ArpC5/5L isoform-dependent fashion, e.g., by defining the positioning of actin filament elongation proteins of the Ena/VASP family at the leading edge. Our results provide novel insights into why specifically ArpC5 over ArpC5L is associated with enhanced cell motility and therefore linked with increased cancer progression and metastasis (*41–43*).

## Results

### ArpC5 isoform composition determines protrusion morphology

To explore the role of the two ArpC5 isoforms in lamellipodia architecture and cell migration, we used a CRISPR/Cas9 knockout (KO) approach (*44*) to generate B16-F1 mouse melanoma cells lacking either ArpC5 (referred to as C5KO), ArpC5L (C5LKO) or both isoforms (C5/C5LKO). For all generated cell lines, successful KO was verified in three independent clones via genomic sequencing (Figure S1) and confirmation of complete depletion of ArpC5 and/or ArpC5L protein levels by Western blotting (Figure S2A).

Double KO cell lines lacking both ArpC5 and ArpC5L isoforms were completely devoid of lamellipodia (Figure S3). These results indicate that the actin assembly activity of Arp2/3 complexes lacking an ArpC5 subunit reported *in vitro* (*45*) is insufficient to maintain branched actin networks in lamellipodia. Single isoform KO cells revealed opposing lamellipodial phenotypes (Figure 1B-J). While C5KO cells exhibited significantly narrower lamellipodia than B16-F1 wildtype (WT) cells (Figure 1F), the lamellipodial width of C5LKOs was significantly increased (Figure 1J). To corroborate our observations of these isoform-specific lamellipodia phenotypes, we generated ArpC5 and ArpC5L KOs in the Rat2 fibroblast cell line (Figures S1, S2B). Loss of either isoform in Rat2 cells resulted in lamellipodial phenotypes identical to B16-F1 cells (Figure S4), demonstrating that observed effects are consistent across at least two different species and cell types. As all lines of a given KO showed consistent phenotypes, we continued our experiments with one selected individual clone for each condition (for B16-F1 C5KO #17, 5L KO #16, C5/C5LKO #20, and for Rat2 C5KO #4 and C5LKO #11).

We transiently transfected cells with dual expression plasmids encoding either RFP alone, RFP and ArpC5L, or RFP and ArpC5 and quantified the lamellipodium width of transfected cells as identified by RFP expression. In this setup, only the construct expressing ArpC5 was able to rescue the lamellipodium phenotype of C5KO cells, resulting in wildtype-like cells (Figure 2C-D). In contrast, transfection with the ArpC5L-encoding construct did not rescue lamellipodia width (Figure 2B, D). Repeating the same transfection approach in the C5/C5LKO background (Figure 2E-H) confirmed these observations and also the functionality of our ArpC5 and ArpC5L encoding vectors, since the expression of ArpC5L or ArpC5 in double KO cells again resulted in the establishment of narrow or wide lamellipodia, respectively (Figure 2H). Considering that a reduction of ArpC5 protein levels via knockdown had been shown to decrease the abundance of ArpC1a in HeLa cells (*21*), we tested whether, in C5KO or C5LKO cells, the levels of other Arp2/3 subunits were affected as well, potentially altering overall Arp2/3 complex activity. We qualitatively assessed the relative abundance of Arp3, Arp2, ArpC1b, and ArpC1a in B16- F1 and Rat2 single KO lines (Figure S5). While we also observed a reduction in ArpC1a levels in B16-F1 C5KO and Rat2 C5KO cells, we did not see a drastic change in Arp3 or Arp2 abundance, indicating that overall Arp2/3 complex levels should not be altered as compared to the respective wildtype cell lines. As the results of the rescue experiments were consistent with ArpC5 and ArpC5L having isoform-specific functions in a lamellipodial context, we explored the putative mechanisms mediating the observed differences. Such mechanisms can include altered Arp2/3 complex protein levels (which we could partially already rule out, Figure S5), selective interactions with Arp2/3 complex regulators, ultrastructural alterations in the architecture of branched actin networks, structural differences of Arp2/3 complex branch junctions resulting in differences in branch nucleation activity, stabilization, and turnover, or – finally - differential recruitment of downstream factors.

**Figure 2:**
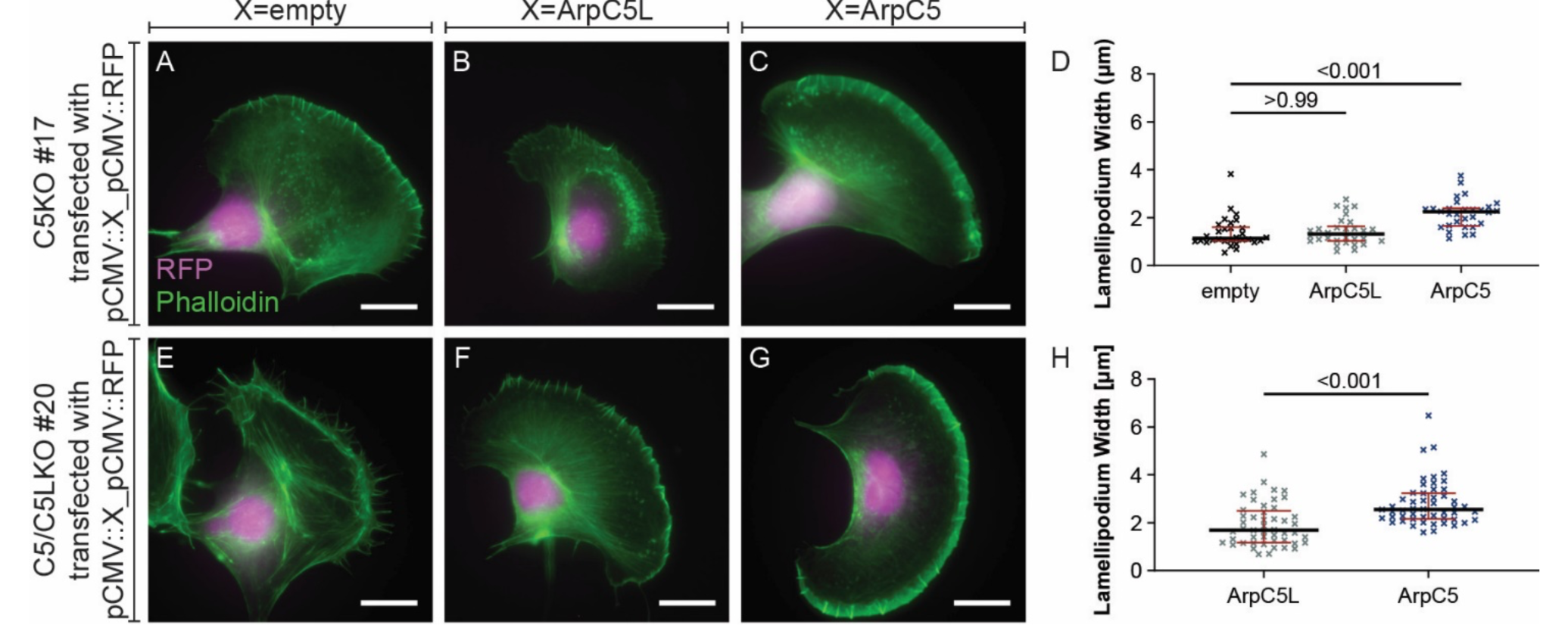
The lamellipodial C5KO phenotype is mediated by isoform specificity and not by gene dose effects. **(A-C)** Representative epifluorescence micrographs of B16-F1 C5KO cells transfected with pCMV::empty_pCMV::RFP (A), pCMV::ArpC5L_pCMV::RFP (B), pCMV::ArpC5_pCMV::RFP (C) visualizing the actin cytoskeleton using fluorescent phalloidin and verifying transfection via RFP fluorescence. **(D)** Quantitation of lamellipodium width of B16-F1 C5KO cells transfected with the same constructs as in (A-C). Measurements were done as shown in Figure 1B. Kruskal-Wallis test combined with Dunn’s multiple comparison test, n=30 cells per experimental condition, p values shown in the chart. Red lines indicate medians (1.143µm, 1.310µm, 2.248µm), and black lines indicate quartile ranges. **(E-G)** Representative epifluorescence micrographs of B16-F1 C5/C5LKO cells transfected with pCMV::empty_pCMV::RFP (E), pCMV::ArpC5L_pCMV::RFP (F), pCMV::ArpC5_pCMV::RFP (G) visualizing the actin cytoskeleton using fluorescent phalloidin and verifying transfection via RFP fluorescence. **(H)** Quantitation of lamellipodium width of B16-F1 C5/C5LKO cells transfected with the same constructs as in (F-G). No values are given for cells transfected with pCMV::empty_pCMV::RFP as those did not exhibit lamellipodia. Mann-Whitney test, n=51, 49 with p values shown in the chart. Red lines indicate medians (1.698µm, 2.549µm), and black lines indicate quartile ranges. All scale bars, 20µm

### WAVE 1 and 2 do not activate the Arp2/3 complex in an ArpC5 isoform-specific manner

We first sought to examine whether upstream regulators of the Arp2/3 complex are causative for the observed phenotypes in C5KO and C5LKO cells.The WAVE-regulatory complex (WRC) is a central nucleation promoting factor (NPF), being crucial for Arp2/3 complex activation in canonical lamellipodia (*46, 47*). Using recently published B16-F1 WAVE1 and WAVE2 KO cell lines (*33*), we aimed to test whether the two WAVE isoforms expressed in B16-F1 cells exhibit preferred activation of either ArpC5 or ArpC5L containing Arp2/3 complexes. We employed isoform- specific antibodies against ArpC5 and ArpC5L (Figure S6) to detect endogenous ArpC5 and ArpC5L in WAVE1 and WAVE2 KO cells (Figure S7). ArpC5 and ArpC5L were localized identically at the leading edge in cells lacking either WAVE1 or 2 (Figure S7D-I). This shows that Arp2/3 complexes with different ArpC5 isoform compositions are not selectively activated by WAVE1 *versus* WAVE2-containing WRCs. In a WAVE1/2 double KO cell line, neither ArpC5 nor ArpC5L were detected at the cell periphery (Figure S7J-L), in line with the WRC’s central role in establishing Arp2/3-dependent protrusions.

### Cortactin activity does not impact lamellipodia dimension in an ArpC5 isoform-specific fashion

Having excluded differential Arp2/3 complex activation by the WRC as a central contributor to ArpC5 isoform specificity in lamellipodia, we focused on the well-established branch junction stabilizer Cortactin (*48, 49*). Cortactin has been described to protect ArpC5L containing branch junctions from Coronin-mediated debranching (*21*).

We performed Cortactin knockdowns in B16-F1 WT, C5KO, and C5LKO cells (Figure S8). We confirmed the reduction in Cortactin levels at the single cell level by immunostaining (Figure S8B, D, F, H, J, L), as well as at the population level by Western blotting (Figure S8M). We then evaluated changes in lamellipodia phenotypes via phalloidin staining (Figure S8A, C, E, G, I, K). Cortactin knockdown did not result in notable differences in lamellipodia dimensions in WT or ArpC5 and ArpC5L KO cells, reminiscent of previous observations in murine Cortactin KO fibroblasts (*50*), but in contrast to observed changes in Vaccinia virus tail length in HeLa cells depleted of specific ArpC5 isoforms (*21*). We conclude that the changes in lamellipodial dimensions observed upon ArpC5 isoform knockouts are not mediated by differential cortactin activities.

### ArpC5 isoforms affect VACV actin tail length in a cell type-specific manner

Abella and colleagues reported VACV actin tails to be shorter after knockdown of ArpC5L and longer after knockdown of ArpC5 (*21*), respectively, representing an inverse phenotype compared to our observations of lamellipodia width in B16-F1 (Figure 1C-J) and Rat2 (Figure S4B-G) KO cell lines.

To determine whether the differential effects of ArpC5 isoform depletion on VACV actin tails and lamellipodia represent different regulatory mechanisms between these two experimental systems, we performed VACV tail length measurements in our B16-F1 and Rat2 WT and KO cell lines (Figure S9). In infected B16-F1 cells, VACV tail lengths were reduced in both C5KO and C5LKO cells compared to WT (Figure S9A-J). Interestingly, the tails observed in C5LKO cells were longer than those in C5KO cells. In Rat2 fibroblasts, we observed shorter tails in C5KO cells and longer tails in C5LKO cells compared to WT (Figure S9K-R). Hence, our observations for VACV tail length in an ArpC5/5L isoform-specific background are consistent with our lamellipodial phenotypes, although to variable extents when comparing Rat2 *versus* B16-F1 cells.

Considering our results and existing literature, ArpC5 isoforms clearly affect VACV actin tail formation but might do so in a cell type-specific manner. We interpreted this as clear hint towards additional factors being involved in contributing to the outcome of ArpC5 isoform-specific Arp2/3 complex activities, not previously anticipated.

### ArpC5 isoforms differentially regulate actin network architecture and branch junction structure

Aiming to understand the differences in the architecture of actin filament networks exhibited by C5KO and C5LKO cells, we used our published cryo-ET specimen preparation protocols (*13, 51*) to acquire tilt-series of vitreously frozen lamellipodia in B16-F1 WT, C5KO and C5LKO cells (Figure 3A-C). We then performed a quantitative ultrastructural analysis of actin network architectures in all cell types (Figure 3D- H) using a recently published workflow (*52*). This analysis revealed clear differences in angular distributions of actin filaments between WT and isoform-specific KO lamellipodia (Figure 3G-H) and provided an ultrastructural explanation for the respective lamellipodial phenotypes. In C5KO cells, a larger fraction of filaments was oriented parallel to the leading edge. Such filaments do not contribute to the formation of a prominent network with high width (*38*), but rather promote the narrow appearance of the lamellipodium typical for this genotype (Figure 1C- F). Correspondingly, in ArpC5L KO cells, a larger number of filaments were found to be oriented almost perpendicular to the leading edge (Figure 3F, H), leading to a wider lamellipodium as observed using fluorescence microscopy (Figure 1G-J). One possible explanation for the altered network architectures could be a different nucleation activity of isoform-specific Arp2/3 complexes, resulting in different numbers of branch junctions. To evaluate this possibility, we exploited the high-resolution molecular information present within our cryo-ET data and used classification-corrected template matching (see Methods) to identify branch junctions within our cryo-electron tomograms. We calculated branch junction densities per square micron in lamellipodia of each cell type and found the numbers of branch junctions with medians of 252.9 µm^-2^, 242 µm^-2^, and 258.7 µm^-2^ for WT, C5KO and C5LKO, respectively, to be comparable across all three genotypes (Figure 3I).

**Figure 3:**
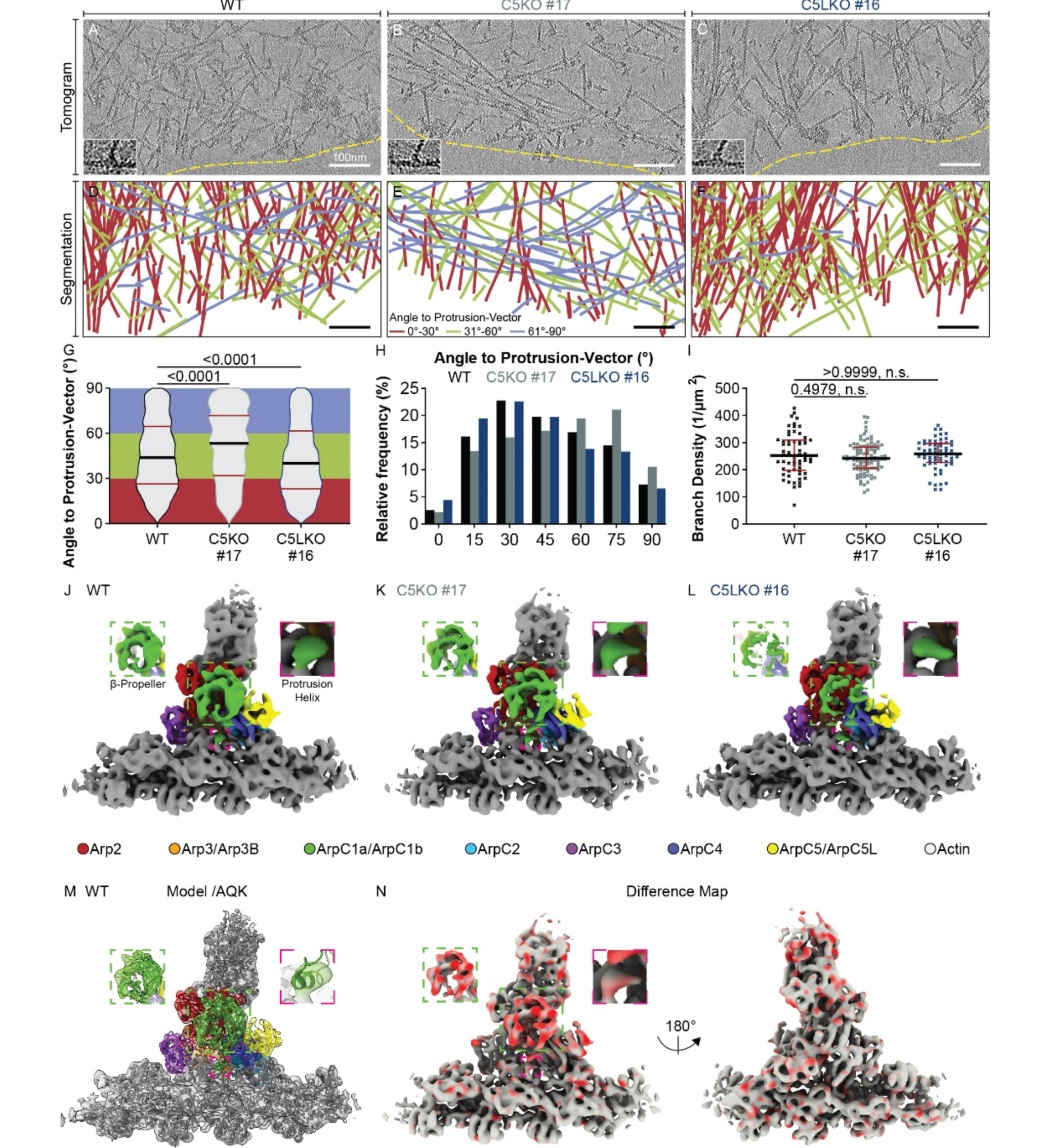
ArpC5 isoforms govern actin network architecture of the lamellipodium and have differential effects on ArpC1 stabilization in the branch junction. **(A-C)** Sums of 7 computational slices (∼9.5nm thickness) through cryo-electron tomograms of lamellipodia of B16-F1 wildtype, C5KO, and C5LKO cells, respectively. Dashed yellow lines indicate the leading edge. Insets show an enlarged view of representative branch junctions. **(D-F)** Segmentations of the actin filaments contained in the full height of the tomograms shown in (A-C). Color coding is given in (E). **(G-H)** Visualization of the angular distribution of actin filaments with respect to the vector of protrusion, pooled from 13 tomograms of B16-F1 wildtype, C5KO, and C5LKO cells each. **(G)** Violin plot. **(H)** Histogram. Kruskal-Wallis test combined with Dunn’s multiple comparison test, n= 2819, 2195, 2410 filaments, p values shown in the chart. Red lines indicate medians (43.93°, 53.27°, 40.20°), and black lines indicate quartile ranges. **(I)** Branch density per tomogram in lamellipodia of B16-F1 wildtype, C5KO, and C5LKO cells. Kruskal-Wallis test combined with Dunn’s multiple comparison test, n=61, 97, 61 tomograms, p values shown in the chart. Red lines indicate medians (252.9μm^-1^, 242.0μm^-1^, 258.7μm^-1^), and black lines indicate quartile ranges. **(J-L)** Isosurface representation of the actin filament - Arp2/3 complex branch junction derived from B16-F1 wildtype, C5KO and C5LKO cells. All structures have been low-pass filtered to 8.1 Å resolution. Isosurface representations are colored to visualize the molecular identities of Arp2/3 complex subunits and actin filaments. Subunit colors are annotated in the figure. Inlays show the ß-propeller (green dashed box) and the protrusion helix (pink dashed box) of the ArpC1 subunit. **(M)** The electron microscopy density map from B16-F1 wildtype cells (shown transparent) with the rigid body fitted model of the branch junction (pdb 7AQK). Color code as in (J-L). **(N)** A difference map (shown in both side views) was calculated from the structures of C5KO and C5LKO branch junctions projected onto the map of the C5KO branch junction. Red coloring indicates differences in occupancy. All scale bars, 100nm, n.s. = not significant.

This is also consistent with our observation that relative abundances of Arp2/3 complex subunits are not strongly altered upon removal of either ArpC5 or ArpC5L (Figure S5). As our results suggested that overall branch junction formation in lamellipodia is not significantly impacted by the selective incorporation of either ArpC5 or ArpC5L, we speculated that the observed differences in Arp2/3 complex activity might be caused by structural differences at the level of individual branch junctions. To test this, we performed subtomogram averaging and multi-particle refinement on identified branch junctions, yielding 8.0Å, 8.1Å, and 7.8Å resolution structures of actin filament Arp2/3 complex branch junctions from B16-F1 WT, ArpC5KO, and ArpC5LKO cells, respectively (Figure 3J-L, Figure S10). All three structures revealed an overall identical branch geometry of ∼70° and corresponded to recently published models of the Arp2/3 complex branch junction (*12, 13*)(Figure 3J-M, Supplementary Movie S1). While the overall branch junction was unaltered, we did observe a striking difference in the structure derived from C5LKO cells. In this structure, the density of the ArpC1 β-propeller was considerably weaker (Figure 3L), an observation further confirmed by calculating a difference map between branch junction structures derived from C5KO and C5LKO cells (Figure 3N). In contrast, the density for the so-called ArpC1 protrusion helix, which tethers ArpC1 to the actin mother filament (*12, 13*), displayed an equally strong density among all three structures (Figure 3J-L, N). These observations suggest that the lack of ArpC1 density in the C5LKO structure is not due to reduced occupancy, but could be due to decreased stability or increased flexibility in its β-propeller region. Our finding that branch junctions in cells lacking ArpC5L display higher flexibility is partially in line with the observation that ArpC5L-containing branches are suggested to be preferentially stabilized by Cortactin (*21*). However, the lack of effects on lamellipodium width upon Cortactin depletion in our isoform-specific KO cells (Figure S8) points towards additional mechanisms of branch junction stabilization.

### ArpC5 isoforms define lamellipodial actin network dynamics and protrusion characteristics

To explore how ArpC5 and ArpC5L regulate actin network dynamics in lamellipodia, we performed live-cell imaging and fluorescence recovery after photobleaching (FRAP) assays (Movie S2-4). In random migration assays, B16-F1 WT and ArpC5LKO cells exhibited comparable average velocities, while C5KO cells were significantly slower (Figure 4A). Correspondingly, C5KO cells, while exhibiting comparable retrograde flow to WT, displayed significantly lower protrusion velocity and thus lower total actin polymerization rate (Figure 4B-E). In contrast, C5LKO cells exhibited a stronger retrograde flow while having comparable overall polymerization rates to WT cells (Figure 4B-E). Even though this meant that both KO cell lines exhibited a lower protrusion efficiency than WT (Figure 4F), the considerably higher polymerization rate found in C5LKO compared to C5KO (Figure 4E) again clearly marked a difference between the two isoforms and is well in line with the wider lamellipodia observed in C5LKO compared to WT (Figure 1J).

**Figure 4:**
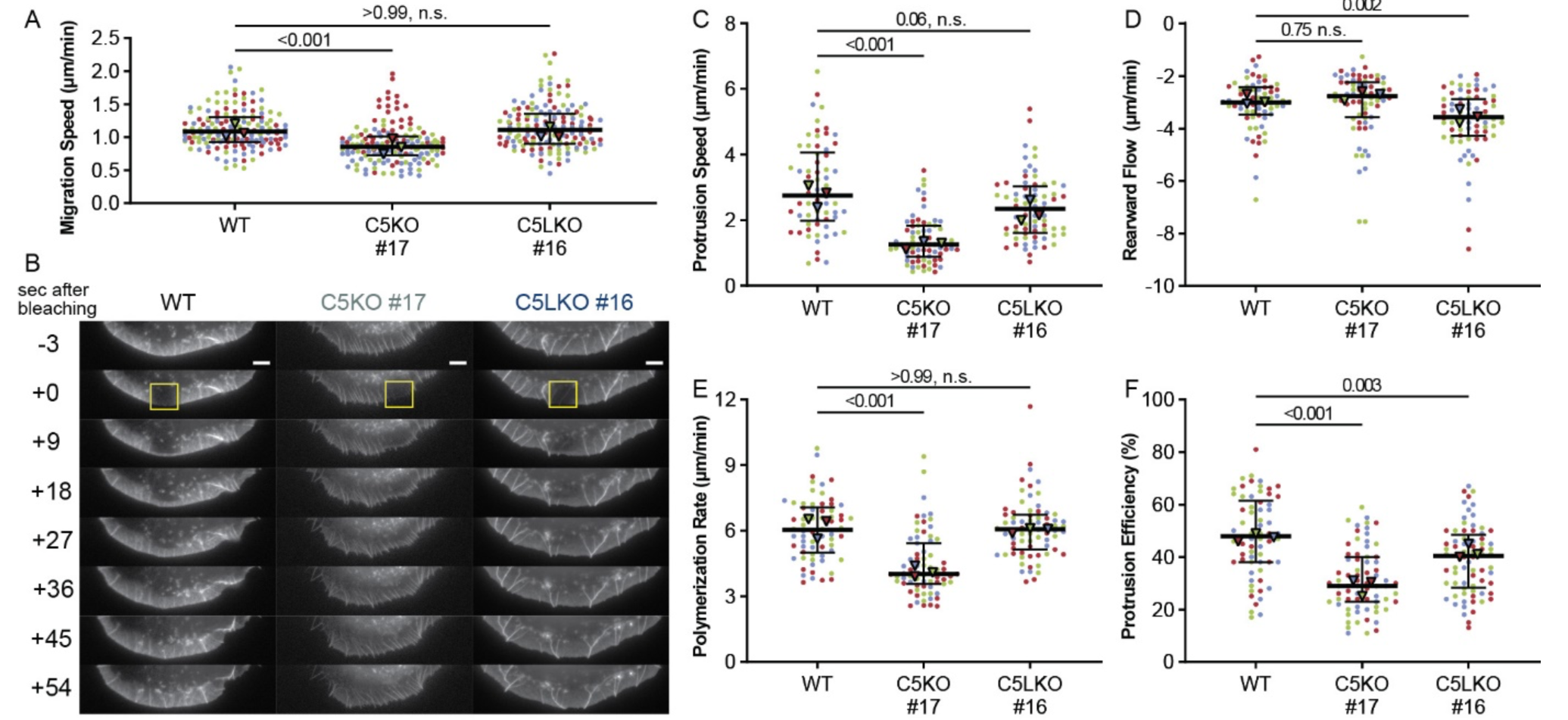
ArpC5 isoforms modulate random migration speed and lamellipodial actin dynamics. **(A)** Random migration speed of B16-F1 wildtype, C5KO and C5LKO cells. Kruskal-Wallis test combined with Dunn’s multiple comparison test on pooled data from 3 independent experiments, n=150 cells per experimental condition, p values shown in the chart. Black lines indicate overall medians (1.085μm/min, 0.8550μm/min, 1.115μm/min) and quartile ranges. **(B)** Representative time-lapse images of lamellipodia in B16-F1 wildtype, C5KO, and C5LKO cells expressing EGFP-actin, showing fluorescence recovery after photobleaching (FRAP). Yellow rectangles indicate FRAP regions. **(C-F)** Protrusion speed (C), rearward flow (D), polymerization rate (E), and protrusion efficiency (F) as determined via FRAP from B16-F1 wildtype, C5KO and C5LKO cells expressing EGFP-actin. Polymerization rate is calculated as the sum of the absolute protrusion speed and the absolute rearward flow. Protrusion efficiency is calculated by dividing protrusion speed by polymerization rate. Kruskal-Wallis test combined with Dunn’s multiple comparison test on pooled data from 3 independent experiments, n=66, 71, 72 cells for each experimental condition, respectively, and with p values shown in the chart. Black lines indicate overall medians (protrusion speed: 2.747 μm/min, 1.258 μm/min, 2.348 μm/min; rearward flow: -3.003 μm/min, -2.768 μm/min, -3.570 μm/min; polymerization rate: 6.035 μm/min, 4.026 μm/min, 6.068 μm/min; protrusion efficiency: 48.00%, 29.00%, 40.50%) and quartile ranges. Data points in (A, C-F) are color-coded according to the individual experiments. All scale bars, 20µm. Triangles indicate the medians of the respective experiments. N.s. = not significant.

### Accumulation at the leading edge of the actin filament elongators Mena/VASP is reduced in the absence of ArpC5

The reduced actin polymerization rate in C5KO cells could be due to either diminished nucleation of new filaments or a reduced elongation of existing filaments. As we already determined that the number of branch junctions is comparable in all cell lines (Figure 3I) and previous work established that ArpC5L- containing Arp2/3 complexes can mediate more efficient actin assembly and branching (*21*), we excluded that Arp2/3 complex-dependent *de-novo* nucleation of filaments is impaired.

Instead, we speculated whether ArpC5 and ArpC5L- containing lamellipodia might exhibit differences in actin filament elongation. We, therefore, hypothesized that ArpC5/5L isoform-specific Arp2/3 complexes might be able to regulate the localization and/or activity of actin filament elongating proteins. We thus tested for the presence of the Ena/VASP family proteins at the leading edge (*53*). Indeed, Mena (Figure 5A-F) and VASP (Figure 5G-L) accumulated at the leading edges of B16-F16 WT (Figure 5A-B, G-H) and C5LKO cells (Figure 5E-F, K-L), but not in C5KO cells (Figure 5C-D, I-J). We corroborated these results via quantification (Figure 5M-O) and found the same pattern in Rat2 WT, C5KO, and C5LKO cells (Figure S11). To determine if the reduced localization of Mena and VASP is a specific effect of incorporation of ArpC5 isoforms into branch junctions and not a mere consequence of the reduced lamellipodium phenotype itself, we tested for the localization of additional lamellipodial tip components determining protrusion and lamellipodia formation.

**Figure 5:**
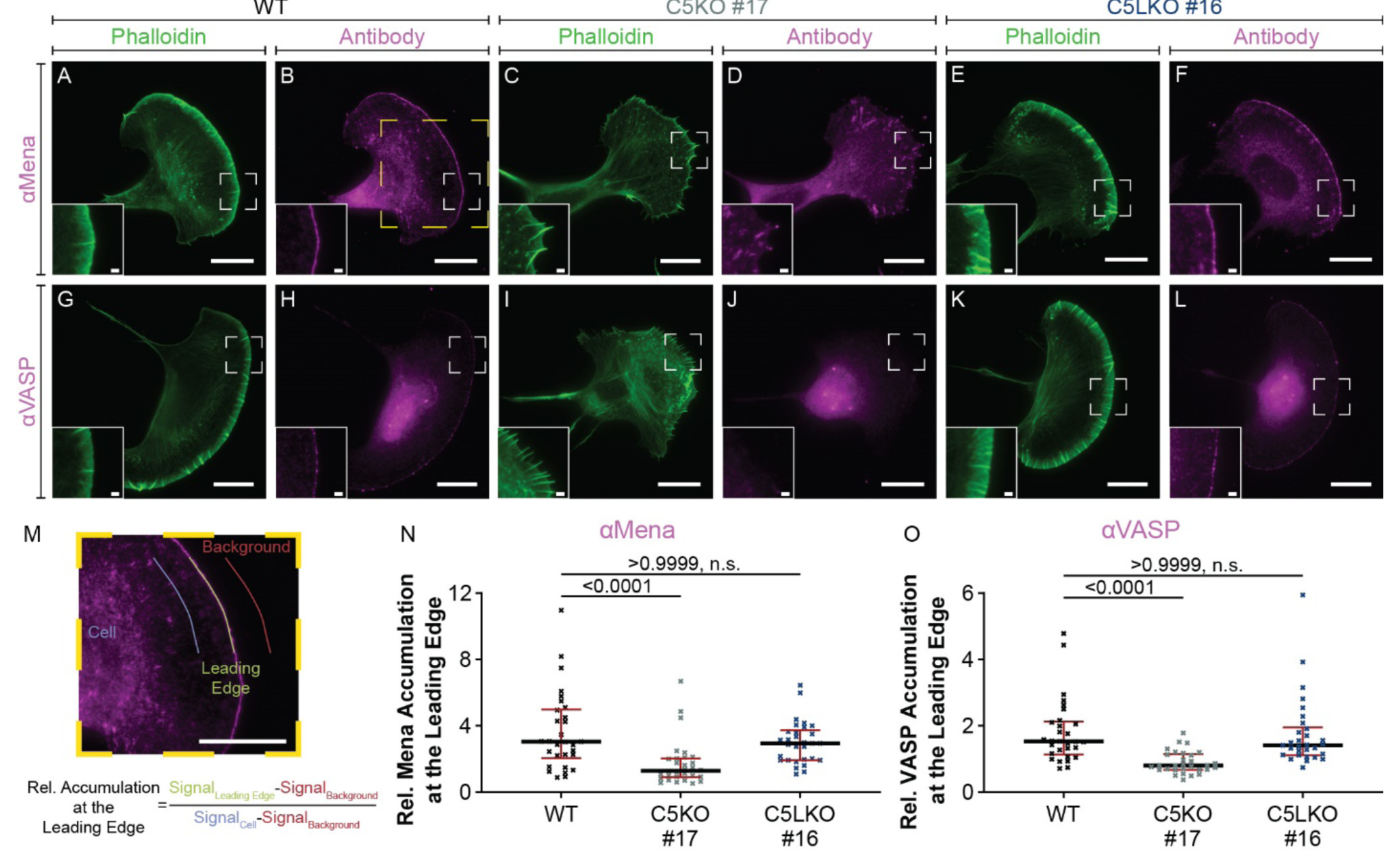
ArpC5 isoforms modulate Mena/VASP recruitment to the leading edge. **(A-L)** Representative epifluorescence micrographs of B16-F1 wildtype (A, B, G, H), C5KO (C, D, I, J), and C5LKO (E, F, K, L) cells visualizing the actin cytoskeleton using fluorescent phalloidin (A, C, E, G, I, K) and the localization of Mena (B, D, F) and VASP (H, J, L) by immunofluorescence. Inlays show magnified areas indicated in respective panels. **(M)** Zoom-in on the square region indicated in (B), visualizing the measurements and the calculations determining the relative accumulation of Mena/VASP at the leading edge. The leading edge was traced for at least 15µm per measurement. To measure the background and cell intensities, the leading edge region of interest was shifted by ∼10µm inside or outside cells while avoiding extremely bright areas caused by secondary antibody aggregates. **(N-O)** Quantitative analysis of the relative accumulation of Mena (N)/VASP (O) at the leading edges of B16-F1 wildtype, C5KO, and C5LKO cells. Kruskal-Wallis test combined with Dunn’s multiple comparison test, n=30 cells for each experimental conditions, for both Mena and VASP, p values shown in the chart. Red lines indicate medians (Mena (panel N): 3.044, 1.292, 2.954; VASP (panel O): - 1.533, 0.8091, 1.419) and black lines quartile ranges. Scale bars, 20µm in standard panels and 2µm in inlays, n.s. = not significant.

We observed a very minor reduction of WAVE2 localization at the leading edges of both C5KO and C5LKO (Figure S12) in stark contrast to the almost complete reduction of Mena and VASP in ArpC5KOs. These results provide evidence that the reduced localization of Ena/VASP family members is not simply dependent on the extent of active lamellipodial protrusion. Instead, it suggests that isoform-specific Arp2/3 complex subpopulations are directly capable of affecting the distribution of specific downstream effectors to determine actin filament elongation.

## Discussion

We have used an integrated cell and structural biology approach to show on the cellular, ultrastructural, and molecular level that the two distinct ArpC5 isoforms regulate cell migration differentially.

We show that C5KO cells exhibit impaired motility and lamellipodia formation, strongly altered actin architecture, and reduced recruitment of actin filament elongators of the Ena/VASP family to the leading edge (Figure 6). These results indicate that ArpC5 isoform composition can act as regulator of actin polymerization efficiency in lamellipodia through altering the extent of Ena/VASP protein accumulation at the leading edge. Our findings further suggest that the Ena/VASP-dependent elongation of ArpC5- containing Arp2/3 complex nucleated filaments is able to overcome the reduced nucleation efficiency of ArpC5-containing Arp2/3 complexes (Figure 6) (*21, 23*). This correlation between ArpC5 isoform levels and changed Ena/VASP localization provides a molecular explanation for the observation that elevated ArpC5 expression is associated with increased metastasis and poorer overall survival in different cancers (*41–43, 54*). It is further well in line with the observation that Ena/VASP protein levels and activity are also associated with cancer progression (*55–57*).

**Figure 6:**
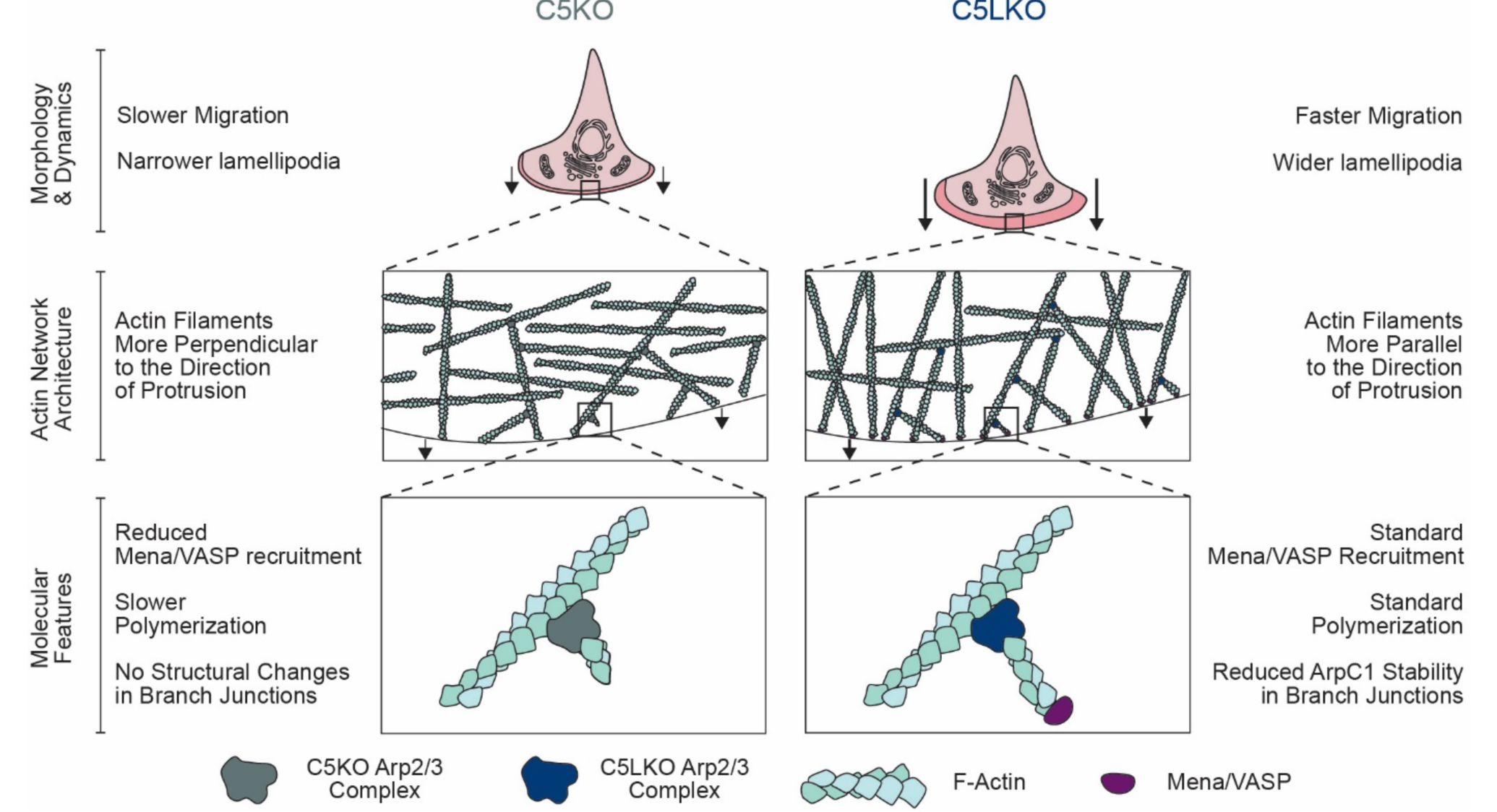
Schematic summary ArpC5 isoform-dependent regulation of lamellipodium characteristics and cell migration. Schematic representation of the phenotypes observed for C5KO and C5LKO cells and of how altered actin filament polymerization velocities caused by differential recruitment of Mena/VASP overcomes the intrinsically different nucleation speeds of the associated Arp2/3 complexes.

It is intriguing to speculate how an Arp2/3 complex of a given isoform-composition is able to define the elongation properties of the filament it has nucleated. Does ArpC5 specifically support the delivery of Ena/VASP, or does ArpC5L prevent the association of these elongation factors with barbed actin filament ends? A prior *in vitro* study using tissue-derived Arp2/3 complex reported the importance of the source tissue of Arp2/3 complex, which can determine the effects of VASP on WAVE-dependent actin polymerization (*58*). In the presence of thymus-derived Arp2/3 complex (high ArpC5/ArpC5L ratio), VASP boosted the otherwise low actin polymerization speed significantly. In contrast, VASP lowered the polymerization of actin in the presence of brain-derived Arp2/3 complexes (low ArpC5/ArpC5L ratio), which appeared to have higher baseline nucleation activity. These findings suggest an antagonistic cross-regulation between ArpC5L and Ena/VASP proteins in WAVE-dependent actin networks, which is consistent with our observations. However, further studies are needed to fully unravel the cross-regulation between variable isoform-dependent Arp2/3 complex activities and the resulting downstream actin filament elongation of newly nucleated filaments.

Despite their isoform-specific differences, the shared functions of ArpC5 and ArpC5L are vital for Arp2/3 activity in cells, as we can show that lack of both isoforms results in complete depletion of lamellipodia (Figure S3). In addition, earlier studies have shown the roles of ArpC5/5L in Arp2/3 complex assembly, stabilization, and nucleation activity (*21, 23, 45*). *In vitro* reconstitution experiments suggested that ArpC5/ArpC5L are involved in integrating ArpC1 into the complex (*45*). Our structures of isoform-specific branch junctions obtained from cells further emphasize such a functional connection between ArpC1 and ArpC5/ArpC5L, as we observe an increased destabilization of ArpC1 within ArpC5-containing Arp2/3 complex branch junctions. Nevertheless, we cannot comment on whether this affects either one or both of the ArpC1 isoforms present in the branch junction. Weakened interfaces between ArpC1 and ArpC5 could be causative for the observed increase in ArpC1 flexibility. However, the resolution of our structures is too low to unambiguously define the interactions that form the ArpC1/ArpC5 interface. A recent single-particle cryo-EM study solving structures of *in vitro* reconstituted, inactive Arp2/3 complexes at resolutions between 4 and 5 Å showed increased flexibility of ArpC5L compared to ArpC5. Still, this observation cannot fully explain a changed ArpC1 stability as we did observe stable ArpC1 in ArpC5L- containing branches. We also did not observe pronounced differences in the densities between ArpC5 and ArpC5L in our isoform-specific branch junction reconstructions from cells. This suggests that ArpC5 isoforms might be differentially stabilized in the inactive and active Arp2/3 complex and that the stabilization of ArpC1 might not be entirely dependent on ArpC5 isoforms. Coronins are known to promote debranching, and it was suggested that multimeric Coronin1B replaces Arp2/3 complexes at the branch junction (*19*). ArpC1 and Coronin share a similar beta-propeller fold, allowing to hypothesize that there might be a competitive displacement of ArpC1 by Coronin depending on which ArpC5 isoform is incorporated into the Arp2/3 complex. However, what specifically regulates ArpC1 stability and how ArpC1 stability affects the branch junction as a whole still needs to be established. Future structural studies, either *in vitro* on reconstituted, active Arp2/3 complexes (*12, 59*) or *in situ*, can provide further insights into how ArpC1 and ArpC5 isoform-specific branches are regulated by branch junction stabilizing proteins, such as Cortactin or branch junction- disassembling proteins, such as GMF (*60*) and Coronin 1b.

Our results show that the ArpC5 isoform is clearly associated with more prolific actin assemblies in lamellipodia of Rat2 fibroblasts and B16-F1 melanoma cells. These observations are in line with recently published work revealing that ArpC5 has the same effect in peripheral actin rings of CD4 T cells (*61*). This supports the notion that ArpC5/5L isoform-specific activities on cellular actin networks are generalizable over different cell types and species. However, ArpC5/5L isoforms showed cell-type-specific differences in VACV actin tail formation in B16-F1 melanoma cells, Rat2 fibroblast, and HeLa cells. Specifically, in HeLa cells, the ArpC5/5L isoform-dependent actin tail length phenotype was inverted compared to B16-F1 and Rat 2 cells. We note that different cell lines can have different protein expression levels for the ArpC5 isoforms (Figure S2, S6). However, our observations that overexpression of either isoform is not able to rescue isoform-specific KO effects argues that Arp2/3 dosage is not the main cause for differing VACV tail lengths in the used cellular systems. Further work is required to establish the molecular determinants that define VACV actin tail formation in different cell lines and species.

CRISPR/Cas9-mediated genome editing has revolutionized all aspects of cell biology and offers a substantial potential for cellular structural biology. In the present study, we were able to integrate structural and cell biology assays in one model system to obtain a better understanding of Arp2/3 complex regulation. Using cryo-ET and subtomogram averaging on genetically engineered cells, we determined structures of isoform-specific Arp2/3 complex branch junctions at ∼8 Å resolution, revealing relevant differences in branch junction stability. This study underscores how the results of cellular cryo-ET can be complemented by a large number of experimental techniques applicable to cells. Furthermore, our work highlights the capabilities of cryo-ET to study challenging biological assemblies, which have remained difficult to study with standard reductionist structural biology approaches (*62*).

## Acknowledgments

This research was supported by the Scientific Service Units (SSUs) of ISTA through resources provided by Scientific Computing (SciComp), the Life Science Facility (LSF), the Imaging and Optics facility (IOF) and the Electron Microscopy Facility (EMF). The authors would like to thank Kristin von Peinen and Brigitte Denker (Helmholtz Centre for Infection Research, Braunschweig, Germany) for experimental and technical assistance, respectively. The authors acknowledge support from ISTA and from the Austrian Science Fund (FWF): P33367 to F.K.M.S, from the Deutsche Forschungsgemeinschaft (DFG), Research Training Group GRK2223 and the Helmholtz Society to K.R..

## Data availability statement

The EM density maps of the WT and ArpC5/5L KO actin filament Arp2/3 complex branch junctions and one representative tomogram per genotype have been deposited in the EMDB (https://www.ebi.ac.uk/pdbe/emdb) under accession numbers: EMD-15136, EMD-15137, EMD-15138, EMD-15139, EMD-15140, and EMD-15141.

This data is the basis for Figure 3, Supplementary Figure 10 and Supplementary Movie 1.

### Contributions

Project administration: F.K.M.S; supervision and funding acquisition: K.R. and F.K.M.S.; conceptualization: F.F.,

K.R. and F.K.M.S; methodology: F.F., F.K.M.S; investigation: F.F., M.G.J, J.D., H.D., F.W.H, G.D., and V.-V.H.; validation, formal analysis, and visualization: F.F., M.G.J, J.D, H.D.; data curation: F.F., K.R. and F.K.M.S; writing— original draft: F.F. and F.K.M.S.; writing—review and editing: F.F., M.G.J, J.D., H.D., K.R., and F.K.M.S.

## Materials and Methods

### Antibodies and phalloidin

Primary antibodies were used in the following concentrations: Anti-ArpC5/C5L (Thermo Fisher Scientific #PA5- 30352) from rabbit, 1:5000 for Western blot; Anti-ArpC5 (Synaptic Systems #305 011) from mouse, 1:200 for immunostaining; Anti-ArpC5L (Abcam # ab169763) from rabbit, 1:100 for immunostaining; Anti-ArpC1a (Abcam, #ab133160) from goat, 1:2500 for Western blot; Anti-ArpC1b (Santa Cruz, #sc-271342) from mouse, 1:250 for Western blot; Anti-Arp2 (Abcam, #ab226476) from rabbit, 1:5000 for Western blot; Anti-Arp3 (Abcam, # ab181164) from rabbit, 1:5000 for Western blot; anti-B5 antibody (MAb VMC-20 (*63, 64*), kindly received from Gary Cohen, University of Pennsylvania) from mouse 1:100 for immunostaining; Anti-Mena (Merck, #HPA028696) from rabbit, 1:2500 for Western blot and 1:200 for immunostaining; Anti-VASP (Merck, # HPA005724) from rabbit, 1:2500 for Western blot and 1:200 for immunostaining; Anti-Actin (Merck, #A1978) from mouse, 1:5000 for Western blot; Anti-GAPDH (Abcam, #ab125247) from mouse, 1:5000 for Western blot; Anti-Vinculin (Merck, #V9131) from mouse, 1:5000 for Western blot; Peroxidase AffiniPure Goat Anti-Rabbit IgG (Jackson Immuno Research #111-035-144), Peroxidase AffiniPure Goat Anti-Mouse IgG (Jackson Immuno Research #115-035-146) and Peroxidase AffiniPure Donkey Anti-Goat IgG (Jackson Immuno Research #705-035- 003) were used in a dilution of 1:5000 as secondary antibodies for detection in Western blot applications. m-IgGk BP-CFL 488 (Santa Cruz, # sc-516176) and Anti-Rabbit-IgG-Atto 594 (Merck, #77671-1ML-F) were employed in 1:200 and 1:500 dilution, respectively, as secondary antibodies for immunostaining. Phalloidin-Atto488 was used complementary with Anti-Rabbit-IgG-Atto 594, and Phalloidin-Atto594 was used with m-IgGk BP-CFL 488 in 1:500 dilution to visualize actin filaments.

### Cloning dual expression plasmids

mRFP was amplified from aleu-mRFP (*65*) using GAGGACGTCGACATGGCCTCCTCCGAGGACGTCATCA/ GTCCTCACTAGTTTATGCTCCAGTACTGTGGCGGCCC and inserted into the second multiple cloning site of PSF-CMV-CMV-SBFI-UB-PURO (Merck, #OGS597-5UG) using SalI/SpeI restriction digest and ligation. This yielded pCMV::empty_pCMV::RFP. ArpC5 and ArpC5L were amplified from an Mus musculus NIH-3T3 fibroblast (RRID:CVCL_0594) cDNA library using GAGAACACATGTCGAAGAACACGGTGTC/ ATCGGGATCCCTACACGGTTTTCCTTGCAGT or GAGAACCCATGGCCCGGAACACACTGTC/

GTCCTCGGATCCTTAAACAGTCTTTCTTGCTGTA, respectively. ArpC5 was digested using PciI/BamHI and inserted into the first multiple cloning site of NcoI/BamHI-digested pCMV::empty_pCMV::RFP by ligation yielding pCMV::ArpC5_pCMV::RFP. ArpC5L was digested using NcoI/BamHI and inserted into the first multiple cloning site of NcoI/BamHI-digested pCMV::empty_pCMV::RFP by ligation yielding pCMV::ArpC5L_pCMV::RFP.

### Cell culture and KO line generation

Wildtype Mus musculus (Mm) B16-F1 (RRID: CVCL_0158) melanoma cells and Rattus norvegicus (Rn) Rat2 fibroblasts (RRID: CVCL_0513) were used in this study. B16-F1 cells were cultured in DMEM GlutaMAX (Thermo Fisher Scientific, #31966047), supplemented with 10 % (v/v) fetal bovine serum (Thermo Fisher Scientific, #10270106) and 1 % (v/v) penicillin-streptomycin (Thermo Fisher Scientific, #15070063). Rat2 fibroblasts were cultured in the same medium with an added 1 % (v/v) of 100X MEM Non-Essential Amino Acids Solution (Thermo Fisher Scientific, # 11140068). Cells were incubated at 37 °C and 5 % CO2. Detachment of cells prior to seeding was performed with 0.05 % (Thermo Fisher Scientific, #25300054) and 0.25 % (Thermo Fisher Scientific, #25200056) trypsin for B16-F1 cells and Rat2 fibroblasts, respectively.

ArpC5 and ArpC5L genes were knocked out individually in B16-F1 and Rat2 cells by CRISPR/Cas9-mediated genome editing resulting in C5KO and C5LKO lines (Cong et al., 2013). The B16-F1 C5/C5L double KO lines were generated by disrupting ArpC5 in the B16-F1 C5LKO #16 background. The corresponding guide sequences GATATGACGAGAACAAGTTCG for MmArpC5, GATTCGTAGACGAGCACGAAG for MmArpC5L, GTCGTCGGCCCGCTTCCGGAAGG for RnArpC5 and GATCCACTCGGCGGAAGCGTGAGG for RnArpC5Lwere cloned into pSpCas9(BB)-2A-Puro (Addgene, ID 48139). Cells were transfected employing Lipofectamine™ LTX Reagent with PLUS™ Reagent (Thermo Fisher Scientific, # 15338100). 16 h after transfection, cells were reseeded and enriched by selection in their respective media with an added 2.5 µg/ml of puromycin for B16-F1 and 3 µg/ml of puromycin (Merck # p9620) for Rat2 cells, respectively, for 3 d. Surviving cells were diluted (∼1:50) and incubated in non-selective media to allow for the formation of single-cell-derived colonies. Approximately 7 d after seeding, individual colonies were selected and expanded for genotyping, first by Western blot, then by genomic sequencing of the target loci (Figures S1, S2).

### Western blotting

Cyt-Ex buffer containing 22.5 % (v/v) 4X Laemmli buffer (Bio Rad, #1610747), 2.5 % (v/v) 2-mercaptoethanol (Bio Rad, #1610710), 75 % (v/v) Cyt-Ex-Pre buffer (10 mM Tris, 100 mM NaCl, 1 mM EDTA, 1 mM EGTA, 1 % (v/v) Triton X-100, 0.1 % (w/v) SDS, adjusted to pH 7.4) and cOmplete™, EDTA-free protease inhibitor cocktail (Merck, #11873580001) was prepared shortly prior to sample extraction.

Cells were washed with pre-chilled PBS and lysed with Cyt-Ex buffer on ice for 10 min prior to scraping and incubation at 95 °C for 10 min. For estimating protein concentration, 10 µl of sample were transferred into 100 µl of Amido-Black-Staining solution (0.25 % (w/v) Amido black 10 B (Merck, #1011670025)), 45 % (v/v) MeOH, 45 % (v/v) ddH2O, 10 % (v/v) glacial HOAc. After sedimentation at 20,000g for 3 min the supernatant was removed and replaced with 150 µl of Amido-Black-Destaining solution (45 % (v/v) MeOH, 45 % (v/v) ddH2O 10 % (v/v) glacial HOAc). After another round of sedimentation at 20,000 g for 3 min the supernatant was removed and the sediment was dissolved in 200 µl of 0.2 M NaOH. 100 µl of the solution was then employed for photometric concentration measurement at 600nm absorption using a TECAN Infinite F500 plate reader (Tecan). 10 µg of extracts per sample were run on SDS–PAGE gels, transferred to PVDF membranes (AL Labortechnik, 42514.01) via wet blotting, and blocked in blocking solution (4 % (w/v) skim milk powder (Merck, #70166-500G) and 1 % (w/v) BSA (Merck, #10735078001) in Tris-based saline containing 0,05% Tween 20 (TBST)) at room temperature (RT) for 1 h. All following washings were performed in TBST in three 5 min steps. Blots were washed and incubated in primary antibodies solved in Antibody-Dilution solution (2 % (w/v) BSA and 0.02 % (w/v) sodium azide in TBST) at 4 °C overnight. Blots were washed again and incubated in secondary antibody diluted in blocking solution at RT for 45 min. Blots were washed one last time and then stored in TBS until detection. ECL1 solution (100µl 250mM Luminol (Merck, #A8511-5G, solved in DMSO) and 44µl 90mM p-coumaric acid (Merck, #C9008-5G, solved in DMSO) dissolved in 1ml 1M Tris pH 8.5, then brought to 10 ml with ddH2O) and ECL2 solution (6 µl 30% (v/v) H2O2 (Merck, #H1009) in 10 ml with ddH2O) were prepared directly prior to application. The chemiluminescence signals were developed in a 1:1 mixture of ECL1 and ECL2. Images were acquired using an Amersham Imager 600 (GE Healthcare). Uncropped versions of all Western blot results are represented in (Figure S13)

### VACV infection assay

For VACV infection assays, cells were infected with VACV (Western reserve strain, kindly provided by Andreas Bergthaler, Medical University of Vienna) for 1 h, washed three times with PBS, and incubated in culture medium for another 7 h. VACV infected cells were fixed with pre-warmed 4% paraformaldehyde (Merck, #P6148) in PBS for 1 h to ensure the inactivation of the virus. The cells were washed three times with PBS and subjected to staining with Phalloidin-Atto594 and a previously characterized monoclonal anti-B5 antibody (*63, 64*) prior to embedding in ProLong™ Gold Antifade Mountant (Thermo Fisher Scientific, #P36934).

### Immunofluorescence and phalloidin staining

For rescue experiments (Figure 2), cells were transfected using 300 ng of dual expression plasmid and Lipofectamine™ LTX Reagent with PLUS™ Reagent (Thermo Fisher Scientific, # 15338100) and incubated 16 h prior to seeding. For immunofluorescence and phalloidin stainings, cells were trypsinized and seeded onto coverslips that were coated with 25 µg/ml laminin (Merck, #11243217001) in laminin coating buffer (50mM TRIS, 150mM NaCl, pH 7.4) at RT for 1 h. Cells were allowed to settle on coverslips for 16 h at 37 °C and 5 % CO2 before they were either employed for infection assays or directly fixed.

For fixation of non-VACV-infected samples, the medium was aspirated and immediately replaced by pre-warmed 4 % paraformaldehyde (Merck, #P6148) in PBS. After 20 min of fixation, cells were washed twice with PBS prior to being permeabilized via incubation with 0.1 % Triton X-100 in PBS for 1 min. Permeabilized cells were washed twice with PBS, subjected to the indicated staining, and embedded using ProLong™ Gold Antifade Mountant (Thermo Fisher Scientific, #P36934).

Non-VACV-infected samples were imaged on a Zeiss Axio Imager (Zeiss) equipped with a CoolLED p300 SB light source (CoolLED) and an LCI Plan-Neofluar 63x/1,3 Imm Corr DIC M27 objective (Zeiss).

VACV-infected samples were imaged on a Zeiss Axio Observer (Zeiss) equipped with 405, 488, 561, and 640 nm laser lines and a Plan-APOCHROMAT 63x/1.4 Imm Corr DIC Oil M27 objective (Zeiss).

Conversion of image formats, generation of merges/overlays, and distance measurements were performed in Fiji v1.52p (*66*).

### Cortactin knockdown

For the Cortactin knockdown experiments, cells were transfected in 96-Well plate wells using either 25 nM siGENOME Non-Targeting Pool #2 (Horizon Discovery, #D-001206-14-05) or 25 nM Cortactin siRNA SMARTPool (Horizon Discovery, #L-044721-00-0005) using 0.5 µl DharmaFECT 1 Transfection Reagent in a total volume of 100 µl. A second round of transfection of the same cells was performed according to the same protocol after 24h. Cells were then incubated for 24h and subsequently either seeded for immunostaining onto coverslips that were coated with 25 µg/ml laminin or for Western blot sample preparation into 12-Well plate wells. Cells were allowed to settle for 24h before they were subjected to immunostaining or Western blot (for details, see above).

### Random migration assay

B16-F1 cells were seeded onto µ-slide 8-well glass-bottom microscopy chambers (Ibidi, #80807) coated with 25 µg/mL laminin. Live-cell time-lapse image series were acquired with an Eclipse-Ti2 (Nikon) equipped with a Plan Apo λ 20x/0.75 DIC air PFS objective (Nikon) and operated with Nikon’s NIS-Elements imaging software. A custom-made environment chamber was used to maintain constant conditions of 37 °C and 5 % CO2. DIC time-lapse images of randomly chosen cell-containing areas were acquired for 15 h every 10 min. Trajectories of individual cells were tracked manually in Fiji v1.47v employing the built-in Fiji Manual Tracking plugin. For deriving the average speed of individual cells, trajectories were analyzed using the Chemotaxis and Migration Tool (Ibidi, https://ibidi.com/chemotaxis-analysis/171-chemotaxis-and-migration-tool.html).

### Fluorescence Recovery after Photobleaching (FRAP)

B16-F1 cells were transfected with 0.5 µg EGFP-β-actin and 1 µL JetPrime transfection reagent (VWR, #114-07) per well in 6-well plates according to the manufacturer’s instructions. 16 hours post transfection, cells were seeded onto µ-slide 8-well glass-bottom microscopy chambers (Ibidi, #80427) coated with 25 µg/mL laminin as described in the section Immunofluorescence and phalloidin staining. Live-cell time-lapse image series were acquired with a 100x/1.4 apochromatic objective on an inverted Axio Observer (Zeiss) featuring an incubation chamber heated to 37°C (Warner instruments), a DG4 light source (Sutter instruments), and a CoolSnap HQ2 camera (Photometrics). Randomly chosen EGFP-actin expressing cells with steadily protruding lamellipodia were imaged at 3 sec per frame with 500 ms exposure time. After 4-5 frames, a rectangular region containing a subsection of the lamellipodium was bleached (see ROIs in Figure 4) using a 405 nm diode laser at 80 mW output power, controlled by a 2D-VisiFRAP Realtime scanner and driven by VisiView software (Visitron Systems). Individual recordings were analyzed using Fiji (v2.1.0). Image stacks were resliced along a line in the bleached region perpendicular to the cell leading edge, creating a kymograph. Retrograde flow and protrusion rate values were obtained by calculating the speed of the recovering actin network moving back or forth, respectively. The total actin polymerization rate was determined by adding the absolute values of retrograde flow and protrusion rate. The ratio of protrusion to total actin polymerization is expressed as protrusion efficiency.

### Cryo-electron tomography

Cells were trypsinized and seeded onto 200 mesh gold holey carbon grids (R2/2-2 C, Quantifoil Micro Tools) placed in 3D printed grid holders (*51*) for cryo-ET specimen preparation. Before seeding, grids were glow discharged using an ELMO glow discharge unit (Cordouan Technologies) for 2 min and then coated with 25 µg/ml laminin for 1 h at RT. Cells were allowed to adhere to the grids at 37 °C and 5 % CO2 for 4 h before they were extracted and fixed according to published protocols (*13, 16, 51*). In detail, grids were removed from grid holders and placed in 50 μl drops of cytoskeleton buffer (10 mM MES, 150 mM NaCl, 5 mM EGTA, 5 mM glucose, and 5 mM MgCl2, pH6.2) supplemented with 0.75 % Triton X-100, 0.25% glutaraldehyde (Electron Microscopy Services, #E16220) and 0.1 µg/ml phalloidin (Merck, #P2141) for 1 min at RT. Grids were then transferred to 50 μl drops of cytoskeleton buffer containing 2 % (w/v) glutaraldehyde and 1 µg/ml phalloidin at RT for 15 min.

Subsequently, grids were vitrified using a Leica GP2 plunger (Leica Microsystems) with blotting chamber conditions of 80% humidity and 4 °C. In the blotting chamber, the fixation solution was manually blotted and replaced with 3 μl 10 nm colloidal gold coated with BSA suspended in PBS for WT or 3 μl PBS for C5KO and C5LKO, respectively. After backside blotting for 3 s using the internal blotting sensor of the Leica GP2, the grids were vitrified in liquid ethane (−185 °C) and then stored under liquid nitrogen conditions.

Tilt-series were acquired under cryogenic conditions on a Thermo Fisher Scientific Krios G3i TEM equipped with a BioQuantum post-column energy filter and a K3 camera (Gatan), using SerialEM v3.8 (*67*). Acquisition of low- and medium-magnification maps allowed for defining areas of interest for subsequent high-resolution tomography data acquisition. Gain reference images were collected before data acquisition. DigitalMicrograph as integrated into the Gatan Microscopy Suite v3.3(Gatan) and SerialEM were used for filter and microscope tuning. Tilt-series were acquired with a filter slit width of 20 eV, and using a dose-symmetric tilt-scheme (*68*) ranging from −66° to 66° with a 2° increment. The nominal defocus range was set to −1.5 to −4.5 µm and the nominal magnification to 53,000x, resulting in a pixel size of 1.693 Å. Tilt-images were acquired as 5,760 × 4092 pixel movies of eight frames. The dose per tilt was set to be 2.79 e/Å² resulting in a calculated cumulative dose of 170 e/Å² for the first 61 tilts per series. Data sets were acquired in separate acquisition sessions contributing 61 (wildtype), 97 (C5KO), and 61 (C5LKO) tilt-series, respectively. For data acquisition settings, see Figure S10B.

### Cryo-ET data processing

Gain correction, frame alignment and defocus estimation of individual tilts were performed in Warp v1.0.9 (*69*). Before exporting tilt-series as .mrc stacks, poor quality tilt-images caused, for example, by grid bars blocking the beam at high tilt-angles, were removed. Tilt-series alignment was done using patch tracking within the IMOD software package v4.11 (*70*).

Alignment parameters as determined in IMOD were imported into Warp. The metrics necessary for dose-dependent low pass filtering were calculated during this step. Defocus parameters were then again determined for the whole tilt-series. 8×-binned (13.544 Å/px) tomograms were exported for template matching in the Dynamo package v1.1.333 (*71*). For template matching of branch junctions, a previously used template (*13*) was resampled to a pixel size of 13.544 Å/px using RELION v3.0.8 (*72, 73*). For actin filaments, a template was generated from a model of aged, nucleotide-bound, and phalloidin-stabilized F-actin (pdb 6T20 from (*74*)) employing the molmap function in chimera. The box of the resulting map was extended using RELION to be cube-shaped.

For cross-correlation calculation during template matching, masks consisting of two cylinders (140 Å radius) covering the branch or of one cylinder surrounding the single actin filament, respectively, were applied. Angular scanning around all three Euler angles was performed for a full 360° with a sampling step of 10°. False-positive cross-correlation peaks from predetermined areas containing gray value outliers were automatically removed. Subsequently, 400 positions from the branch junction search and up to 1000 positions from the actin filament search with the highest cross-correlation values per tomogram were selected for further processing.

Particles from different data sets were processed independently. Dynamo2m scripts v0.2.2 (*75*) were employed to transform particle coordinates from Dynamo to RELION format. 24,400 wildtype / 38,800 C5KO / 24,400 C5LKO branch junction-containing particles (30 px side length, 13.544 Å/px) and 61,061 wildtype / 87,300 C5KO / 54,900 C5LKO) actin filament-containing particles (60 px side length, 6.722 Å/px) were exported using Warp. 3D classification in RELION was used to remove junk particles reducing the number of branch junction-containing particles to 9,084 wildtype / 15,339 C5KO / 9,831 C5LKO and actin filament-containing particles to 59,472 wildtype / 85,966 C5KO / 54,332 C5LKO.

Using Warp, all particles were then extracted into particles of 120 px side length with a pixel spacing of 3.386 Å. These datasets were then subjected to RELION 3D auto-refine using averages determined from the current particle orientations (low-pass filtered to 40 Å) as reference. Particles were automatically distributed into even and odd subsets during the RELION workflow. Subsequently, multi-particle refinement was performed in M v1.0.9 (*76*), simultaneously considering F-actin and branch junctions particles. Tilt-series were refined using image warp with a 9 × 6 grid and volume warp with a 4 × 6 × 2 × 10 grid as well as tilt-angle optimization. Particle poses were refined for one temporal sampling point. Particles were re-extracted from the refined tilt-series at a pixel spacing of 1.693 Å and a side length of 200 px and were again aligned with RELION 3D auto-refine, using the result of the previous iteration filtered to 40 Å as reference. Multi-particle refinement was then again performed in M, simultaneously considering F-actin and branch junction particles. Tilt-series were refined using image warp with a 9 × 6 grid and volume warp with a 4 × 6 × 2 × 10 grid, as well as tilt-angle optimization and frame alignment. Particle poses were refined for one temporal sampling point. The filtered and sharpened maps of the branch junctions were considered the final structures for their respective data sets. Resolution at the

0.143 FSC criterion was calculated with RELION post-process employing the halfmaps from M as input. The masks employed for FSC measurements are shown in (Figure S10). The difference map was calculated using TEMPy: DiffMap (*77*) as implemented in the CCP-EM software suite v1.5 (*78, 79*), employing low-pass filtering of both maps to 8.1 Å, a fractional difference cut-off of 0.4 and dust filtering.

### Statistical analysis

Statistical analyses were performed with Prism v9.0.2 (GraphPad Software). Differences were assessed by a non-parametric two-tailed Student’s t-test (two groups) or a Kruskal-Wallis test followed by a Dunn’s multiple comparison test with the respective wildtype as reference (more than two groups). We considered p-values smaller than 0.05 as significant. All cell biological experiments were repeated three times on independent samples. Quantifications from ultrastructural analysis stem from a single data acquisition per genotype. Sample numbers, medians, and p-values can be found in the figures and figure legends. (Figure S14) contains descriptive statistics, normality tests, and test statistics for all quantifications.

## Supplementary Figures

**Figure S1:**
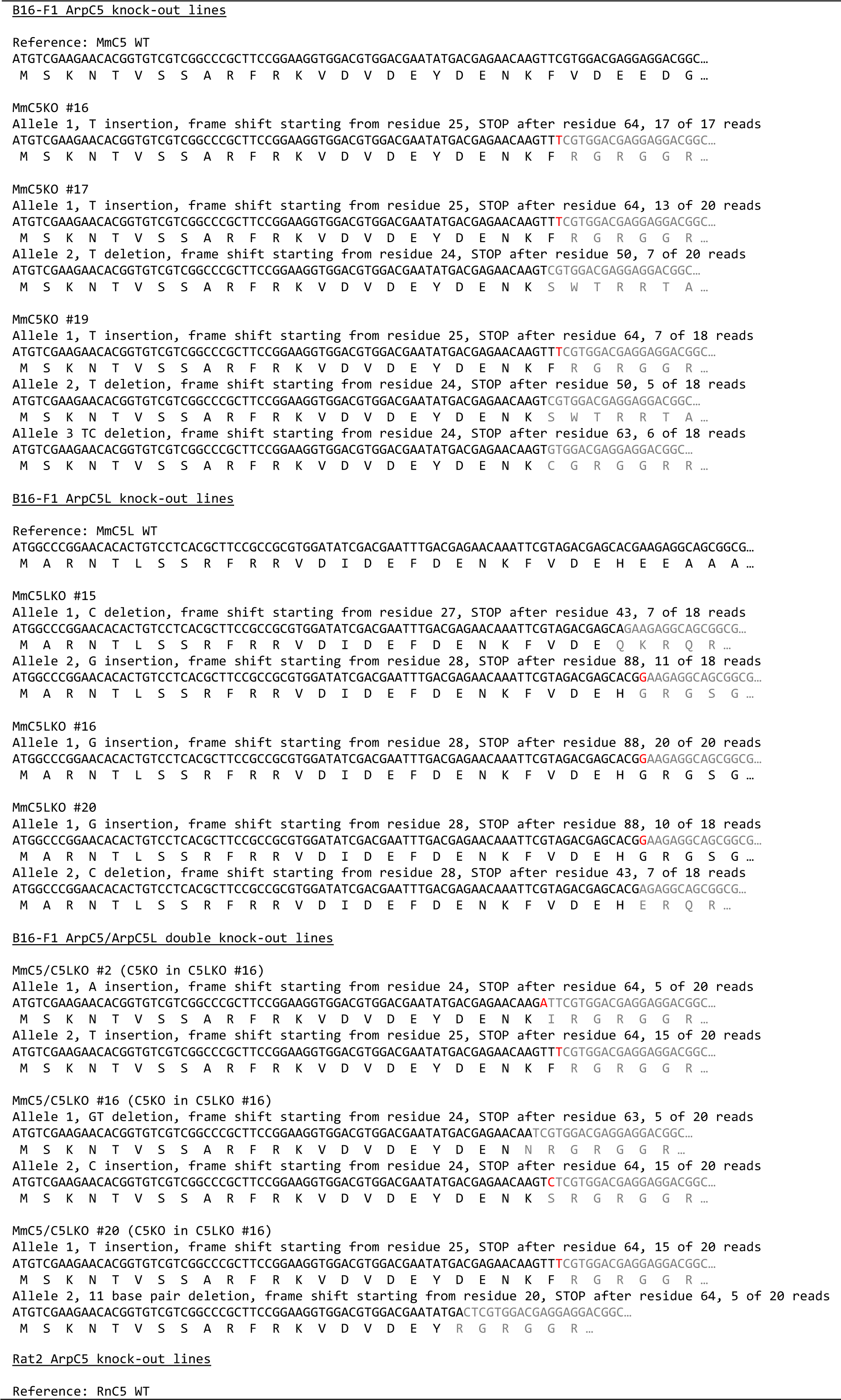
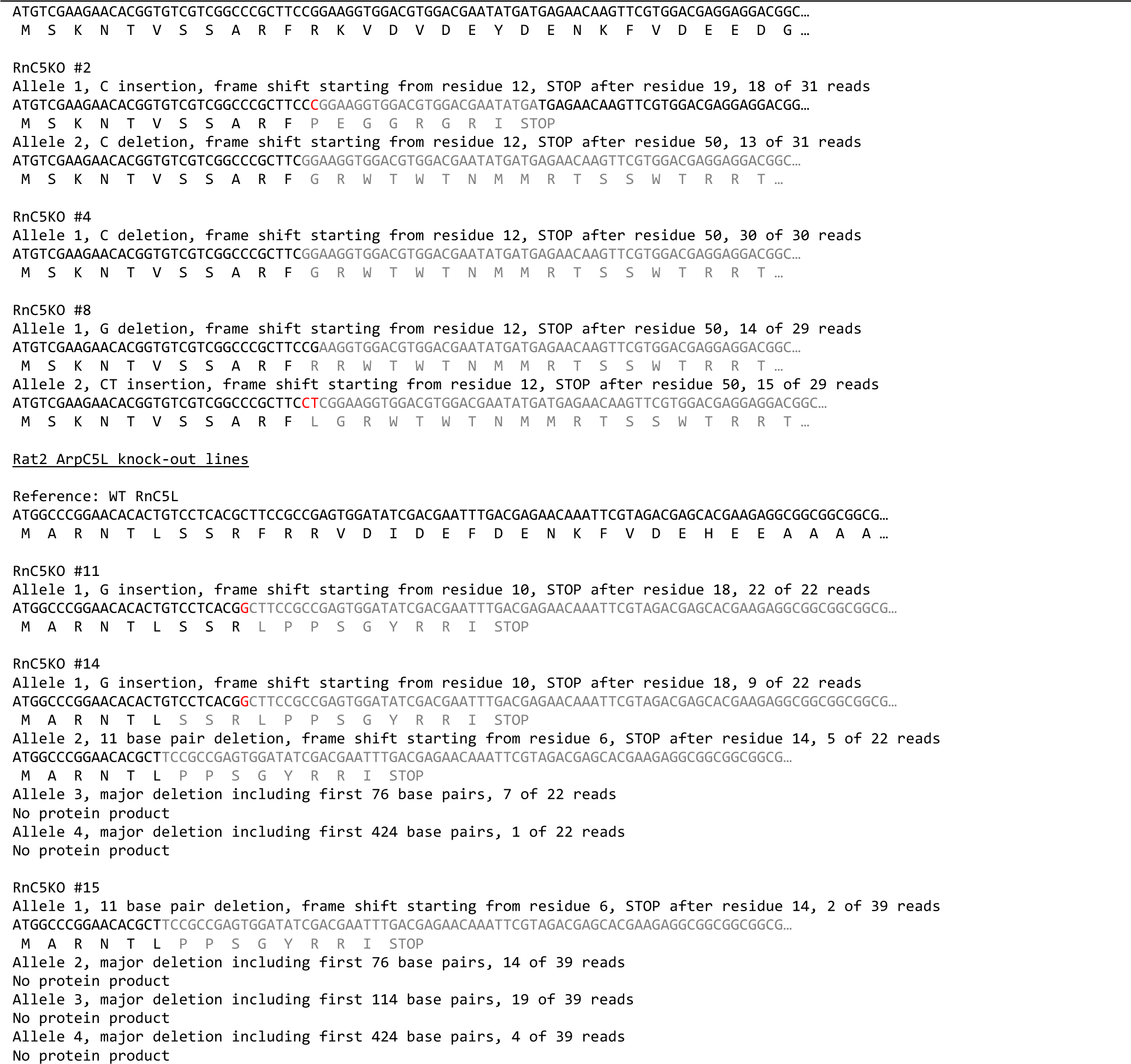
Sequencing results used to assess the ArpC5 and ArpC5L genotypes of cell lines used in this study.

**Figure S2:**
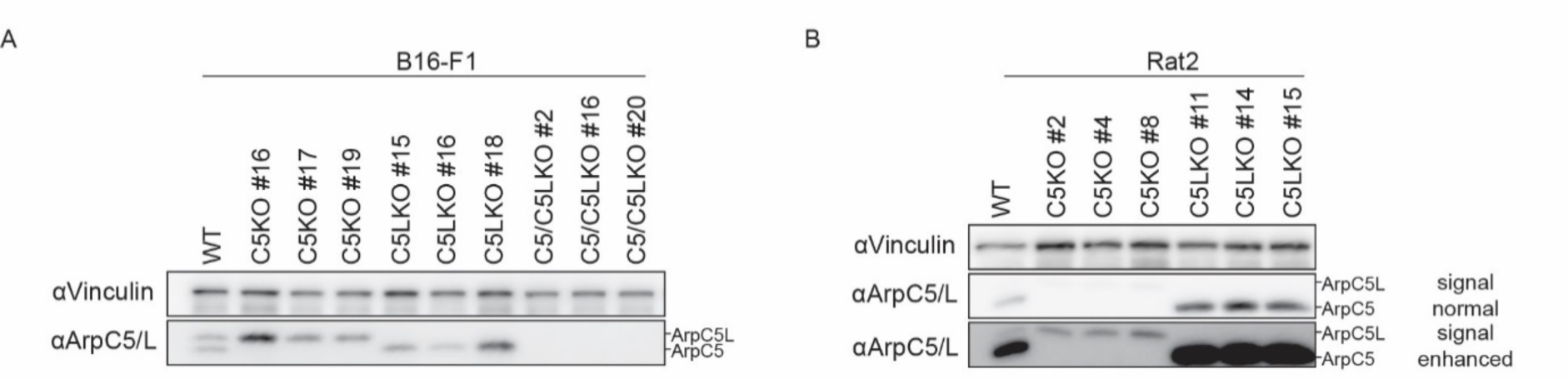
Western blot results used to assess the ArpC5 and ArpC5L genotypes of cell lines used in this study. **(A, B)** Representative Western blots showing levels of ArpC5 and ArpC5L proteins in three independent B16-F1 (A) and Rat2 (B) KO cell lines using a polyclonal antibody able to detect both isoforms. Double knockouts were only generated and analyzed in B16-F1 cells (A). Two visualizations of the same blot with different signal amplifications are shown for the Rat2 cells to allow visualization of both isoforms (B). Vinculin was used as loading control.

**Figure S3:**
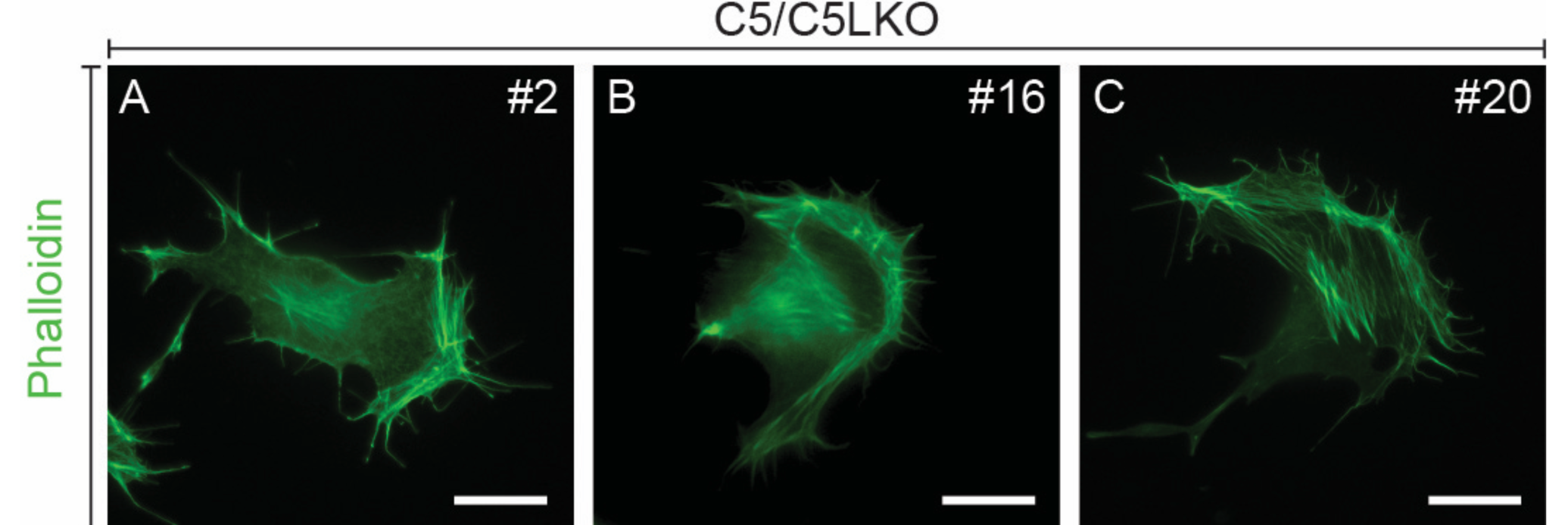
C5/C5LKO cells do not exhibit any lamellipodia. **(A-C)** Representative epifluorescence micrographs of B16-F1 cells from three different C5/C5LKO lines visualizing the actin cytoskeleton using fluorescent phalloidin. All scale bars, 20µm.

**Figure S4:**
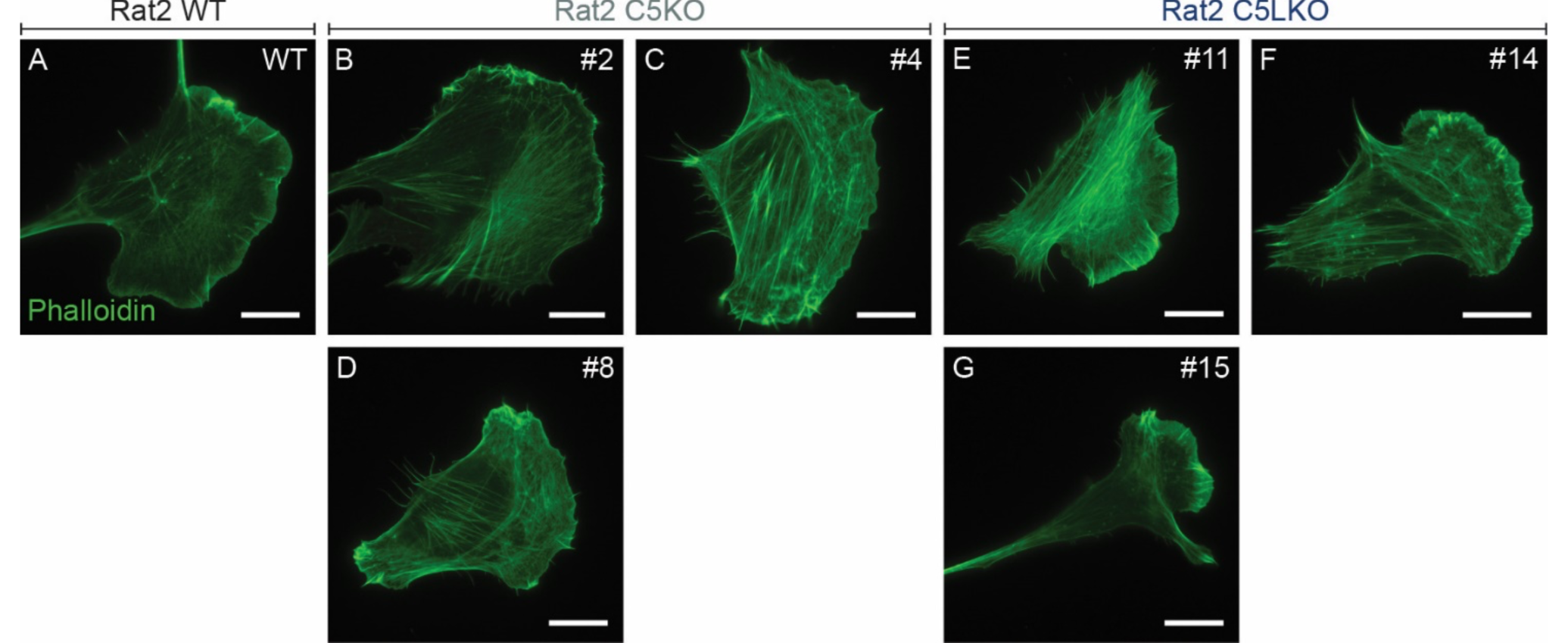
Rat2 C5KO and C5LKO cells recapitulate the lamellipodia phenotypes observed in the respective B16 knockout lines. (A-G) Representative epifluorescence micrographs of Rat2 wildtype (A), C5KO (B-D), and C5LKO (E-G) cells visualizing the actin cytoskeleton using fluorescent phalloidin. All scale bars, 20μm.

**Figure S5:**
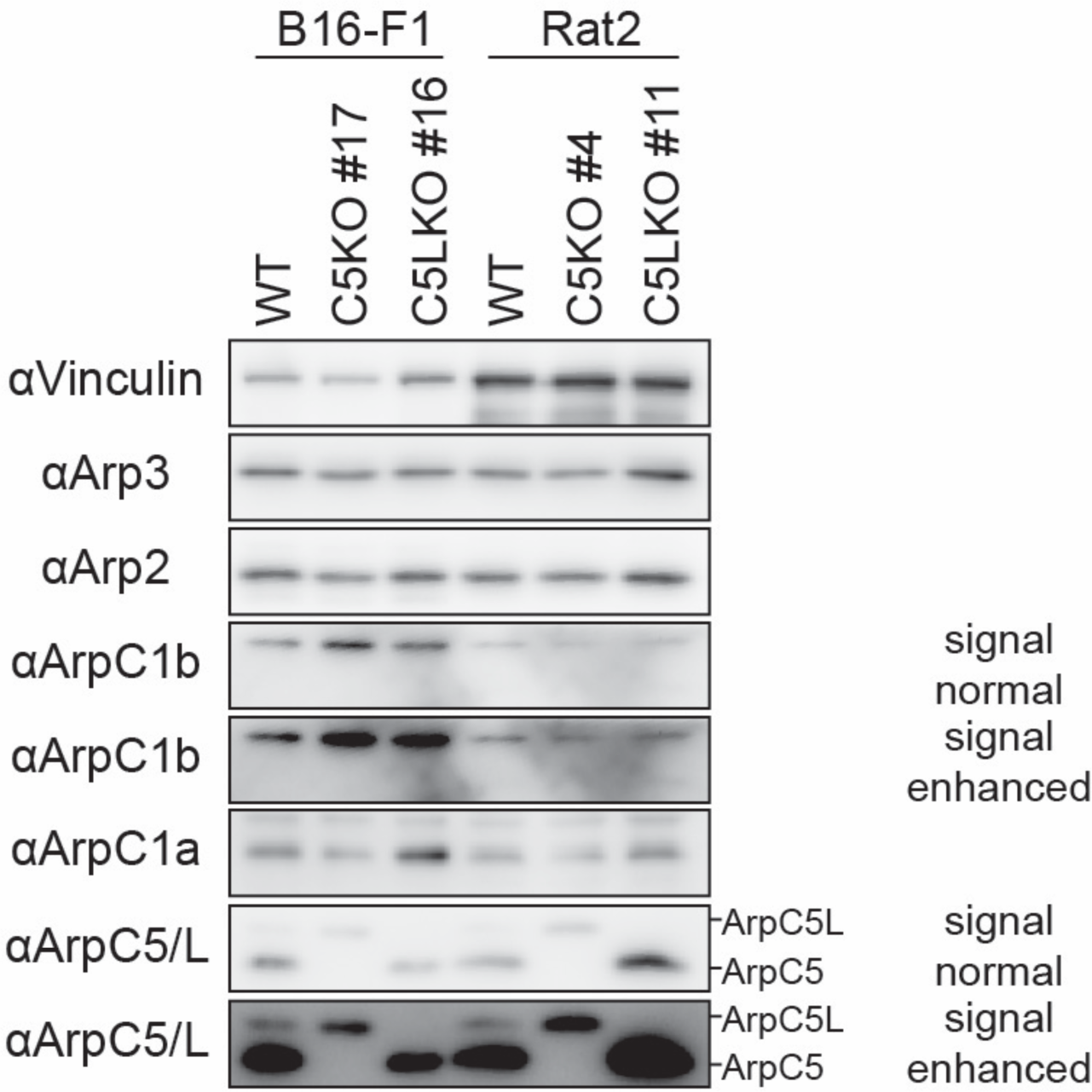
Expression levels of other Arp2/3 subunits. Western blot detecting Arp3, Arp2, ArpC1b, and ArpC1a confirms that abundance of these proteins is not severely reduced in B16-F1 and Rat2 C5KO and C5LKO cells compared to their respective wildtype counterparts. Two versions with different signal amplifications are shown for the ArpC1b and the polyclonal ArpC5/ArpC5L antibody to allow visualization in both cell types on the same membrane. Vinculin was used as loading control. Polyclonal ArpC5/ArpC5L antibody was employed to verify the sample genotypes.

**Figure S6:**
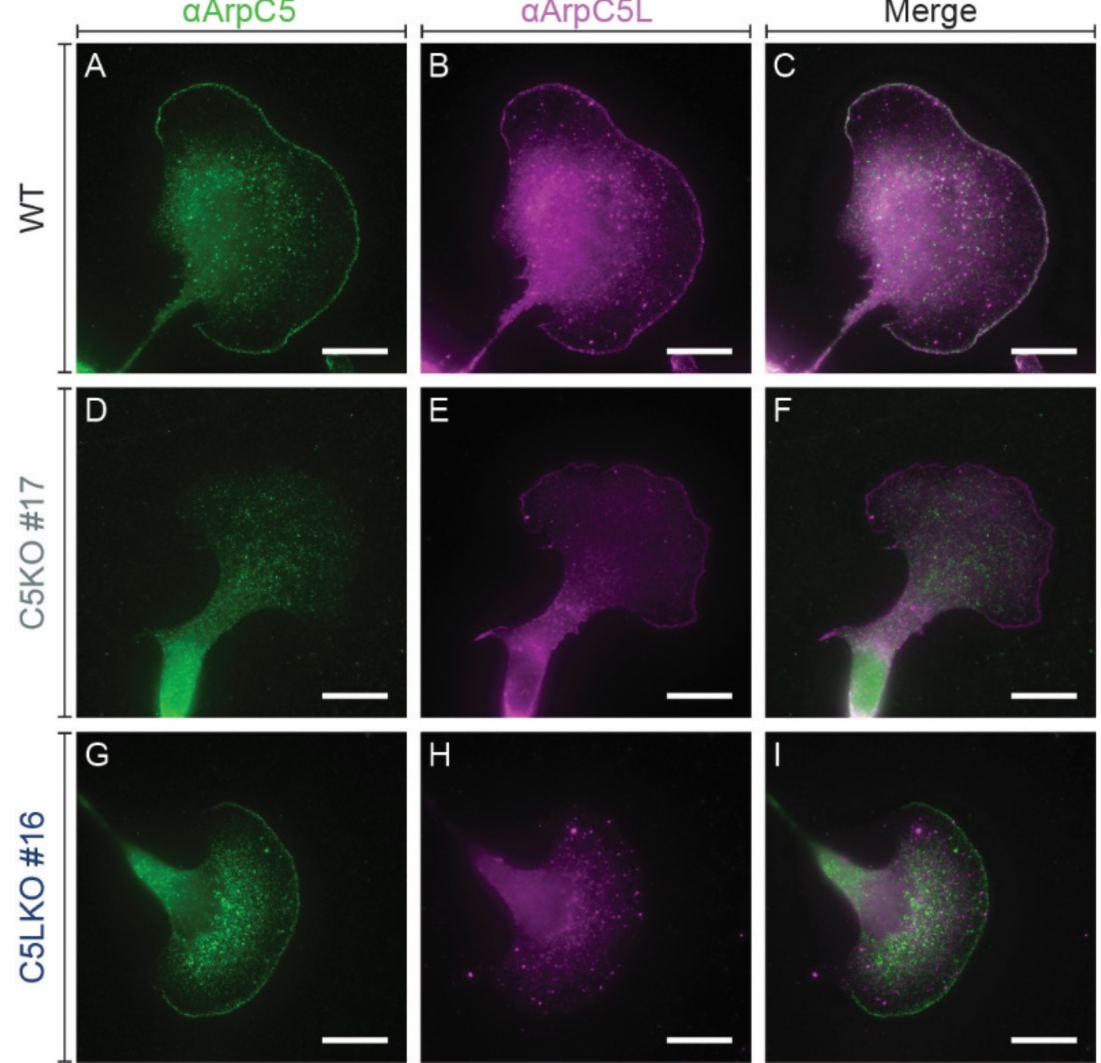
Testing monoclonal antiArpC5 and antiArpC5L antibodies for isoform-specificity. **(A-I)** Representative epifluorescence micrographs of B16-F1 wildtype (A-C), C5KO (D-F), and C5LKO (G-I) cells visualizing the presence/absence and the localization of ArpC5 (A, D, G) and ArpC5L (B, E, I) via immunofluorescence. Merges are shown in (C, F, I). All scale bars, 20µm.

**Figure S7:**
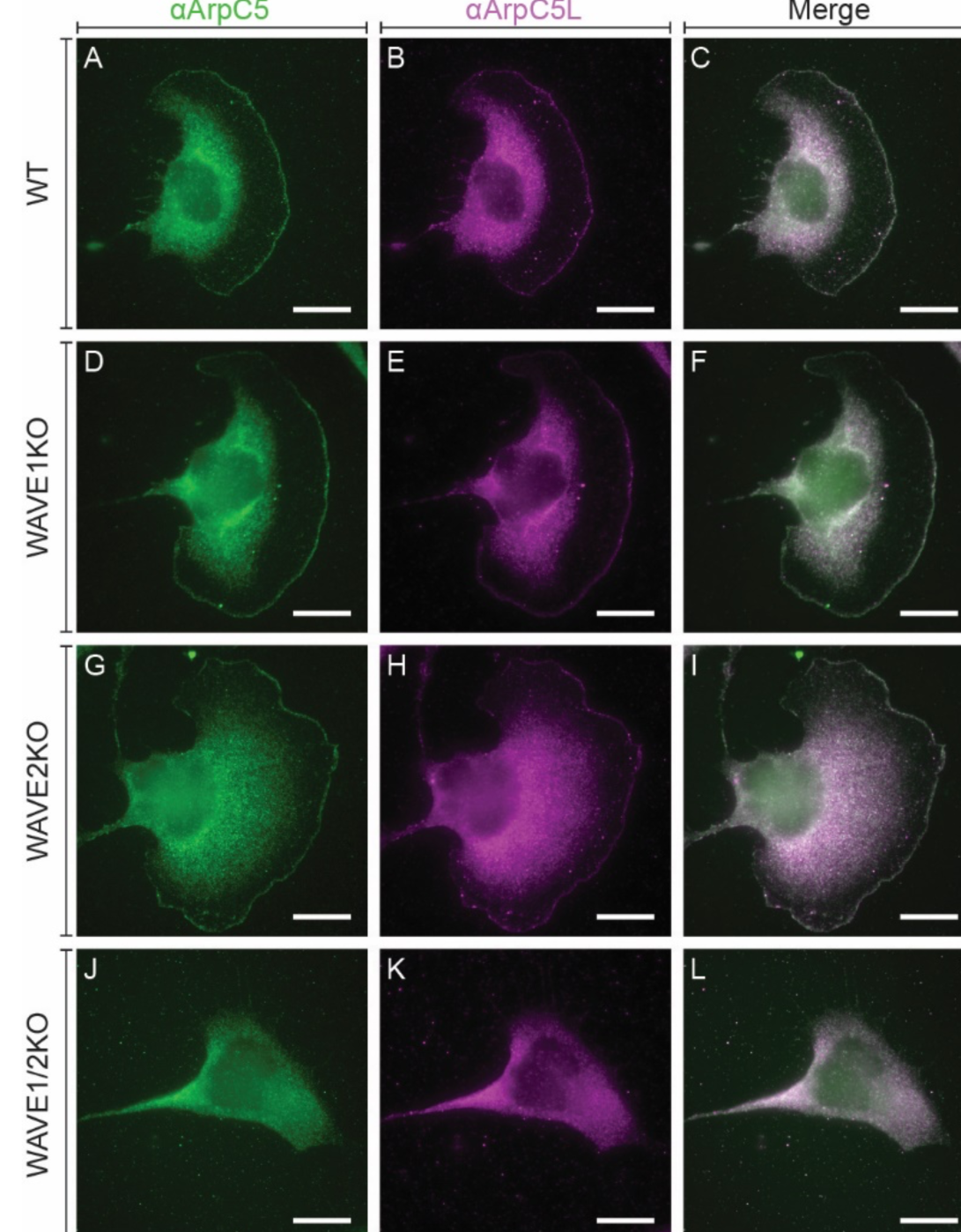
ArpC5 isoforms are not recruited by specific WAVE isoforms. **(A-L)** Representative epifluorescence micrographs of B16-F1 wildtype (A-C), WAVE1KO (D-F), WAVE2KO (G-I), and WAVE1/2KO (J-L) cells visualizing the presence/absence and the localization of ArpC5 (A, D, G, J), and ArpC5L (B, E, I, K) via immunofluorescence. Overlays of the signals are shown in (C, F, I, L). All scale bars, 20µm.

**Figure S8:**
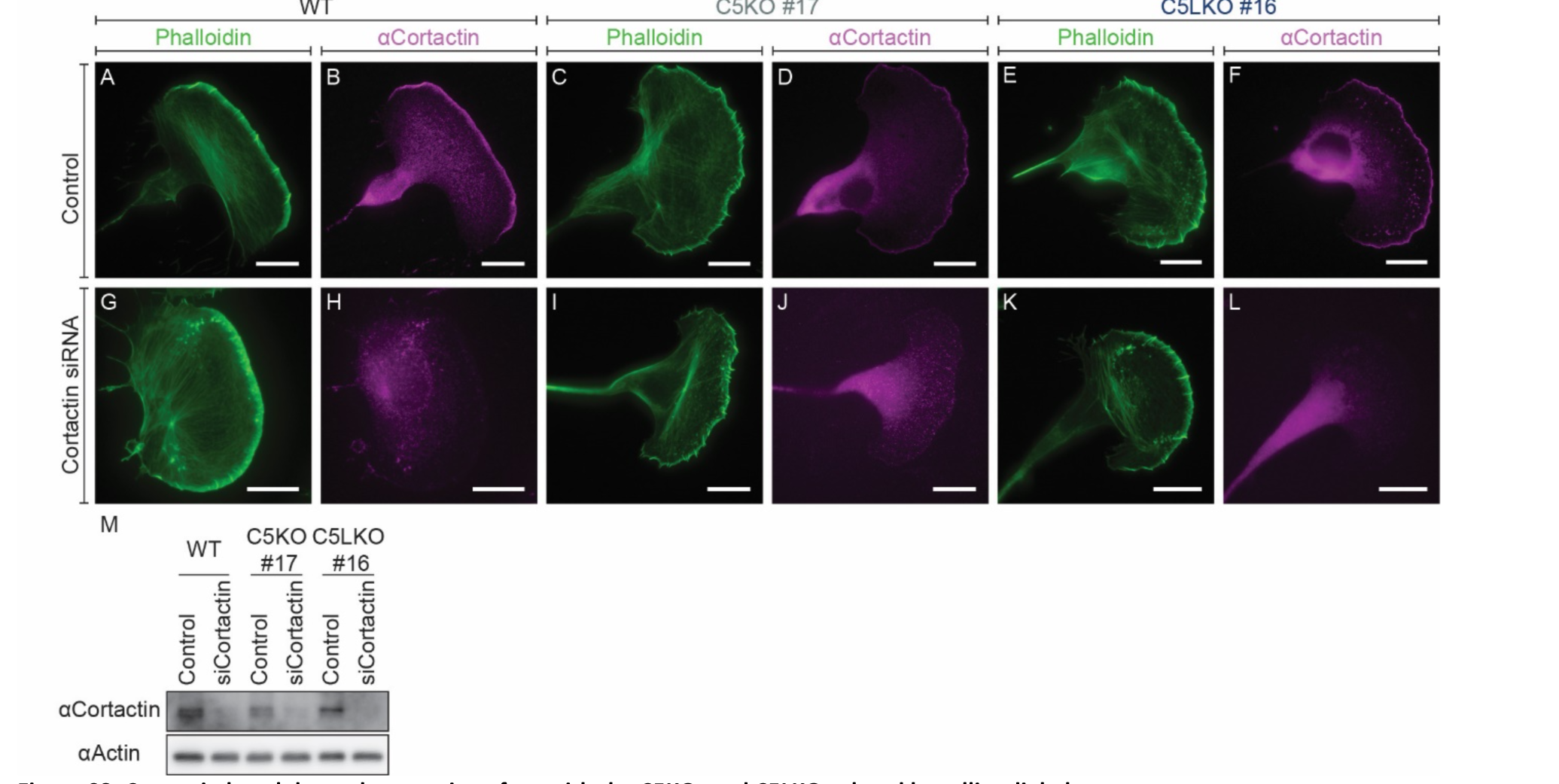
Cortactin knockdown does not interfere with the C5KO- and C5LKO-related lamellipodial phenotypes. **(A-L)** Representative epifluorescence micrographs of B16-F1 wildtype (A, B, G, H), C5KO (C, D, I, J), and C5LKO (E, F, K, L) cells after their transfection with control siRNA (A-F) or siRNA targeting Cortactin (G-L). The actin cytoskeleton is visualized by fluorescent phalloidin (A, C, E, G, I, K) and the presence/absence and the localization of Cortactin (B, D, F, H, J, L) via immunofluorescence. **(M)** Western blot analysis of a subpopulation of cells transfected together with the cells shown in (A-L) to verify Cortactin knockdown on a population scale. All scale bars, 20µm.

**Figure S9:**
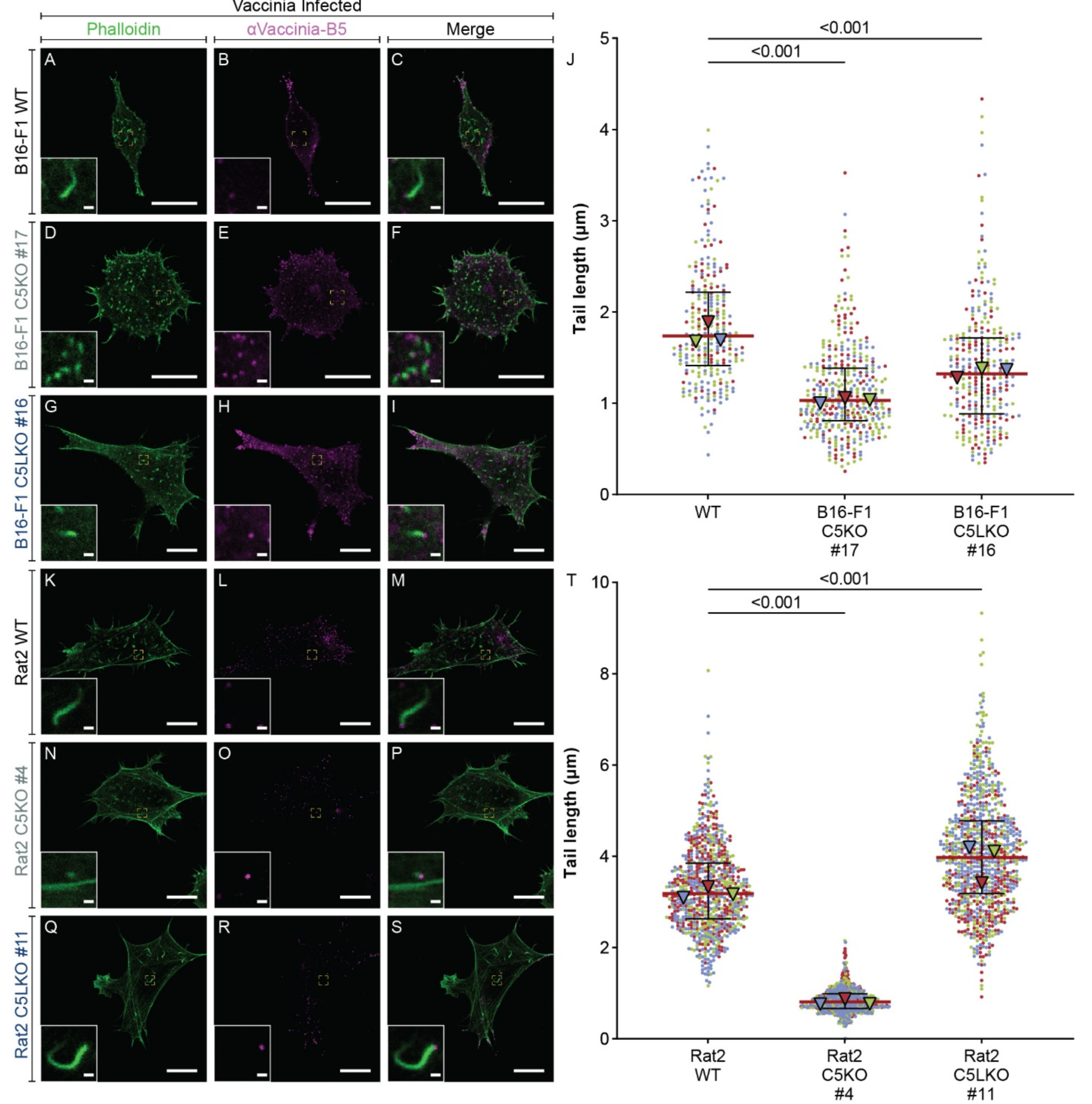
Effects of ArpC5 isoform depletion on Vaccinia virus tails. **(A-I)** Representative epifluorescence micrographs of Vaccinia virus-infected B16-F1 wildtype (A, B, C), C5KO (D, E, F), and C5LKO (G, H, I) cells visualizing the actin cytoskeleton and thus the Vaccinia tails using fluorescent phalloidin (A, D, G,) and the viruses using Vaccinia-B5 antibody (B, E, H). Overlays of the signals are shown in (C, F, I). Inlays show individual viruses and their tails. Dashed yellow rectangles indicate their position in the micrographs. **(J)** Vaccinia tail lengths were measured in infected B16-F1 wildtype, C5KO, and C5LKO cells. Kruskal-Wallis test combined with Dunn’s multiple comparison test on pooled data from 3 independent experiments, n=263, 391, 292, p values shown in the chart. Black lines indicate overall medians (1.739µm, 1.033µm, 1.,324µm) and quartile ranges. **(K-S)** Representative epifluorescence micrographs of Vaccinia virus-infected Rat2 wildtype (K, L, M), C5KO (N, O, P), and C5LKO (Q, R, S) cells visualizing the actin cytoskeleton and thus Vaccinia tails using the same staining as in (A-I). Inlays show individual viruses and their tails. Their position in the micrographs is indicated by dashed yellow rectangles. **(T)** Vaccinia tail lengths were measured in infected Rat2 wildtype, C5KO, and C5LKO cells. Kruskal-Wallis test combined with Dunn’s multiple comparison test on pooled data from 3 independent experiments, n=738, 728, 846, p values shown in the chart. Black lines indicate overall medians (3.183µm, 0.8095µm, 3.974µm) and quartile ranges. Scale bars, 20µm in standard panels and 1µm in inlays. Data points are color-coded according to the individual experiments. Triangles indicate the medians of the respective experiments.

**Figure S10:**
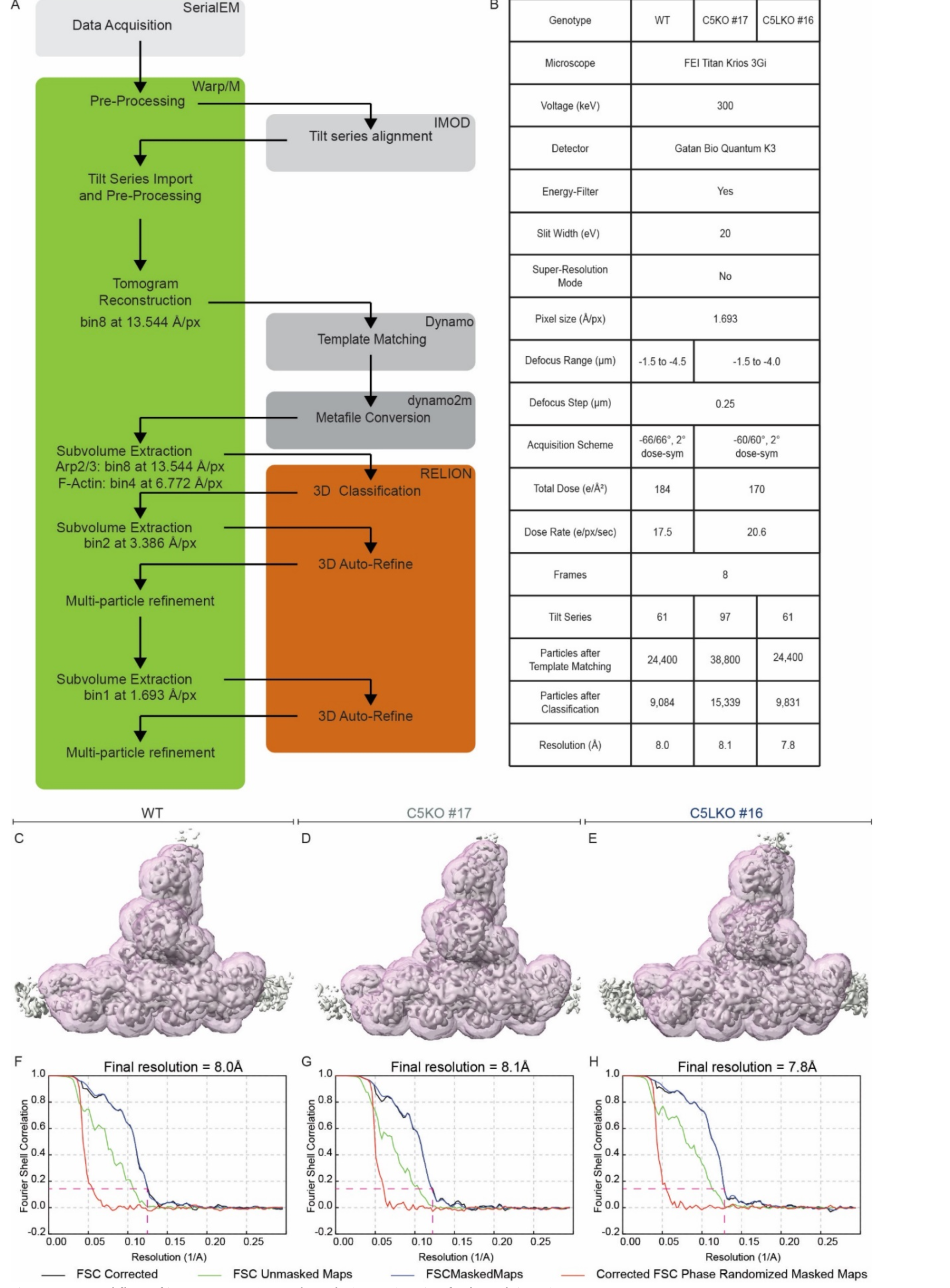
Workflow of image processing and resolution estimation for branch junction structures. **(A)** Flow chart indicating the data processing steps involved in generating the three structures of the active Arp2/3 complexes within respective branch junctions. Colored boxes indicate the use of specific software packages. **(B)** Data acquisition and image processing parameters. **(C-E)** Isosurface representation of the actin filament Arp2/3 complex branch junction in B16-F1 wildtype, C5KO, and C5LKO cells (shown in solid grey) filtered to their respective resolution determined by gold standard FSC calculation. Masks (transparent purple) used for FSC calculations are superimposed onto the respective structures. **(F-H)** FSC curves with indicated resolution cut-offs according to the 0.143 criterion.

**Figure S11:**
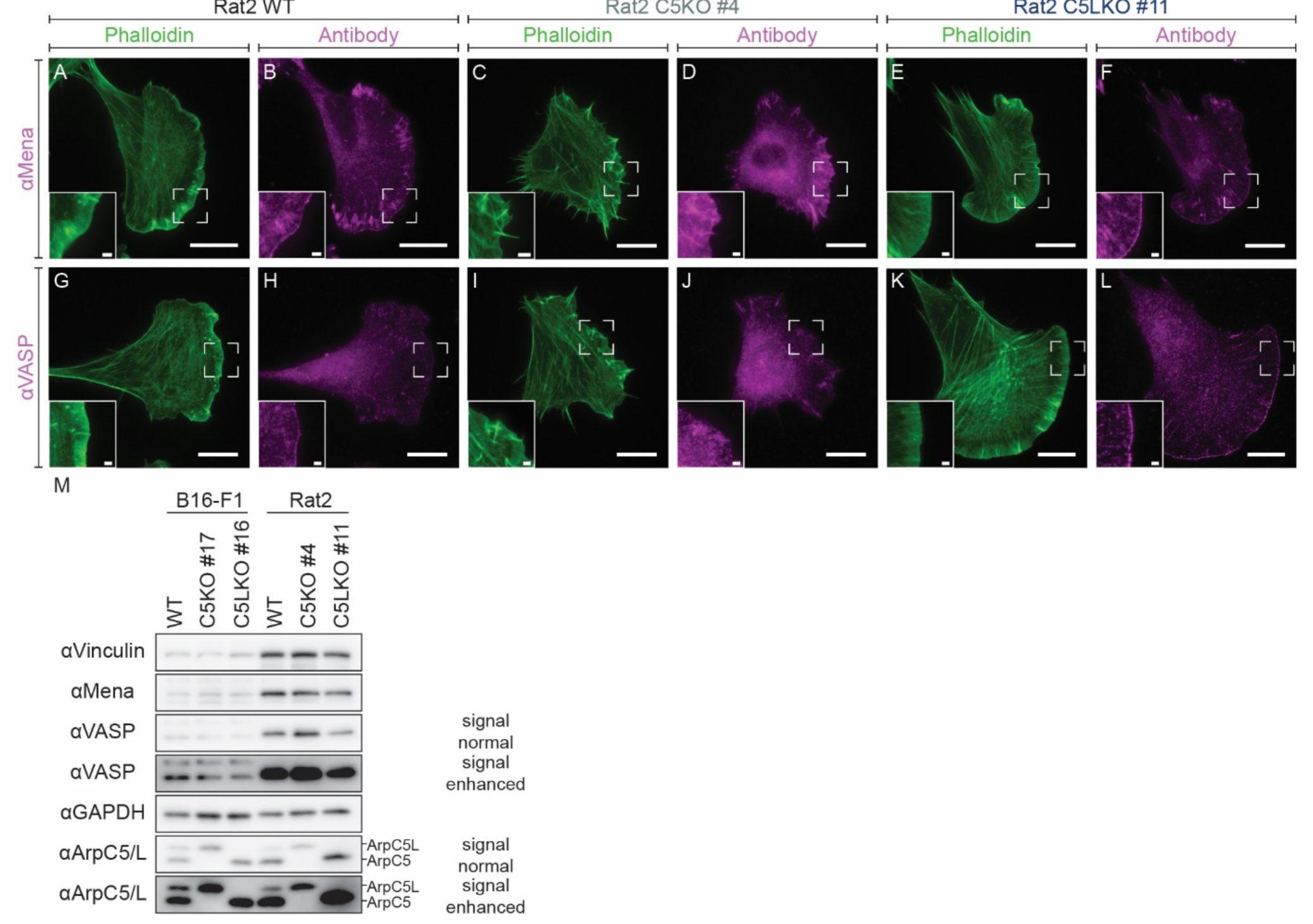
Rat2 C5KO and C5LKO cells recapitulate the Mena/VASP recruitment phenotypes observed in the respective B16 knockout lines. **(A-L)** Representative epifluorescence micrographs of Rat2 wildtype (A, B, G, H), C5KO (C, D, I, J), and C5LKO (E, F, K, L) cells visualizing the actin cytoskeleton using fluorescent phalloidin (A, C, E, G, I, K) and the localization of Mena (B, D, F) and VASP (H, J, L) via immunofluorescence. Inlays show magnified areas indicated in the respective panels. **(M)** Western blot detecting Mena and VASP confirms that their absence at the leading edge in B16- F1 and Rat2 C5KO cells is due to reduced recruitment and not due to reductions at the protein level. Two versions with different signal amplifications are shown for the VASP and the polyclonal ArpC5/ArpC5L antibodies to allow visualization in both cell types on the same membrane. Vinculin and GAPDH were used as loading controls. Polyclonal ArpC5/ArpC5L antibody was employed to verify the sample genotypes. Scale bars, 20µm in standard panels and 2µm in inlays, n.s. = not significant.

**Figure S12:**
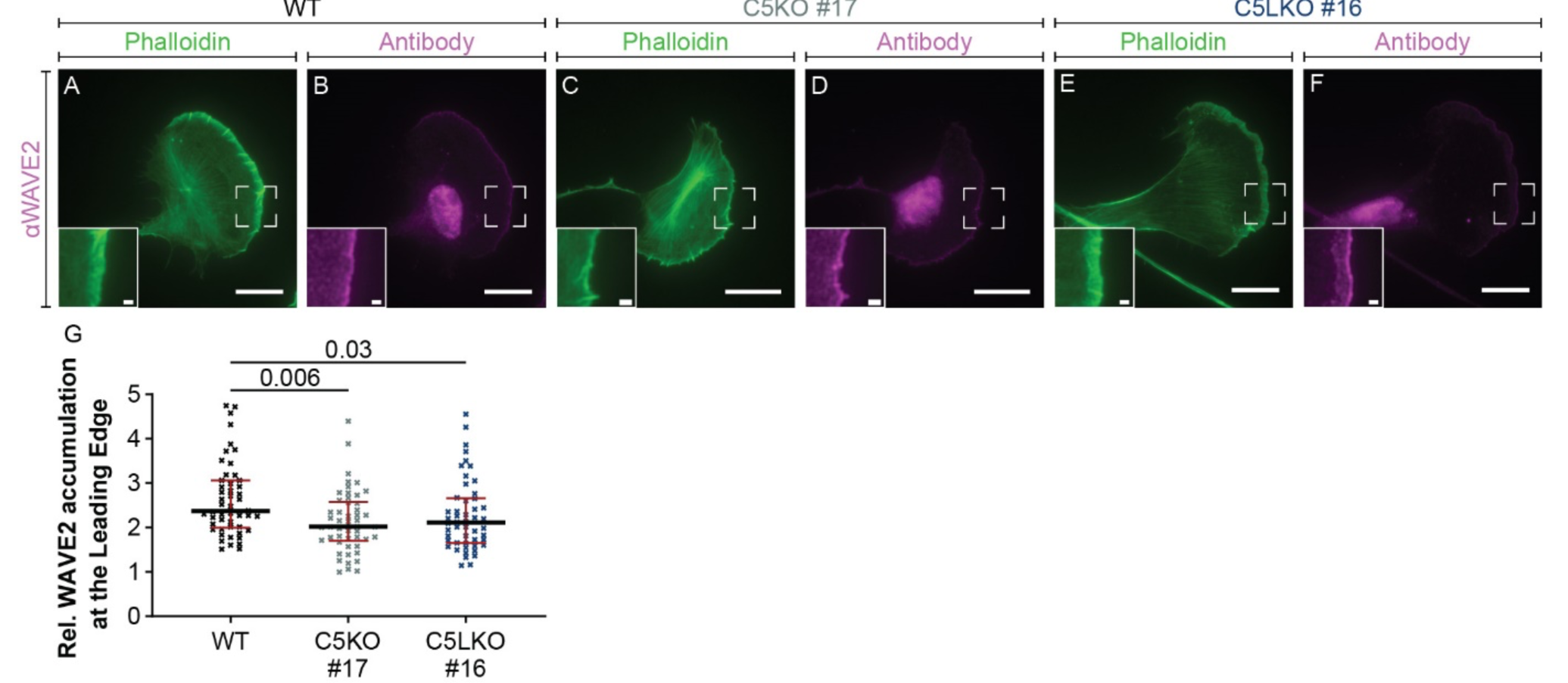
Depletion of either ArpC5 isoform marginally reduces WAVE2 accumulation at the leading edge of protruding lamellipodia. **(A-L)** Representative epifluorescence micrographs of B16-F1 wildtype (A, B), C5KO (C, D), and C5LKO (E, F) cells visualizing the actin cytoskeleton using fluorescent phalloidin (A, C, E) and the localization of WAVE2 (B, D, F) by immunofluorescence. Inlays show magnified areas indicated in the respective panels. **(G)** Quantitative analysis of the relative accumulation of WAVE2 at the leading edges of B16-F1 wildtype, C5KO, and C5LKO cells. Kruskal-Wallis test combined with Dunn’s multiple comparison test, n=50 cells for each experimental conditions, for both Mena and VASP, p values shown in the chart. Red lines indicate medians (2.367, 2.019, 2.110) and black lines quartile ranges. Scale bars, 20µm in standard panels and 1µm in inlays.

**Figure S13:**
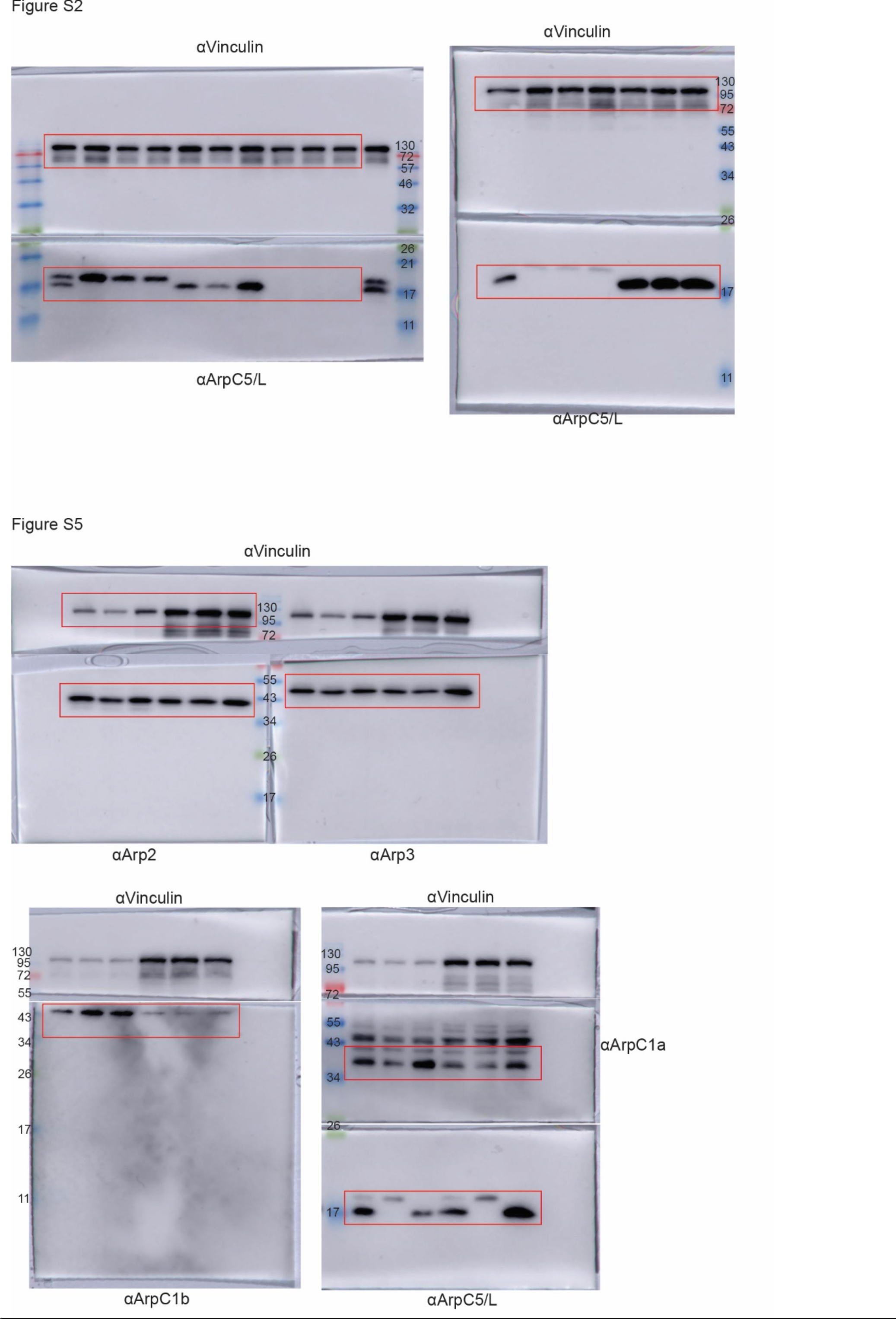
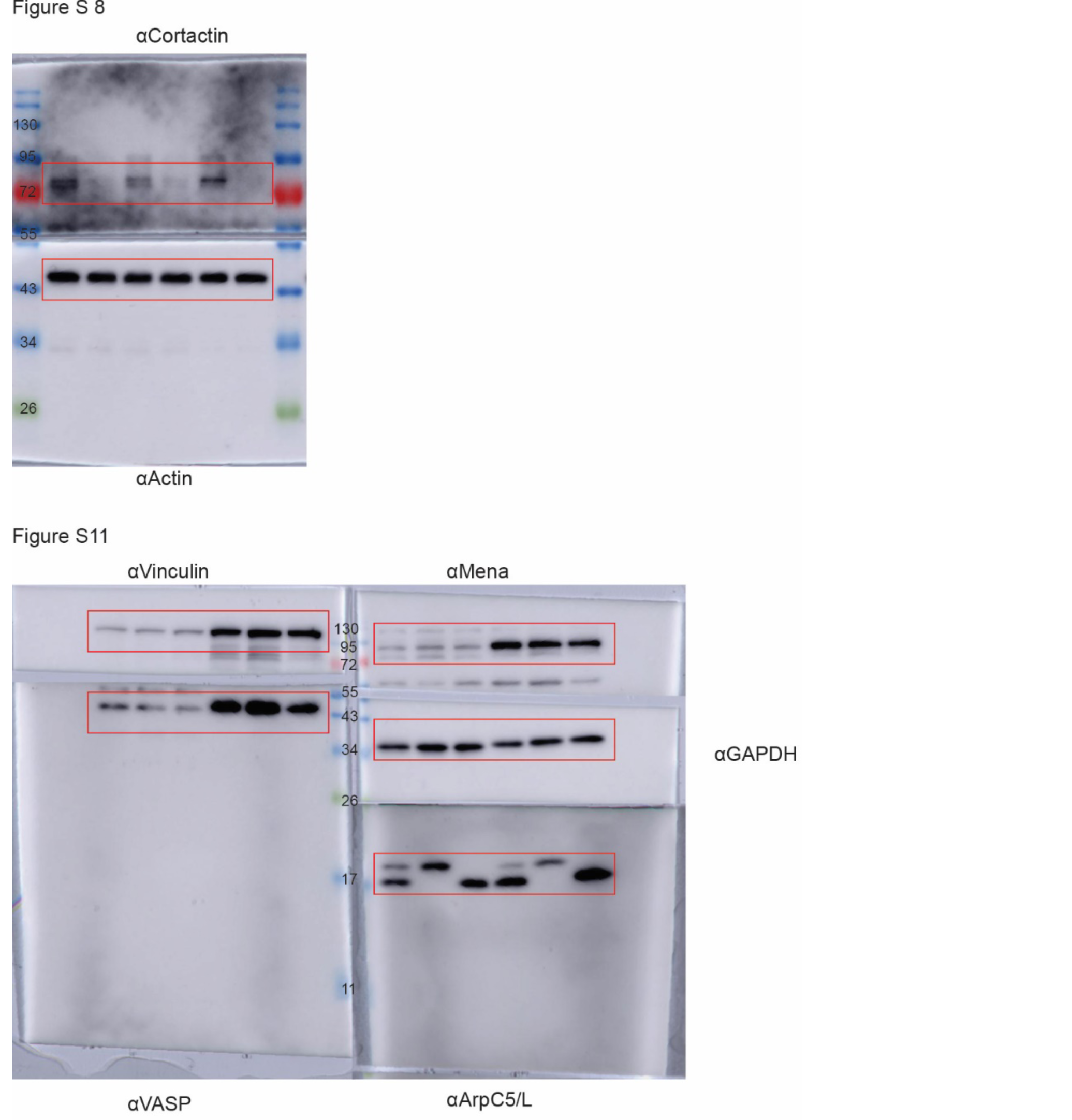
Full Western blots. Chemiluminescence signals and color images of the PVDF membranes are overlaid. Corresponding figures, employed antibodies, and identity of the marker bands are annotated next and on the blots, respectively. Red rectangles highlight areas shown in the corresponding figures.

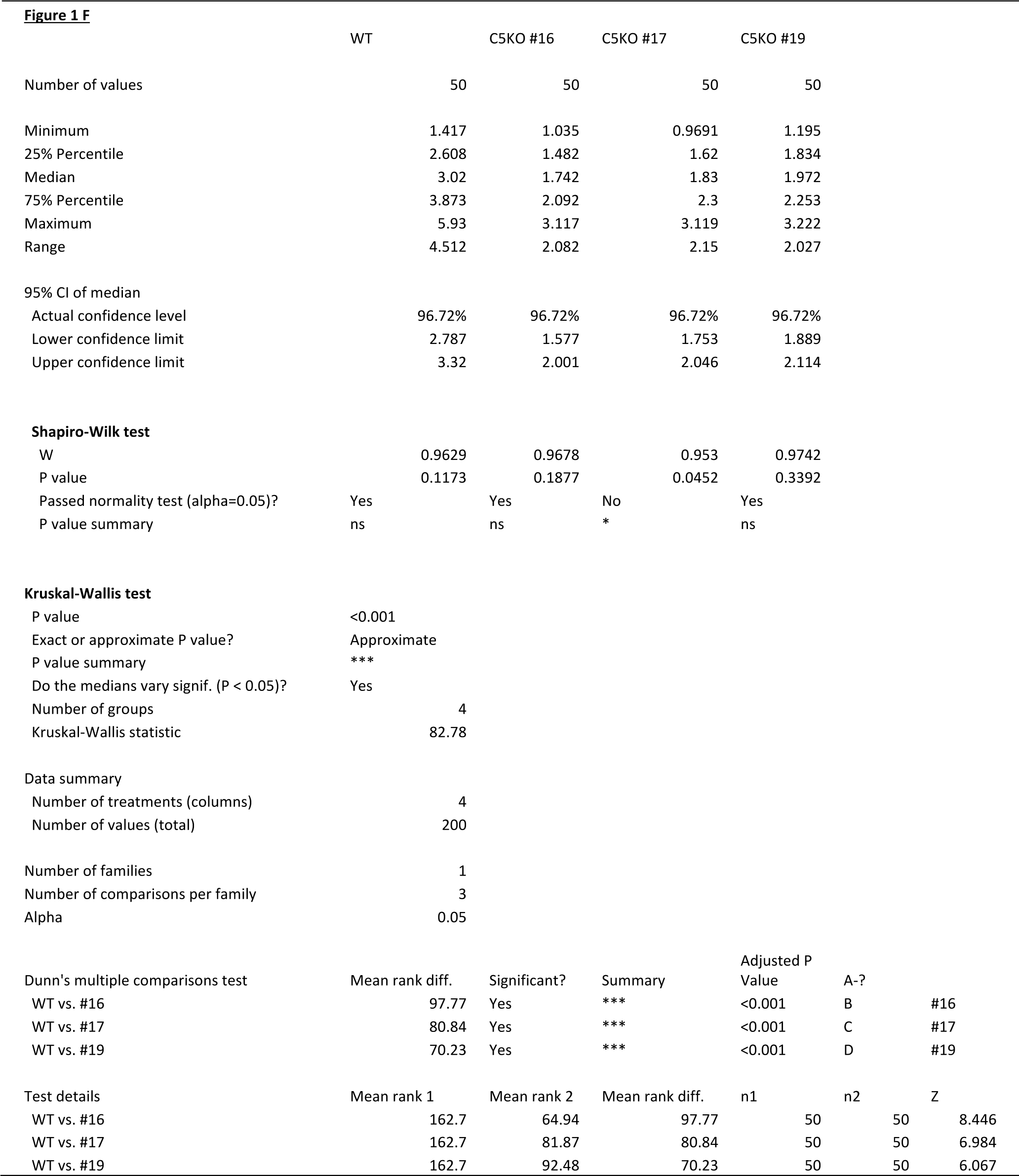
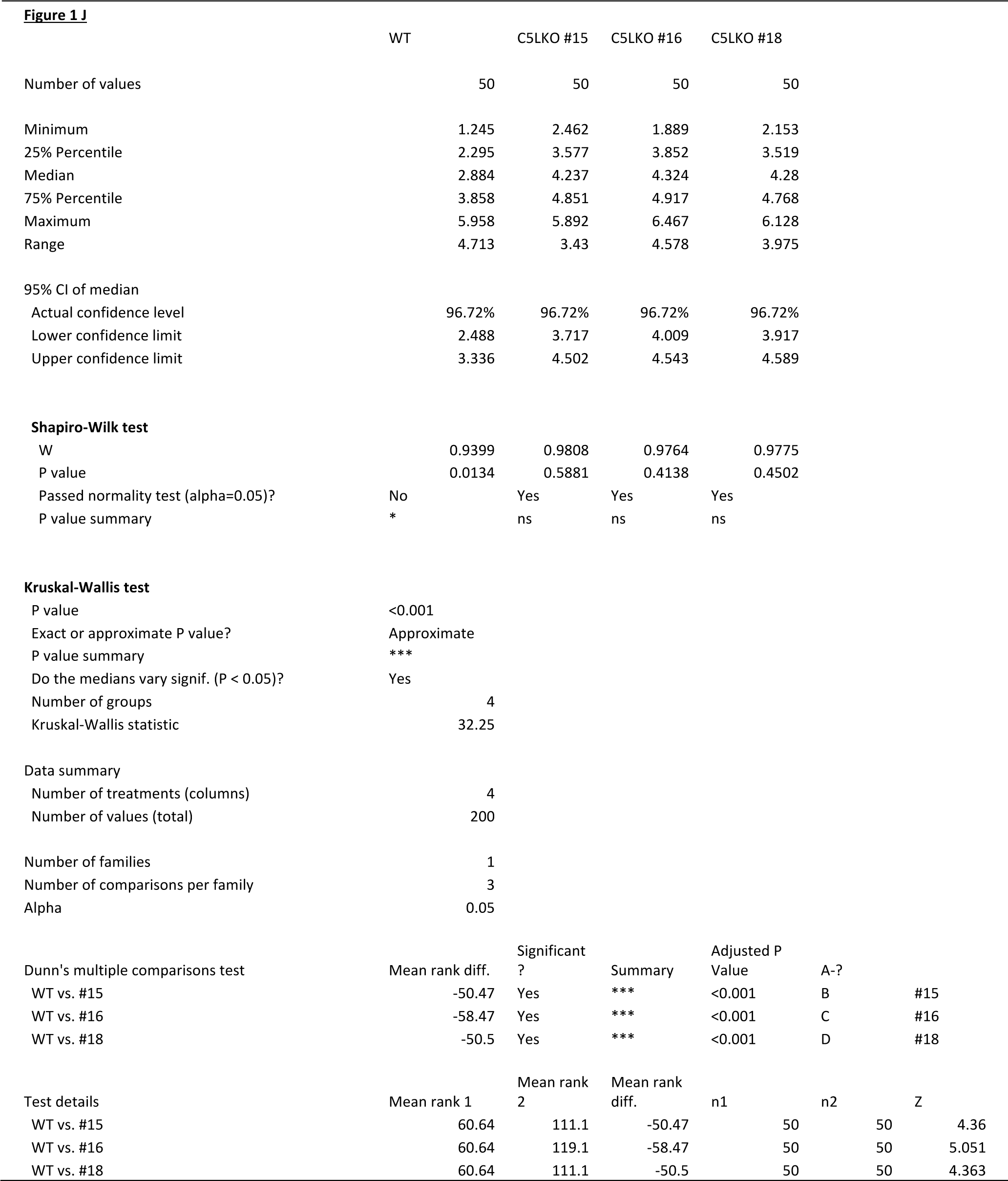
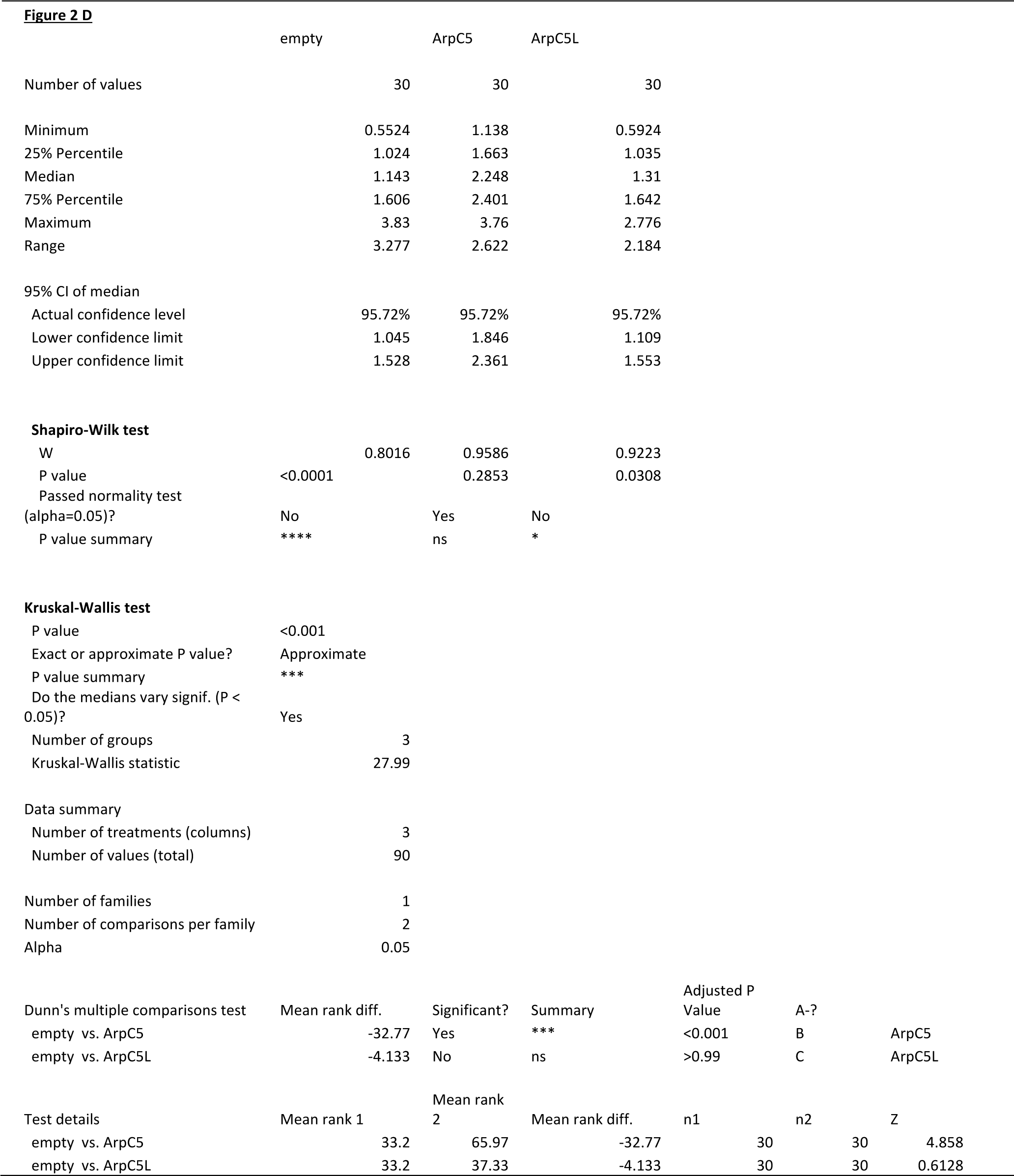
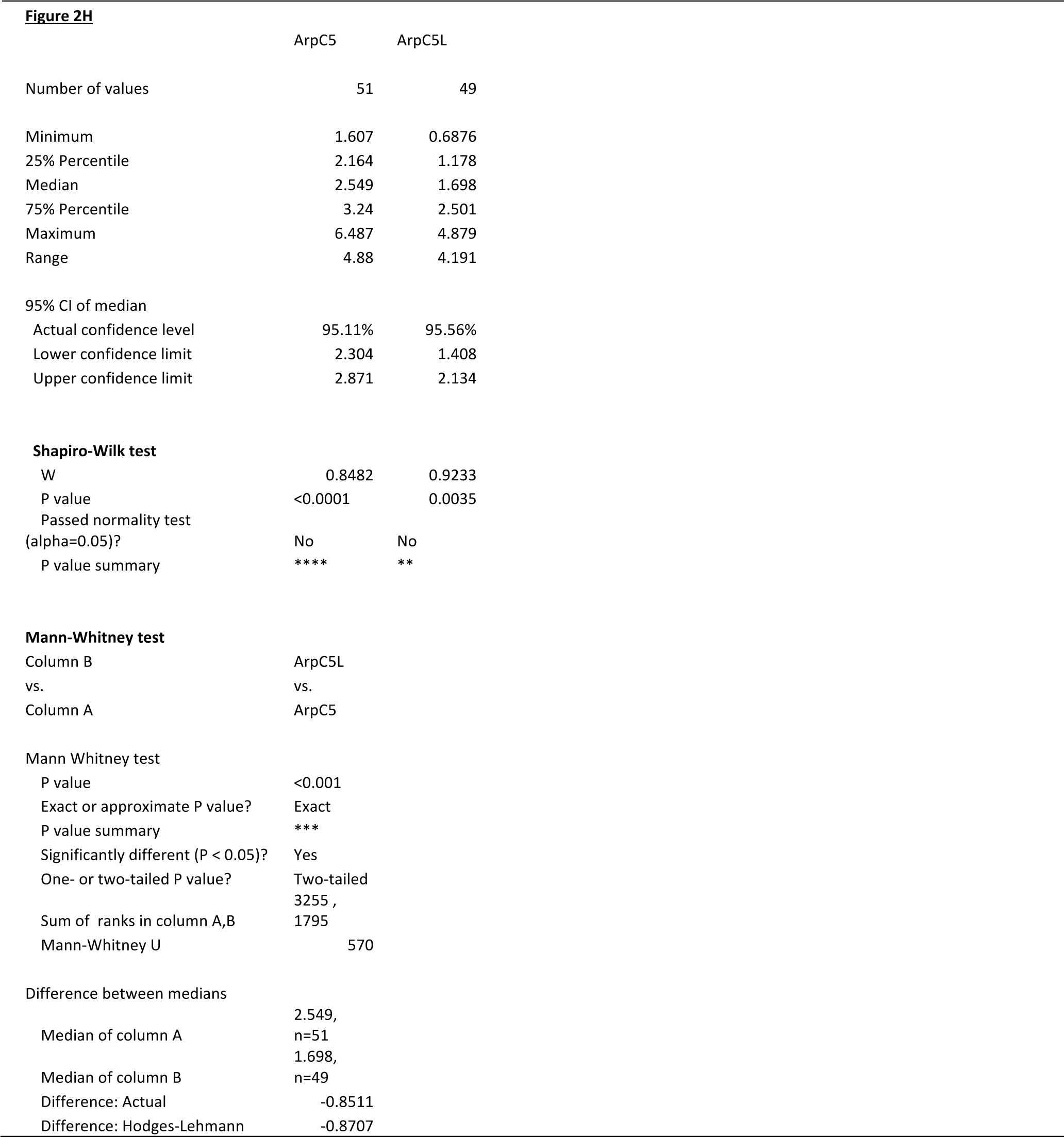
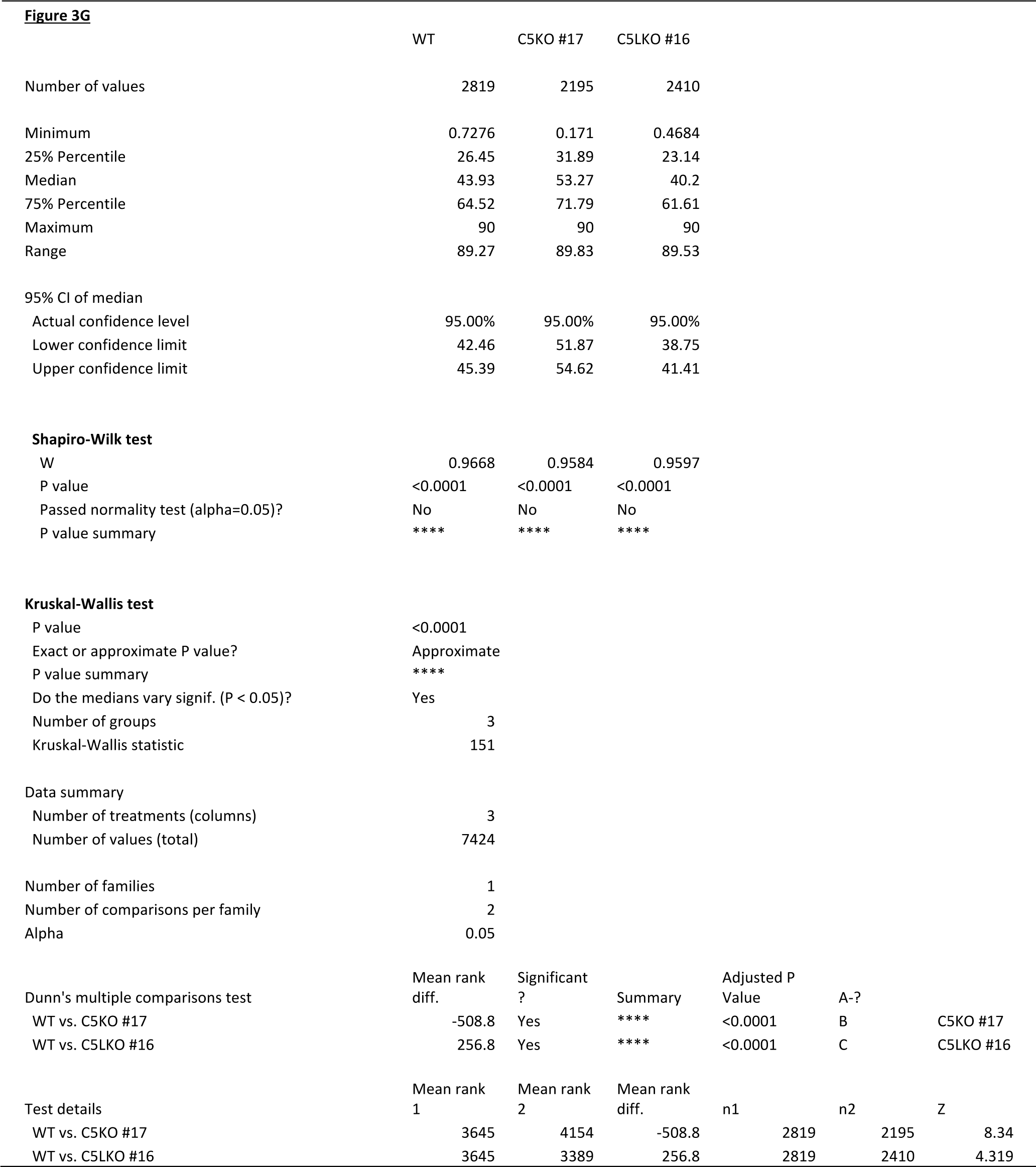
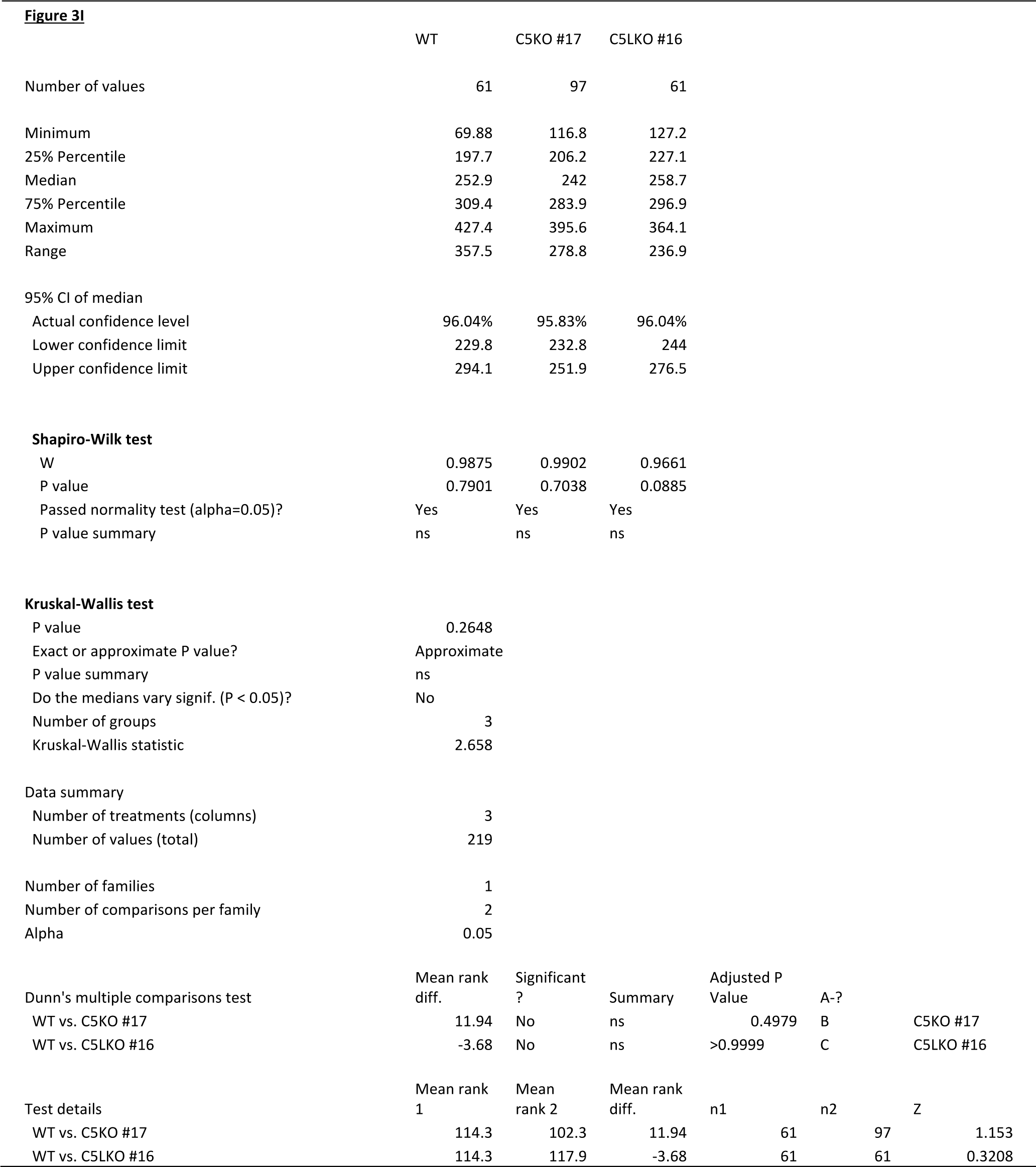
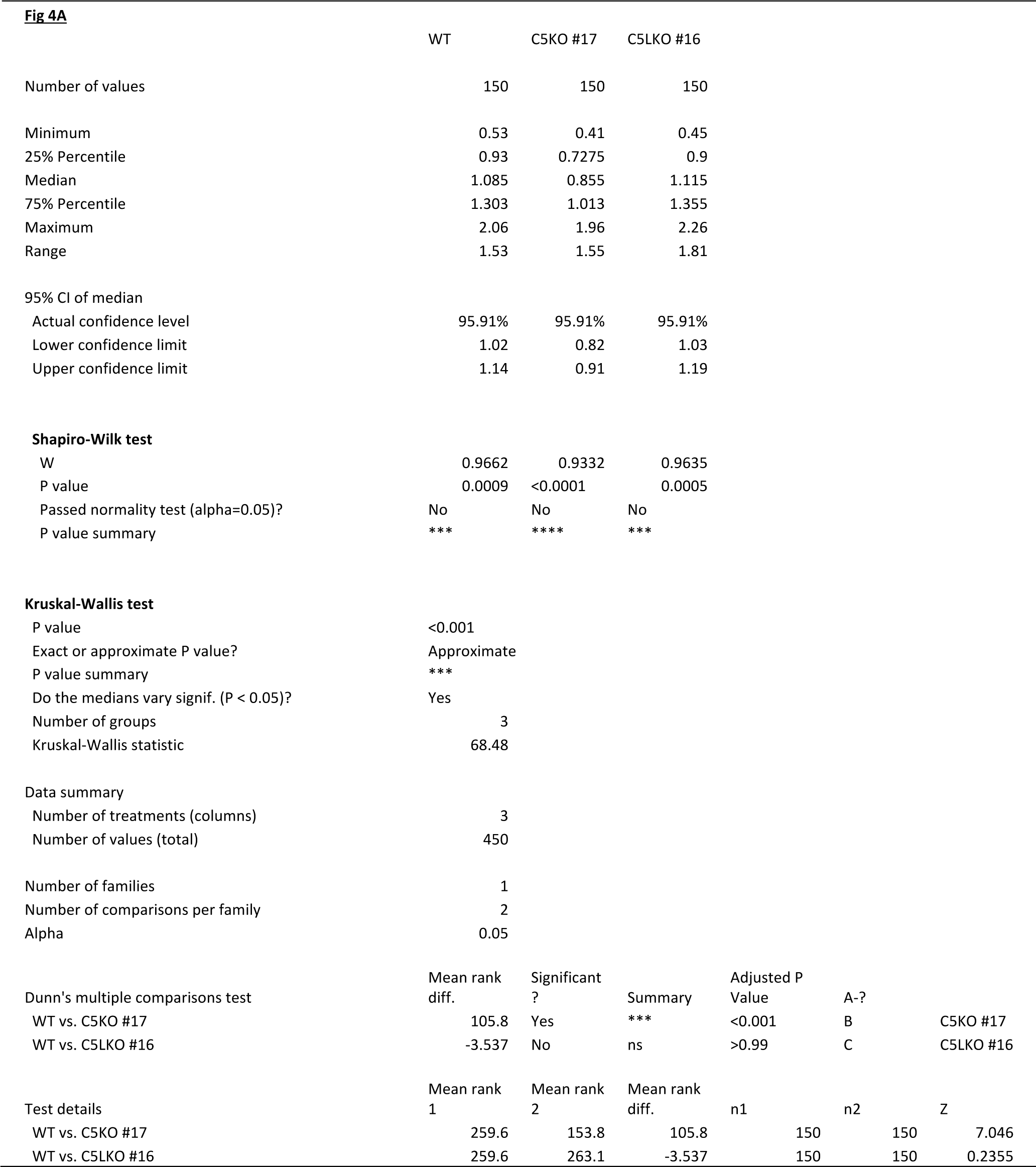
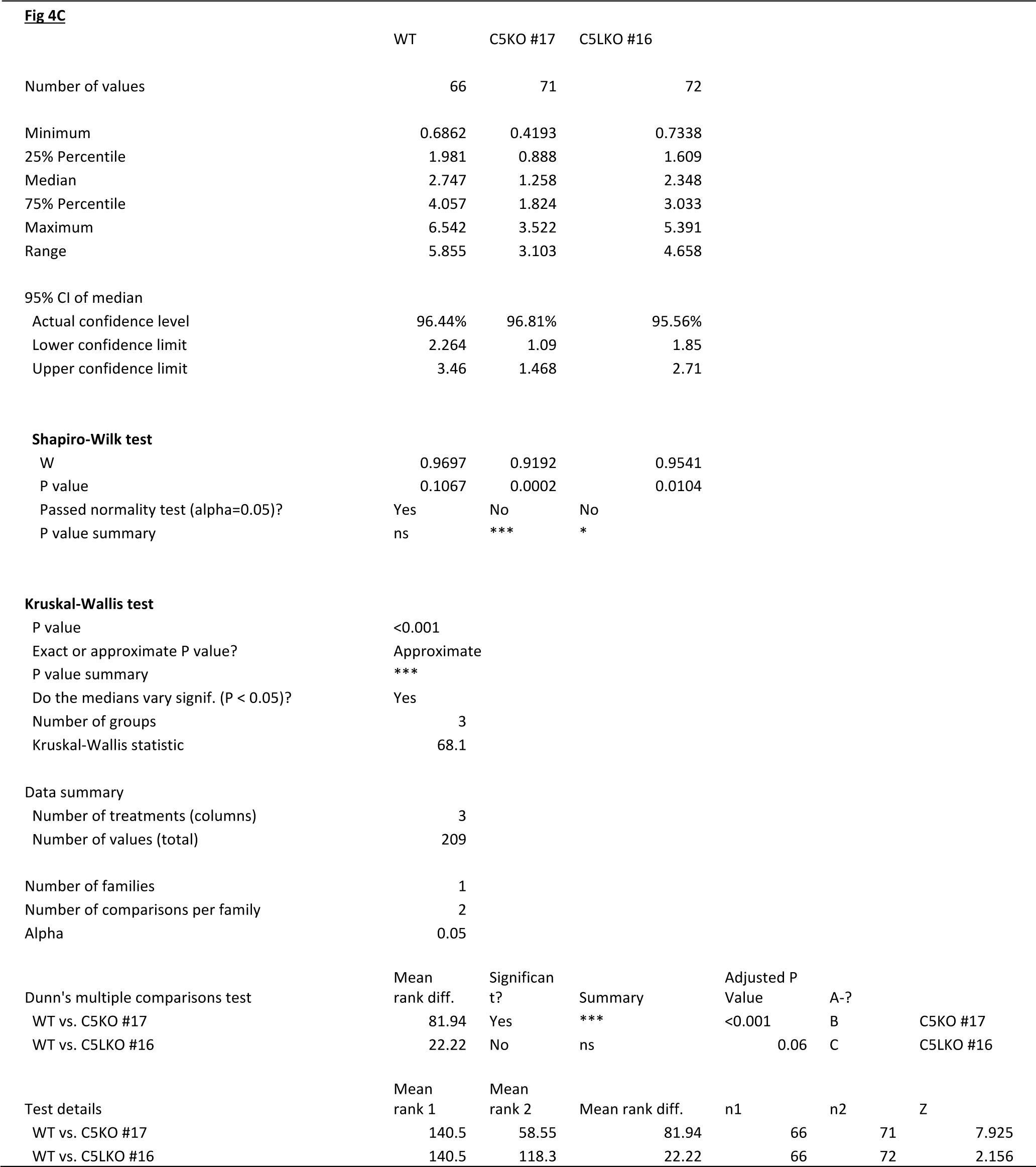
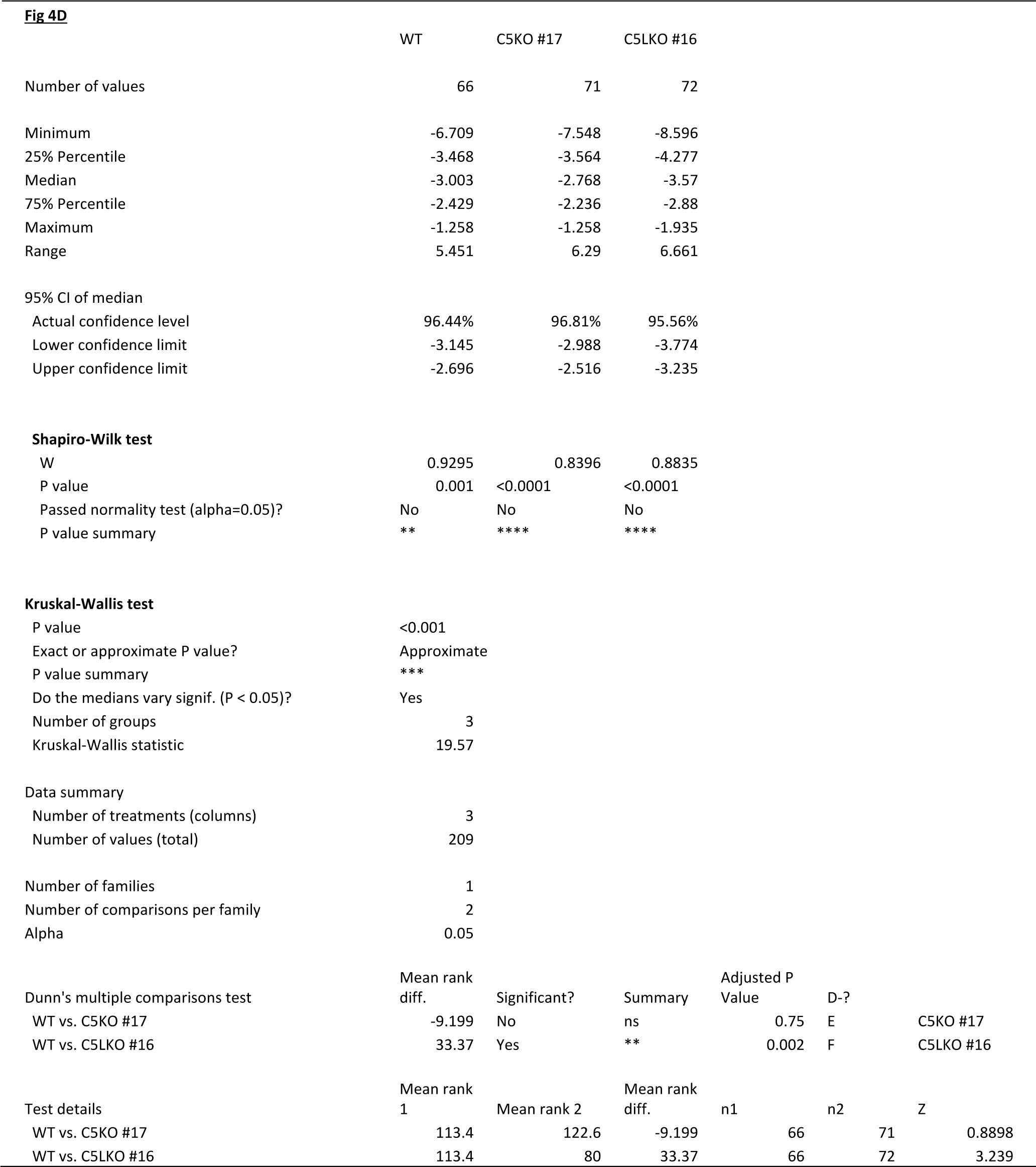
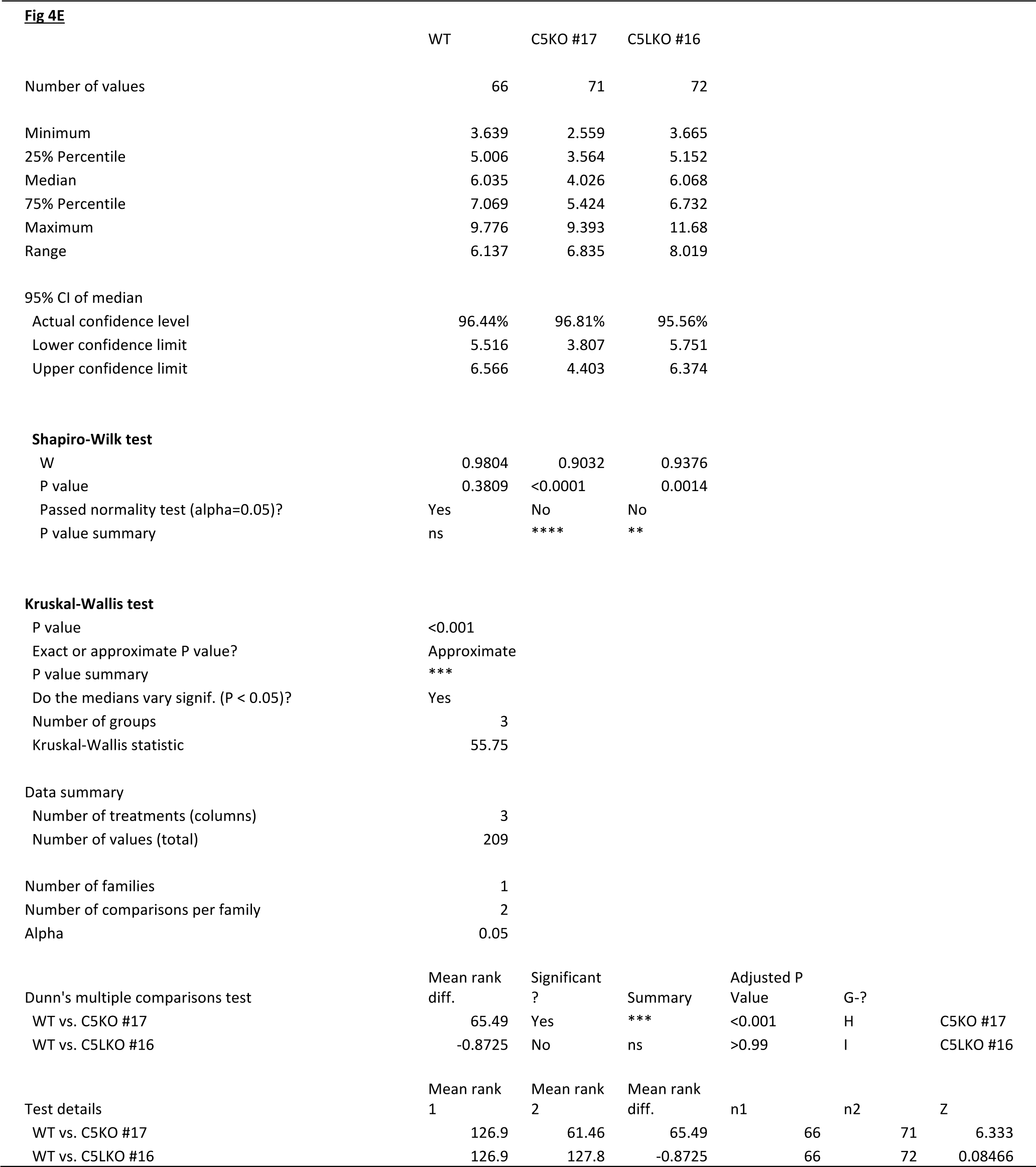
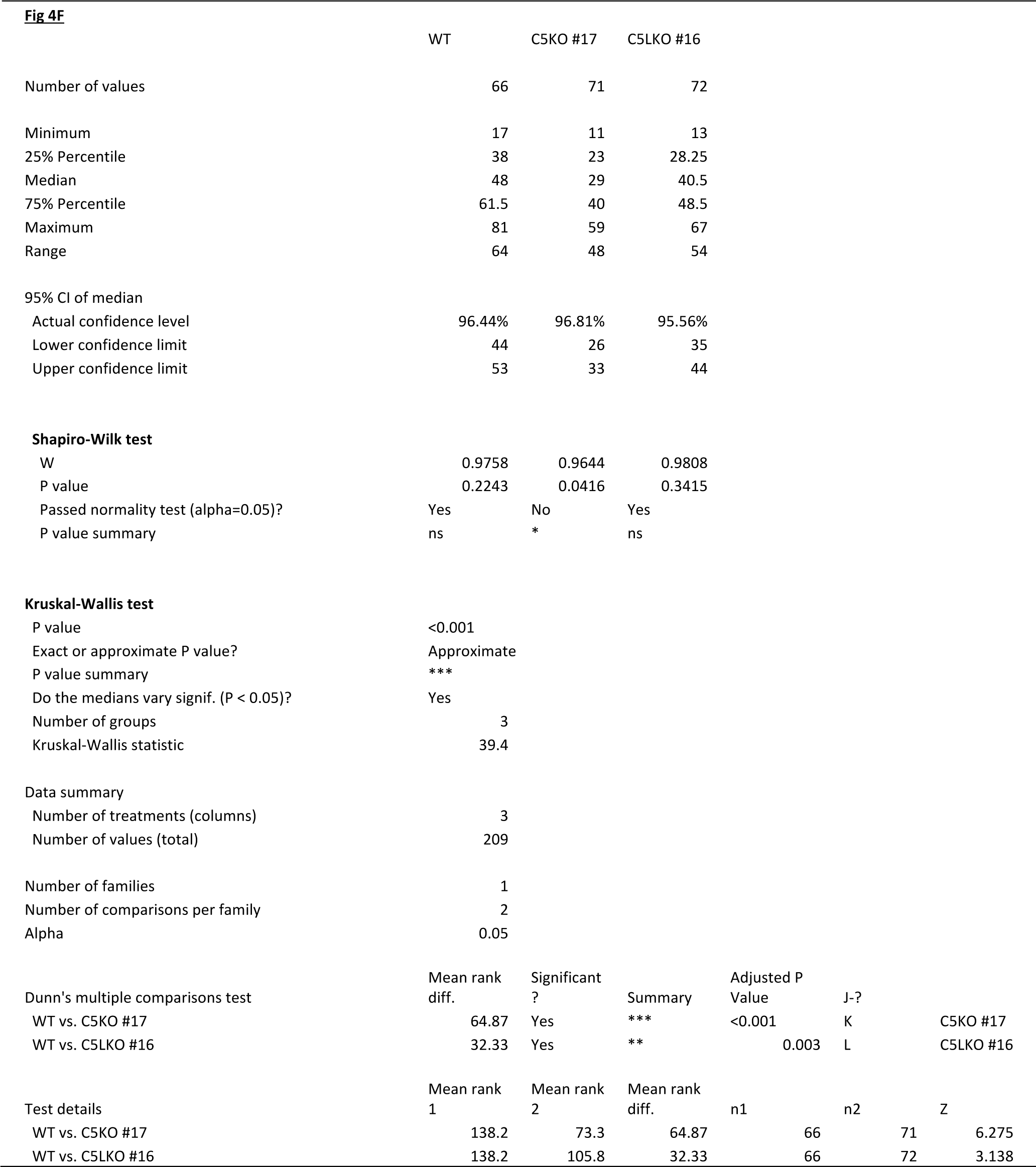
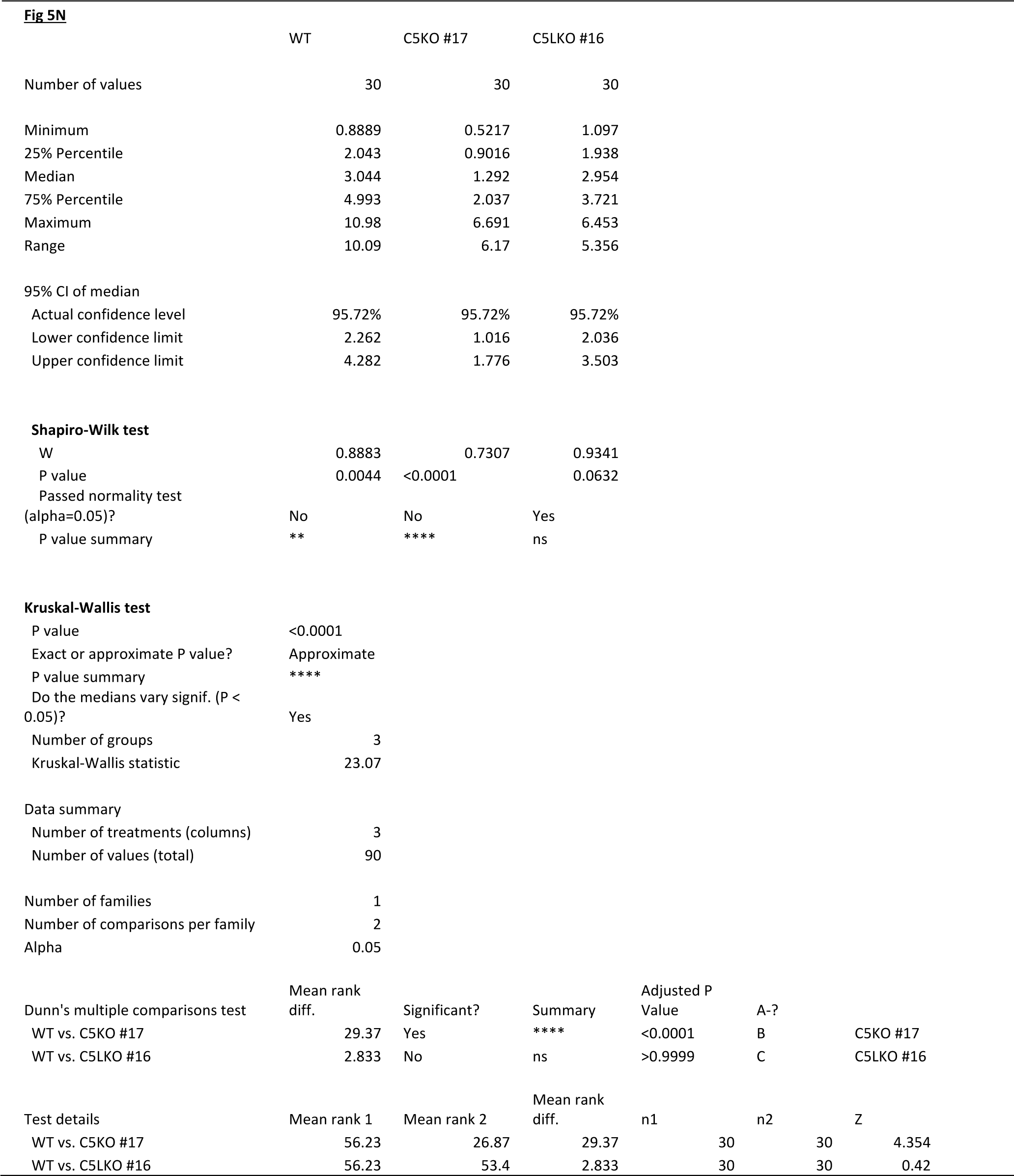
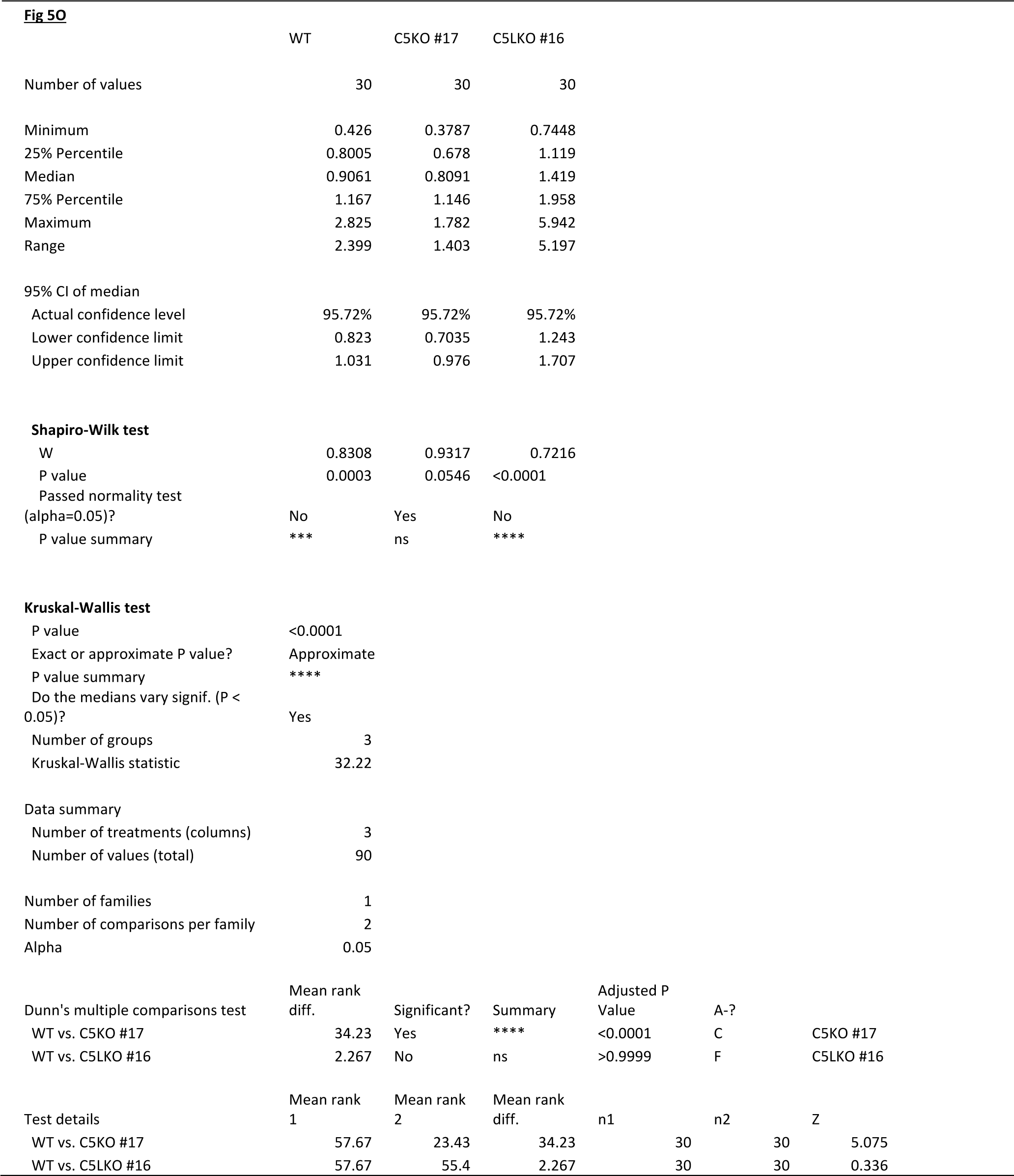
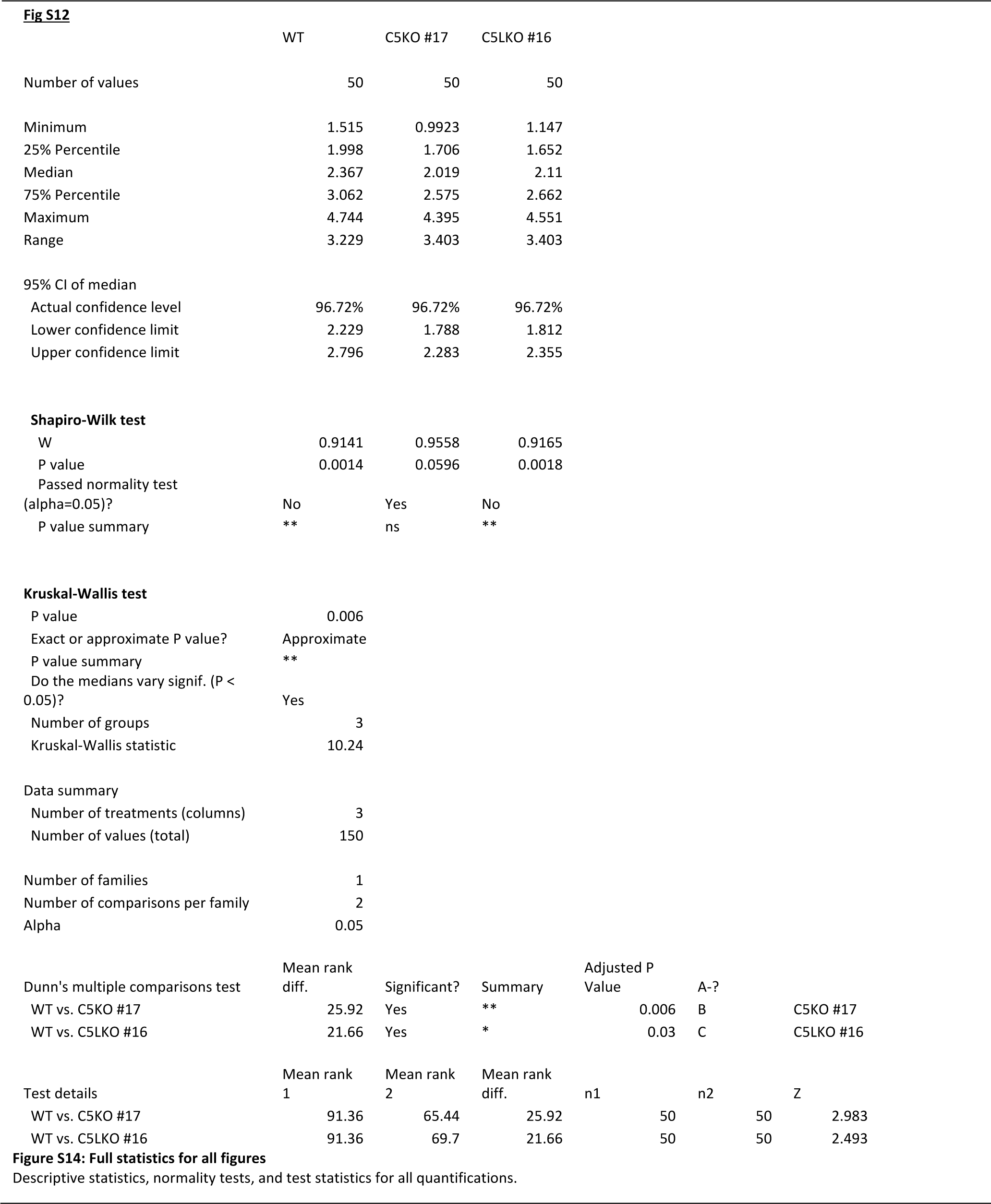

Video S1 Comparison of B16-F1 wildtype, C5KO, and C5LKO branch junction structures

Video S2 FRAP analysis of representative B16-F1 wildtype cell

Video S3 FRAP analysis of representative B16-F1 C5KO #17cell

Video S4 FRAP analysis of representative B16-F1 C5LKO #16 cell

## References

1. B. R. Schrank, T. Aparicio, Y. Li, W. Chang, B. T. Chait, G. G. Gundersen, M. E. Gottesman, J. Gautier, Nuclear ARP2/3 drives DNA break clustering for homology-directed repair. Nature. 559, 61–66 (2018).

2. A. M. Gautreau, F. E. Fregoso, G. Simanov, R. Dominguez, Nucleation, stabilization, and disassembly of branched actin networks. Trends in Cell Biology (2021), doi:10.1016/j.tcb.2021.10.006.

3. V. Papalazarou, L. M. Machesky, The cell pushes back: The Arp2/3 complex is a key orchestrator of cellular responses to environmental forces. Current Opinion in Cell Biology. 68, 37–44 (2021).

4. F. Frischknecht, V. Moreau, S. Röttger, S. Gonfloni, I. Reckmann, G. Superti-Furga, M. Way, Actin-based motility of vaccinia virus mimics receptor tyrosine kinase signalling. Nature. 401, 926–929 (1999).

5. N. Molinie, A. Gautreau, The Arp2/3 Regulatory System and Its Deregulation in Cancer. Physiological Reviews. 98, 215–238 (2018).

6. J. M. Stevens, E. E. Galyov, M. P. Stevens, Actin-dependent movement of bacterial pathogens. Nat Rev Microbiol. 4, 91–101 (2006).

7. M. D. Welch, A. Iwamatsu, T. J. Mitchison, Actin polymerization is induced by Arp 2/3 protein complex at the surface of Listeria monocytogenes. Nature. 385, 265–269 (1997).

8. L. Blanchoin, K. J. Amann, H. N. Higgs, J. B. Marchand, D. A. Kaiser, T. D. Pollard, Direct observation of dendritic actin filament networks nucleated by Arp2/3 complex and WASP/Scar proteins. Nature. 404, 1007–1011 (2000).

9. R. D. Mullins, J. A. Heuser, T. D. Pollard, The interaction of Arp2/3 complex with actin: nucleation, high affinity pointed end capping, and formation of branching networks of filaments. Proc Natl Acad Sci U S A. 95, 6181–6186 (1998).

10. N. Volkmann, K. J. Amann, S. Stoilova-McPhie, C. Egile, D. C. Winter, L. Hazelwood, J. E. Heuser, R. Li, T. D. Pollard, D. Hanein, Structure of Arp2/3 Complex in Its Activated State and in Actin Filament Branch Junctions. Science. 293, 2456–2459 (2001).

11. M. D. Welch, A. H. DePace, S. Verma, A. Iwamatsu, T. J. Mitchison, The Human Arp2/3 Complex Is Composed of Evolutionarily Conserved Subunits and Is Localized to Cellular Regions of Dynamic Actin Filament Assembly. J Cell Biol. 138, 375–384 (1997).

12. B. Ding, H. Y. Narvaez-Ortiz, Y. Singh, G. M. Hocky, S. Chowdhury, B. J. Nolen, Structure of Arp2/3 complex at a branched actin filament junction resolved by single-particle cryo-electron microscopy. Proceedings of the National Academy of Sciences. 119, e2202723119 (2022).

13. F. Fäßler, G. Dimchev, V.-V. Hodirnau, W. Wan, F. K. M. Schur, Cryo-electron tomography structure of Arp2/3 complex in cells reveals new insights into the branch junction. Nat Commun. 11, 6437 (2020).

14. A. M. Gautreau, F. E. Fregoso, G. Simanov, R. Dominguez, Nucleation, stabilization, and disassembly of branched actin networks. Trends in Cell Biology (2021), doi:10.1016/j.tcb.2021.10.006.

15. I. Rouiller, X.-P. Xu, K. J. Amann, C. Egile, S. Nickell, D. Nicastro, R. Li, T. D. Pollard, N. Volkmann, D. Hanein, The structural basis of actin filament branching by the Arp2/3 complex. J Cell Biol. 180, 887–895 (2008).

16. M. Vinzenz, M. Nemethova, F. Schur, J. Mueller, A. Narita, E. Urban, C. Winkler, C. Schmeiser, S. A. Koestler, K. Rottner, G. P. Resch, Y. Maeda, J. V. Small, Actin branching in the initiation and maintenance of lamellipodia. Journal of Cell Science. 125, 2775–2785 (2012).

17. L. A. Helgeson, B. J. Nolen, Mechanism of synergistic activation of Arp2/3 complex by cortactin and N-WASP. eLife. 2, e00884 (2013).

18. I. Dang, R. Gorelik, C. Sousa-Blin, E. Derivery, C. Guérin, J. Linkner, M. Nemethova, J. G. Dumortier, F. A. Giger, T. A. Chipysheva, V. D. Ermilova, S. Vacher, V. Campanacci, I. Herrada, A.-G. Planson, S. Fetics, V. Henriot, V. David, K. Oguievetskaia, G. Lakisic, F. Pierre, A. Steffen, A. Boyreau, N. Peyriéras, K. Rottner, S. Zinn-Justin, J. Cherfils, I. Bièche, A. Y. Alexandrova, N. B. David, J. V. Small, J. Faix, L. Blanchoin, A. Gautreau, Inhibitory signalling to the Arp2/3 complex steers cell migration. Nature. 503, 281–284 (2013).

19. L. Cai, A. M. Makhov, D. A. Schafer, J. E. Bear, Coronin 1B Antagonizes Cortactin and Remodels Arp2/3- Containing Actin Branches in Lamellipodia. Cell. 134, 828–842 (2008).

20. C. A. Ydenberg, S. B. Padrick, M. O. Sweeney, M. Gandhi, O. Sokolova, B. L. Goode, GMF Severs Actin- Arp2/3 Complex Branch Junctions by a Cofilin-like Mechanism. Current Biology. 23, 1037–1045 (2013).

21. J. V. G. Abella, C. Galloni, J. Pernier, D. J. Barry, S. Kjær, M.-F. Carlier, M. Way, Isoform diversity in the Arp2/3 complex determines actin filament dynamics. Nat Cell Biol. 18, 76–86 (2016).

22. C. Galloni, D. Carra, J. V. G. Abella, S. Kjær, P. Singaravelu, D. J. Barry, N. Kogata, C. Guérin, L. Blanchoin, M. Way, MICAL2 enhances branched actin network disassembly by oxidizing Arp3B-containing Arp2/3 complexes. Journal of Cell Biology. 220, e202102043 (2021).

23. O. von Loeffelholz, A. Purkiss, L. Cao, S. Kjaer, N. Kogata, G. Romet-Lemonne, M. Way, C. A. Moores, Cryo-EM of human Arp2/3 complexes provides structural insights into actin nucleation modulation by ARPC5 isoforms. Biology Open. 9, bio054304 (2020).

24. K. Rottner, T. E. B. Stradal, How distinct Arp2/3 complex variants regulate actin filament assembly. Nat Cell Biol. 18, 1–3 (2016).

25. S. Dolati, F. Kage, J. Mueller, M. Müsken, M. Kirchner, G. Dittmar, M. Sixt, K. Rottner, M. Falcke, On the relation between filament density, force generation, and protrusion rate in mesenchymal cell motility. Molecular Biology of the Cell (2018), doi:10.1091/mbc.E18-02-0082.

26. K. Iwaya, K. Norio, K. Mukai, Coexpression of Arp2 and WAVE2 predicts poor outcome in invasive breast carcinoma. Mod Pathol. 20, 339–343 (2007).

27. K. Iwaya, K. Oikawa, S. Semba, B. Tsuchiya, Y. Mukai, T. Otsubo, T. Nagao, M. Izumi, M. Kuroda, H. Domoto, K. Mukai, Correlation between liver metastasis of the colocalization of actin-related protein 2 and 3 complex and WAVE2 in colorectal carcinoma. Cancer Science. 98, 992–999 (2007).

28. T. Otsubo, K. Iwaya, Y. Mukai, Y. Mizokami, H. Serizawa, T. Matsuoka, K. Mukai, Involvement of Arp2/3 complex in the process of colorectal carcinogenesis. Mod Pathol. 17, 461–467 (2004).

29. M. Abercrombie, J. E. M. Heaysman, S. M. Pegrum, The locomotion of fibroblasts in culture I. Movements of the leading edge. Experimental Cell Research. 59, 393–398 (1970).

30. M. Abercrombie, E. Joan, M. Heaysman, S. M. Pegrum, The locomotion of fibroblasts in culture: II. “Ruffling.” Experimental Cell Research. 60, 437–444 (1970).

31. K. Rottner, M. Schaks, Assembling actin filaments for protrusion. Current Opinion in Cell Biology. 56, 53– 63 (2019).

32. J. Damiano-Guercio, L. Kurzawa, J. Mueller, G. Dimchev, M. Schaks, M. Nemethova, T. Pokrant, S. Brühmann, J. Linkner, L. Blanchoin, M. Sixt, K. Rottner, J. Faix, Loss of Ena/VASP interferes with lamellipodium architecture, motility and integrin-dependent adhesion. eLife. 9, e55351 (2020).

33. Q. Tang, M. Schaks, N. Koundinya, C. Yang, L. W. Pollard, T. M. Svitkina, K. Rottner, B. L. Goode, WAVE1 and WAVE2 have distinct and overlapping roles in controlling actin assembly at the leading edge. MBoC. 31, 2168–2178 (2020).

34. G. Dimchev, B. Amiri, A. C. Humphries, M. Schaks, V. Dimchev, T. E. B. Stradal, J. Faix, M. Krause, M. Way, M. Falcke, K. Rottner, Lamellipodin tunes cell migration by stabilizing protrusions and promoting adhesion formation. J Cell Sci. 133, jcs239020 (2020).

35. F. Kage, M. Winterhoff, V. Dimchev, J. Mueller, T. Thalheim, A. Freise, S. Brühmann, J. Kollasser, J. Block, G. Dimchev, M. Geyer, H.-J. Schnittler, C. Brakebusch, T. E. B. Stradal, M.-F. Carlier, M. Sixt, J. Käs, J. Faix, K. Rottner, FMNL formins boost lamellipodial force generation. Nat Commun. 8, 14832 (2017).

36. M. Hakala, H. Wioland, M. Tolonen, T. Kotila, A. Jegou, G. Romet-Lemonne, P. Lappalainen, Twinfilin uncaps filament barbed ends to promote turnover of lamellipodial actin networks. Nat Cell Biol. 23, 147– 159 (2021).

37. K. L. Anderson, C. Page, M. F. Swift, P. Suraneni, M. E. W. Janssen, T. D. Pollard, R. Li, N. Volkmann, D. Hanein, Nano-scale actin-network characterization of fibroblast cells lacking functional Arp2/3 complex. Journal of Structural Biology. 197, 312–321 (2017).

38. J. Mueller, G. Szep, M. Nemethova, I. de Vries, A. D. Lieber, C. Winkler, K. Kruse, J. V. Small, C. Schmeiser, K. Keren, R. Hauschild, M. Sixt, Load Adaptation of Lamellipodial Actin Networks. Cell. 171, 188–200.e16 (2017).

39. J. V. Small, M. Herzog, K. Anderson, Actin filament organization in the fish keratocyte lamellipodium. Journal of Cell Biology. 129, 1275–1286 (1995).

40. T. M. Svitkina, G. G. Borisy, Arp2/3 Complex and Actin Depolymerizing Factor/Cofilin in Dendritic Organization and Treadmilling of Actin Filament Array in Lamellipodia. Journal of Cell Biology. 145, 1009– 1026 (1999).

41. T. Kinoshita, N. Nohata, H. Watanabe-Takano, H. Yoshino, H. Hidaka, L. Fujimura, M. Fuse, T. Yamasaki, H. Enokida, M. Nakagawa, T. Hanazawa, Y. Okamoto, N. Seki, Actin-related protein 2/3 complex subunit 5 (ARPC5) contributes to cell migration and invasion and is directly regulated by tumor-suppressive microRNA-133a in head and neck squamous cell carcinoma. International Journal of Oncology. 40, 1770– 1778 (2012).

42. T. Xiong, Z. Luo, The Expression of Actin-Related Protein 2/3 Complex Subunit 5 (ARPC5) Expression in Multiple Myeloma and its Prognostic Significance. Med Sci Monit. 24, 6340–6348 (2018).

43. S. Huang, D. Li, L. Zhuang, L. Sun, J. Wu, Identification of Arp2/3 Complex Subunits as Prognostic Biomarkers for Hepatocellular Carcinoma. Front Mol Biosci. 8, 690151 (2021).

44. F. A. Ran, P. D. Hsu, J. Wright, V. Agarwala, D. A. Scott, F. Zhang, Genome engineering using the CRISPR- Cas9 system. Nat Protoc. 8, 2281–2308 (2013).

45. H. Gournier, E. D. Goley, H. Niederstrasser, T. Trinh, M. D. Welch, Reconstitution of Human Arp2/3 Complex Reveals Critical Roles of Individual Subunits in Complex Structure and Activity. Molecular Cell. 8, 1041–1052 (2001).

46. K. Rottner, J. Faix, S. Bogdan, S. Linder, E. Kerkhoff, Actin assembly mechanisms at a glance. Journal of Cell Science. 130, 3427–3435 (2017).

47. F. Kage, H. Döring, M. Mietkowska, M. Schaks, F. Grüner, S. Stahnke, A. Steffen, M. Müsken, T. E. B. Stradal, K. Rottner, Lamellipodia-like actin networks in cells lacking WAVE Regulatory Complex (2021), p. 2021.06.18.449030, , doi:10.1101/2021.06.18.449030.

48. T. Uruno, J. Liu, P. Zhang, Y. Fan, C. Egile, R. Li, S. C. Mueller, X. Zhan, Activation of Arp2/3 complex-mediated actin polymerization by cortactin. Nat Cell Biol. 3, 259–266 (2001).

49. A. M. Weaver, A. V. Karginov, A. W. Kinley, S. A. Weed, Y. Li, J. T. Parsons, J. A. Cooper, Cortactin promotes and stabilizes Arp2/3-induced actin filament network formation. Current Biology. 11, 370–374 (2001).

50. F. P. L. Lai, M. Szczodrak, J. M. Oelkers, M. Ladwein, F. Acconcia, S. Benesch, S. Auinger, J. Faix, J. V. Small, S. Polo, T. E. B. Stradal, K. Rottner, Cortactin Promotes Migration and Platelet-derived Growth Factor-induced Actin Reorganization by Signaling to Rho-GTPases. MBoC. 20, 3209–3223 (2009).

51. F. Fäßler, B. Zens, R. Hauschild, F. K. M. Schur, 3D printed cell culture grid holders for improved cellular specimen preparation in cryo-electron microscopy. Journal of Structural Biology. 212, 107633 (2020).

52. G. Dimchev, B. Amiri, F. Fäßler, M. Falcke, F. K. Schur, Computational toolbox for ultrastructural quantitative analysis of filament networks in cryo-ET data. Journal of Structural Biology. 213, 107808 (2021).

53. J. Faix, K. Rottner, Ena/VASP proteins in cell edge protrusion, migration and adhesion. Journal of Cell Science. 135, jcs259226 (2022).

54. W. Wang, S. Goswami, K. Lapidus, A. L. Wells, J. B. Wyckoff, E. Sahai, R. H. Singer, J. E. Segall, J. S. Condeelis, Identification and Testing of a Gene Expression Signature of Invasive Carcinoma Cells within Primary Mammary Tumors. Cancer Research. 64, 8585–8594 (2004).

55. U. Philippar, E. Sahai, J. Wyckoff, H. Yamaguchi, M. Oser, S. Giampieri, Y. Wang, J. Condeelis, F. Gertler, Ena/VASP proteins promote cancer cell invasion. Cancer Research. 67, 559A (2007).

56. L.-D. Hu, H.-F. Zou, S.-X. Zhan, K.-M. Cao, EVL (Ena/VASP-like) expression is up-regulated in human breast cancer and its relative expression level is correlated with clinical stages. Oncology Reports. 19, 1015– 1020 (2008).

57. M. Barone, M. Müller, S. Chiha, J. Ren, D. Albat, A. Soicke, S. Dohmen, M. Klein, J. Bruns, M. van Dinther, R. Opitz, P. Lindemann, M. Beerbaum, K. Motzny, Y. Roske, P. Schmieder, R. Volkmer, M. Nazaré, U. Heinemann, H. Oschkinat, P. ten Dijke, H.-G. Schmalz, R. Kühne, Designed nanomolar small-molecule inhibitors of Ena/VASP EVH1 interaction impair invasion and extravasation of breast cancer cells. Proceedings of the National Academy of Sciences. 117, 29684–29690 (2020).

58. S. Havrylenko, P. Noguera, M. Abou-Ghali, J. Manzi, F. Faqir, A. Lamora, C. Guérin, L. Blanchoin, J. Plastino, WAVE binds Ena/VASP for enhanced Arp2/3 complex–based actin assembly. MBoC. 26, 55–65 (2015).

59. M. Shaaban, S. Chowdhury, B. J. Nolen, Cryo-EM reveals the transition of Arp2/3 complex from inactive to nucleation-competent state. Nat Struct Mol Biol. 27, 1009–1016 (2020).

60. Q. Luan, B. J. Nolen, Structural basis for regulation of Arp2/3 complex by GMF. Nat Struct Mol Biol. 20, 1062–1068 (2013).

61. L. Sadhu, N. Tsopoulidis, V. Laketa, M. Way, O. T. Fackler, ARPC5 Isoforms Drive Distinct Arp2/3- dependant Actin Remodeling Events in CD4 T Cells (2022), p. 2022.01.25.477674, , doi:10.1101/2022.01.25.477674.

62. S. Asano, B. D. Engel, W. Baumeister, In Situ Cryo-Electron Tomography: A Post-Reductionist Approach to Structural Biology. J Mol Biol. 428, 332–343 (2016).

63. L. Aldaz-Carroll, J. C. Whitbeck, M. Ponce de Leon, H. Lou, L. Hirao, S. N. Isaacs, B. Moss, R. J. Eisenberg, G. H. Cohen, Epitope-Mapping Studies Define Two Major Neutralization Sites on the Vaccinia Virus Extracellular Enveloped Virus Glycoprotein B5R. Journal of Virology. 79, 6260–6271 (2005).

64. L. Aldaz-Carroll, J. C. Whitbeck, M. P. de Leon, H. Lou, L. K. Pannell, J. Lebowitz, C. Fogg, C. L. White, B. Moss, G. H. Cohen, R. J. Eisenberg, Physical and immunological characterization of a recombinant secreted form of the membrane protein encoded by the vaccinia virus L1R gene. Virology. 341, 59–71 (2005).

65. F. Künzl, S. Früholz, F. Fäßler, B. Li, P. Pimpl, Receptor-mediated sorting of soluble vacuolar proteins ends at the trans-Golgi network/early endosome. Nature Plants. 2, 1–10 (2016).

66. J. Schindelin, I. Arganda-Carreras, E. Frise, V. Kaynig, M. Longair, T. Pietzsch, S. Preibisch, C. Rueden, S. Saalfeld, B. Schmid, J.-Y. Tinevez, D. J. White, V. Hartenstein, K. Eliceiri, P. Tomancak, A. Cardona, Fiji: an open-source platform for biological-image analysis. Nat Methods. 9, 676–682 (2012).

67. D. N. Mastronarde, Automated electron microscope tomography using robust prediction of specimen movements. J Struct Biol. 152, 36–51 (2005).

68. W. J. H. Hagen, W. Wan, J. A. G. Briggs, Implementation of a cryo-electron tomography tilt-scheme optimized for high resolution subtomogram averaging. Journal of Structural Biology. 197, 191–198 (2017).

69. D. Tegunov, P. Cramer, Real-time cryo-electron microscopy data preprocessing with Warp. Nat Methods. 16, 1146–1152 (2019).

70. J. R. Kremer, D. N. Mastronarde, J. R. McIntosh, Computer visualization of three-dimensional image data using IMOD. J Struct Biol. 116, 71–76 (1996).

71. D. Castaño-Díez, M. Kudryashev, M. Arheit, H. Stahlberg, Dynamo: a flexible, user-friendly development tool for subtomogram averaging of cryo-EM data in high-performance computing environments. J Struct Biol. 178, 139–151 (2012).

72. S. H. W. Scheres, RELION: Implementation of a Bayesian approach to cryo-EM structure determination. J Struct Biol. 180, 519–530 (2012).

73. J. Zivanov, T. Nakane, B. O. Forsberg, D. Kimanius, W. J. Hagen, E. Lindahl, S. H. Scheres, New tools for automated high-resolution cryo-EM structure determination in RELION-3. eLife. 7, e42166 (2018).

74. S. Pospich, F. Merino, S. Raunser, Structural Effects and Functional Implications of Phalloidin and Jasplakinolide Binding to Actin Filaments. Structure. 28, 437–449.e5 (2020).

75. A. Burt, L. Gaifas, T. Dendooven, I. Gutsche, A flexible framework for multi-particle refinement in cryo-electron tomography. PLOS Biology. 19, e3001319 (2021).

76. D. Tegunov, L. Xue, C. Dienemann, P. Cramer, J. Mahamid, Multi-particle cryo-EM refinement with M visualizes ribosome-antibiotic complex at 3.5 Å in cells. Nat Methods. 18, 186–193 (2021).

77. A. P. Joseph, I. Lagerstedt, A. Jakobi, T. Burnley, A. Patwardhan, M. Topf, M. Winn, Comparing Cryo-EM Reconstructions and Validating Atomic Model Fit Using Difference Maps. J. Chem. Inf. Model. 60, 2552– 2560 (2020).

78. C. Wood, T. Burnley, A. Patwardhan, S. Scheres, M. Topf, A. Roseman, M. Winn, Collaborative Computational Project for Electron cryo-Microscopy. Acta Cryst D. 71, 123–126 (2015).

79. T. Burnley, C. M. Palmer, M. Winn, Recent developments in the CCP-EM software suite. Acta Cryst D. 73, 469–477 (2017).

